# Language model-guided anticipation and discovery of unknown metabolites

**DOI:** 10.1101/2024.11.13.623458

**Authors:** Hantao Qiang, Fei Wang, Wenyun Lu, Xi Xing, Hahn Kim, Sandrine A.M. Merette, Lucas B. Ayres, Eponine Oler, Jenna E. AbuSalim, Asael Roichman, Michael Neinast, Ricardo A. Cordova, Won Dong Lee, Ehud Herbst, Vishu Gupta, Samuel Neff, Mickel Hiebert-Giesbrecht, Adamo Young, Vasuk Gautam, Siyang Tian, Bo Wang, Hannes Röst, Russell Greiner, Li Chen, Chad W. Johnston, Leonard J. Foster, Aaron M. Shapiro, David S. Wishart, Joshua D. Rabinowitz, Michael A. Skinnider

**Author notes:** These authors contributed equally (Hantao Qiang, Fei Wang). These authors jointly supervised this work (David S. Wishart, Joshua D. Rabinowitz, Michael A. Skinnider).

## Abstract

Despite decades of study, large parts of the mammalian metabolome remain unexplored. Mass spectrometry-based metabolomics routinely detects thousands of small molecule-associated peaks within human tissues and biofluids, but typically only a small fraction of these can be identified, and structure elucidation of novel metabolites remains a low-throughput endeavor. Biochemical large language models have transformed the interpretation of DNA, RNA, and protein sequences, but have not yet had a comparable impact on understanding small molecule metabolism. Here, we present an approach that leverages chemical language models to discover previously uncharacterized metabolites. We introduce DeepMet, a chemical language model that learns the latent biosynthetic logic embedded within the structures of known metabolites and exploits this understanding to anticipate the existence of as-of-yet undiscovered metabolites. Prospective chemical synthesis of metabolites predicted to exist by DeepMet directs their targeted discovery. Integrating DeepMet with tandem mass spectrometry (MS/MS) data enables automated metabolite discovery within complex tissues. We harness DeepMet to discover several dozen structurally diverse mammalian metabolites. Our work demonstrates the potential for language models to accelerate the mapping of the metabolome.

---

High-throughput techniques now enable routine measurement of the DNA, RNA, and protein content of any given biospecimen. In contrast, enumerating the complete complement of small molecules—the metabolome—has proven more challenging. Mass spectrometry-based metabolomics typically detects thousands of distinct chemical entities in any given biological sample^1^. However, even in human tissues or biofluids, the majority of mass spectrometric signals cannot routinely be linked to a known chemical structure^2,3^. This profusion of unidentified chemical entities, which collectively account for most of the information acquired in a typical metabolomic study, have been dubbed the chemical “dark matter” of the metabolome^4^.

The existence of this metabolic dark matter suggests that existing metabolic maps are far from complete. Recent discoveries have described important roles for previously unidentified metabolites in human physiology and disease^5–9^, implying that a more comprehensive map of metabolism could accelerate the discovery of new biological mechanisms, biomarkers, and therapeutic targets. However, structure elucidation of unknown metabolites remains a fundamentally low-throughput endeavour. Isolation of purified compounds followed by detailed structural characterization is time- and labour-intensive, restricting such efforts to at most a handful of carefully selected metabolites^10^. New approaches are needed to illuminate the dark matter of the metabolome in a systematic and high-throughput manner.

Generative models based on deep neural networks have emerged as a powerful approach to study the structure and function of biological macromolecules^11^. Language models trained on protein sequences are capable of learning the latent evolutionary forces that have shaped observed sequences in order to design new proteins with desired functions, predict the impacts of unseen variants, and even forecast protein sequences likely to evolve in the future^12–16^. Language models can also be trained on the chemical structures of small molecules by leveraging formats that represent these structures as short strings of text^17–19^. To date, however, this paradigm has primarily been applied to explore synthetic chemical space in the setting of drug discovery. We hypothesized that, in analogy to protein language models, a language model of metabolism could learn the biosynthetic logic that has sculpted the structures of known metabolites, and exploit this understanding to predict the structures of metabolites that have yet to be discovered.

Here, we present an approach that leverages language models to systematically fill the gaps in existing maps of the metabolome (**Fig. 1a**). We introduce DeepMet, a chemical language model trained on the structures of known metabolites. We demonstrate that DeepMet systematically anticipates the existence of as-of-yet undiscovered metabolites, and validate dozens of these predictions experimentally. We develop approaches to integrate DeepMet with mass spectrometry-based metabolomics data that enable *de novo* identification of novel metabolites in complex tissues. In total, we harness DeepMet to discover 47 structurally diverse metabolites.

**Fig. 1.**
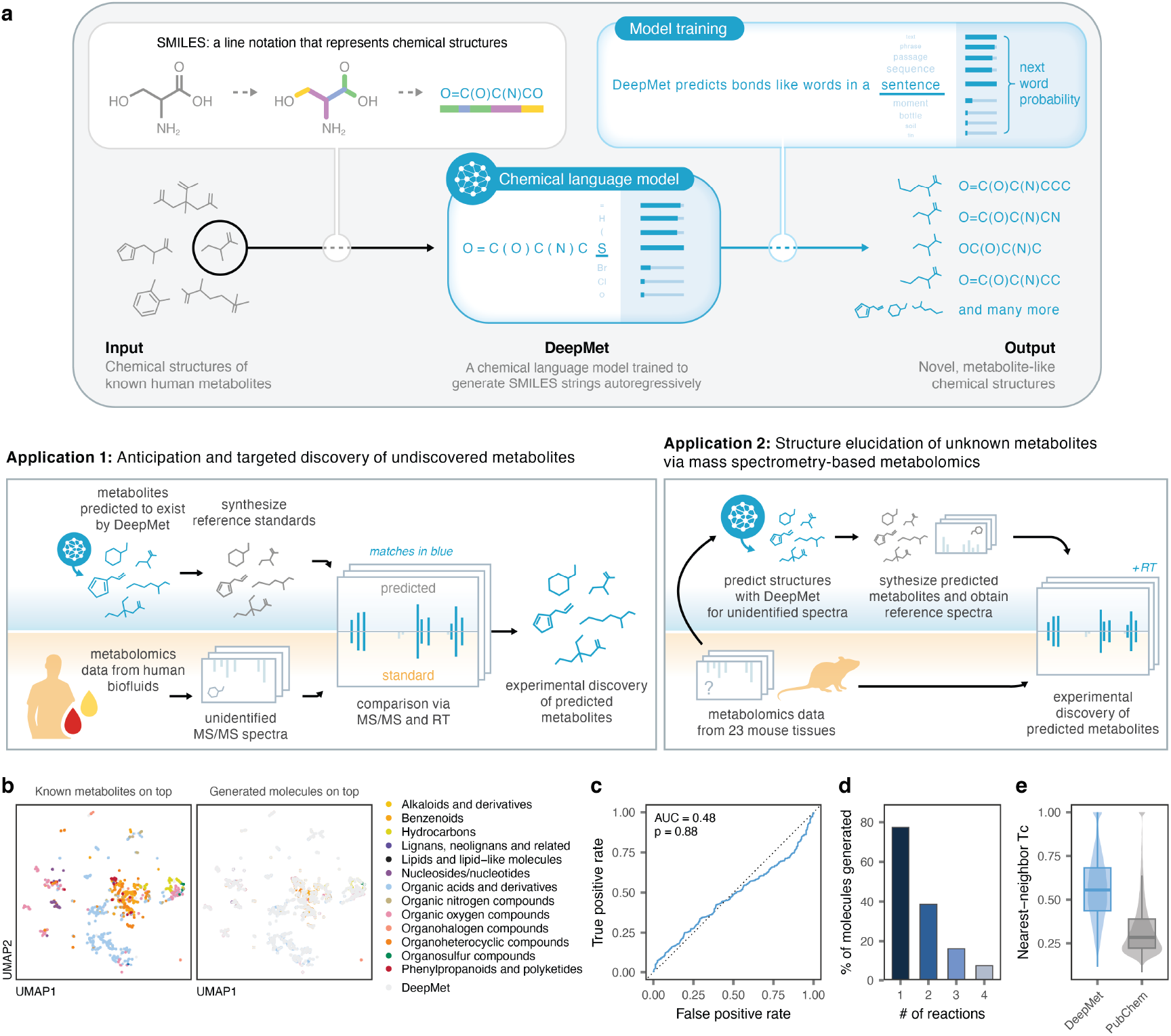
Learning the language of metabolism. **a**, Schematic overview of DeepMet, a chemical language model for the systematic prioritization and discovery of unknown metabolites. **b**, UMAP visualization of the chemical space occupied by known metabolites and generated molecules. Left, known metabolites superimposed over generated molecules; right, vice versa. Metabolites are colored by their assigned superclasses in the ClassyFire chemical ontology. **c**, Receiver operating characteristic (ROC) curve of a random forest classifier trained to distinguish between known metabolites and generated molecules in cross-validation. Inset text shows the area under the ROC curve (AUC) and its statistical significance. **d**, Proportion of enzymatic biotransformations of known metabolites recapitulated by DeepMet, shown as a function of the number of rule-based transformations applied sequentially to the original metabolite. **e**, Nearest-neighbor Tanimoto coefficient to a known metabolite for generated molecules versus molecules with identical molecular formulas sampled randomly from PubChem.

## Results

### Learning the language of metabolism

Metabolites are synthesized from a small pool of precursors, such as amino acids, organic acids, sugars, and acetylCoA, via a limited repertoire of enzymatic transformations. These shared biosynthetic origins result in the overrepresentation of certain physicochemical properties and substructures among metabolites, as compared to synthetic compounds made in the laboratory^20–23^. We hypothesized that a chemical language model could learn the biosynthetic logic embedded within the structures of known metabolites, and then leverage this understanding to access novel structures from metabolite-like chemical space.

To test this hypothesis, we assembled a training set of 2,046 metabolites that had been experimentally detected in human tissues or biofluids^24^ and represented these as short strings of text in simplified molecular-input line-entry system (SMILES) notation^25^. We trained a language model on this dataset of known metabolite structures, and used the trained model to generate 500,000 SMILES strings in order to evaluate its understanding of metabolism.

Several lines of evidence indicated that our language model was able to appreciate the structural features of known metabolites and exploit this understanding to generate novel, metabolite-like structures. First, we visualized the areas of chemical space occupied by generated molecules and known metabolites using the nonlinear dimensionality reduction algorithm UMAP^26^. Generated molecules overlapped extensively with known metabolites (**Fig. 1b**). Second, we trained a random forest classifier to distinguish the generated molecules from a set of known human metabolites that had been deliberately withheld from the language model during training. We found that this classifier was unable to separate the two classes of molecules to a degree better than random guessing (area under the receiver operating characteristic curve [AUC] = 0.48; **Fig. 1c** and **Supplementary Fig. 1a-b)**, indicating that the generated molecules were structurally indistinguishable from known metabolites. Third, because many biosynthetic enzymes are known to be promiscuous in the substrates they accept, we asked whether the generated molecules could be rationalized as enzymatic transformations of known metabolites. In line with this hypothesis, we found that the language model recapitulated 77.5% of one-step enzymatic transformations of known metabolites predicted by the rule-based platform BioTransformer^27^ (**Fig. 1d** and Supplementary **Fig. 1c-d**). Fourth, we quantified the structural similarity between each generated molecule and its nearest neighbor in the training set using the Tanimoto coefficient^28^, and found that the generated molecules exhibited a markedly higher degree of structural similarity to known metabolites than molecules with identical molecular formulas sampled at random from PubChem (**Fig. 1e** and **Supplementary Fig. 1e**).

These results introduce a language model of metabolite-like chemical space, which we named DeepMet.

### Anticipation of undiscovered metabolites

Language models are trained to identify parameters that maximize the likelihood of the sequences in the training dataset. In the setting of protein biochemistry, the likelihoods assigned to unobserved amino acid sequences by language models can be leveraged to predict the functional impact of unseen mutations and even forecast the evolution of future proteins^12–15^. We hypothesized that the same principle could be applied to predict the structures of as-of-yet undiscovered metabolites. Unlike nucleotide or protein sequences, however, chemical structures do not have a unique textual representation^29^, and we observed that DeepMet frequently assigned markedly different likelihoods to different SMILES strings representing the same chemical structure (**Supplementary Fig. 2a-c**).

In lieu of calculating the likelihood of individual SMILES strings, we reasoned that novel chemical structures viewed by DeepMet as more plausible extensions of the training set would be sampled more frequently in aggregate, considering all possible representations. To test this hypothesis, we drew a sample of 1 billion SMILES strings from DeepMet, and then tabulated the frequency with which each unique chemical structure appeared within this sample (**Fig. 2a**). Whereas the vast majority of these structures appeared at most a handful of times within the language model output, certain previously unreported structures were generated thousands of times (**Fig. 2b**).

**Fig. 2.**
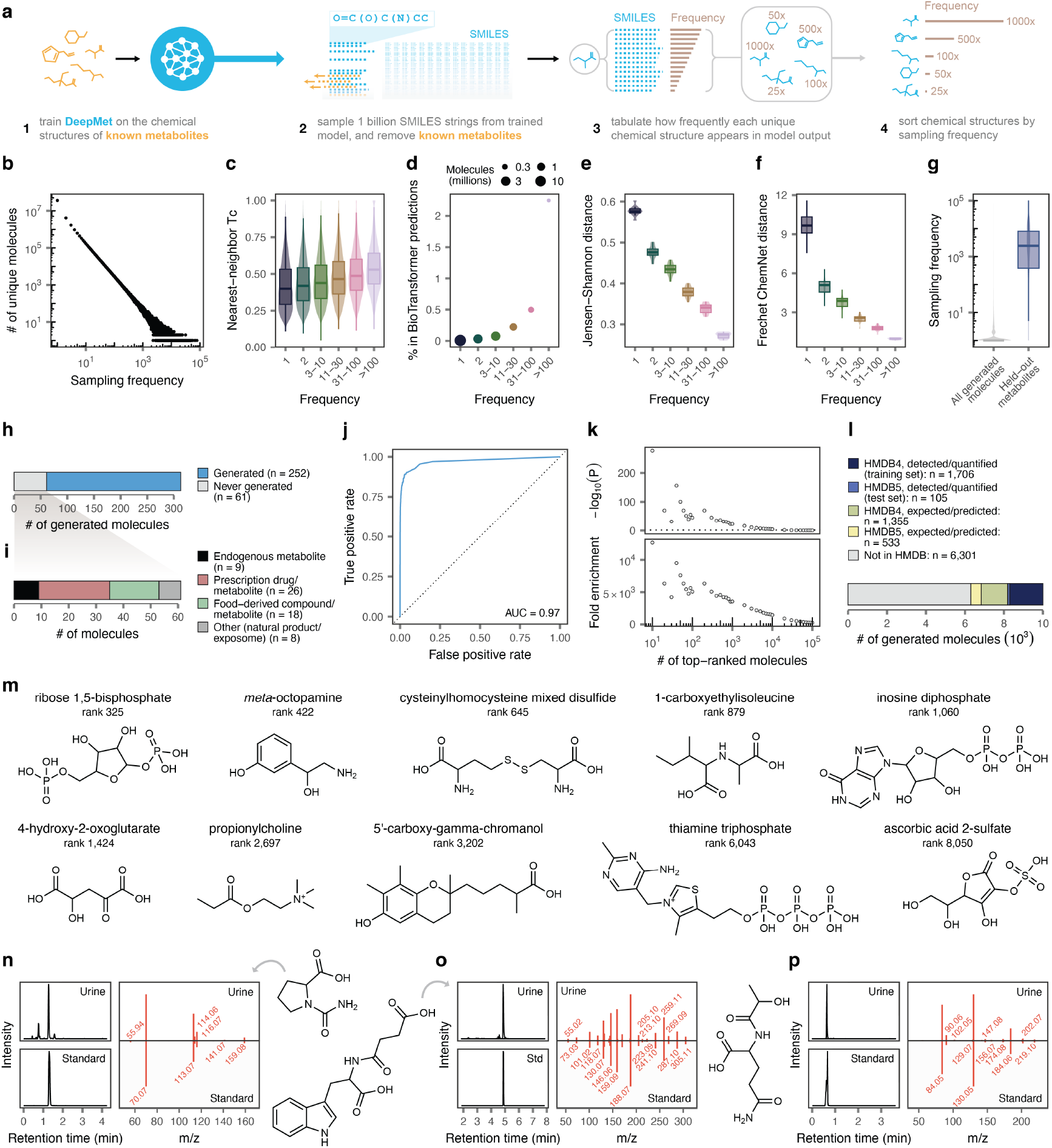
Anticipation and language model-guided discovery of human metabolites. **a**, Schematic overview of the workflow to calculate sampling frequencies for generated molecules. **b**, Distribution of sampling frequencies within a sample of 1 billion SMILES strings from DeepMet. **c**, Tanimoto coefficient between generated molecules and their nearest neighbour in the training set of known metabolites, for molecules generated with progressively increasing frequencies. **d**, Proportion of generated metabolites recapitulating one-step enzymatic transformations of human metabolites predicted by BioTransformer, for molecules generated with progressively increasing frequencies. **e**, Jensen–Shannon distance between the Murcko scaffold compositions of generated molecules and the training set of known metabolites, for molecules generated with progressively increasing frequencies. **f**, Fréchet ChemNet distances between generated molecules and the training set of known metabolites, for molecules generated with progressively increasing frequencies. **g**, Frequencies with which known metabolites withheld from the training set were sampled, as compared to the background distribution of all generated molecules. **h**, Proportion of human metabolites added in version 5 of the HMDB that were generated by DeepMet. **i**, Categorization of the 61 HMDB version 5 metabolites that were not successfully generated by DeepMet. **j**, ROC curve showing the prioritization of human metabolites added in version 5 of the HMDB on the basis of sampling frequency, as compared to the background distribution of all generated molecules. **k**, Enrichment of newly-discovered human metabolites among the most frequently generated molecules. Bottom, fold enrichment within the top-10 to top-10^5^ most frequently generated molecules; top, statistical significance of the enrichment (*χ*^2^ test). **l**, Proportion of known or ‘predicted/expected’ metabolites from versions 4 or 5 of the HMDB within the top-10,000 molecules most frequently generated by DeepMet. **m**, Examples of metabolites annotated as ‘predicted’ or ‘expected’ in HMDB that are actually well-studied human metabolites, and which were generated with frequencies comparable to experimentally detected metabolites by DeepMet despite being withheld entirely from the training set. **n-p**, Examples of novel human metabolites predicted to exist by DeepMet and experimentally discovered in human urine (chemical structures of the predicted metabolites; extracted ion chromatograms from the chemical standard and a representative urine metabolome; and mirror plots showing the similarity between MS/MS spectra from the standard versus the experimental spectrum from human urine). **n**, N-carbamylproline; **o**, N-succinyl-tryptophan; **p**, N-lactoyl-glutamine.

We sought to characterize the properties of these frequently generated molecules. Molecules sampled more frequently by DeepMet exhibited a higher degree of structural similarity to known metabolites (**Fig. 2c** and **Supplementary Fig. 2d**); were disproportionately likely to overlap with plausible enzymatic transformations^27^ of known metabolites (**Fig. 2d** and **Supplementary Fig. 2e**); were more likely to share a chemical scaffold with a known metabolite^30^ (Fig. 2e); and, as quantified by the Fréchet ChemNet distance^31^, were predicted to have a more similar spectrum of biological activities to known metabolites (**Fig. 2f**). Thus, molecules generated more frequently by DeepMet were disproportionately metabolite-like.

This finding led us to more directly test whether this sampling frequency could be used to prioritize candidate metabolites for discovery. To evaluate this possibility, we trained DeepMet in cross-validation, withholding 10% of the known metabolites in the training set at a time in order to simulate the discovery of unknown metabolites. The with-held metabolites were generally among the most frequently generated molecules proposed by the language model (**Fig. 2g**), such that the sampling frequency alone separated with-held metabolites from other generated molecules with a near-perfect AUC of 0.98 (**Supplementary Fig. 2f**).

Encouraged by this finding, we sought to prospectively evaluate the ability of DeepMet to predict the structures of as-of-yet undiscovered metabolites. We compiled a prospective test set of 313 metabolites that had been added to the Human Metabolome Database (HMDB) after our training dataset was finalized^32^. DeepMet successfully generated 252 of these 313 newly-discovered metabolites (81%; **Fig. 2h**). Among the 61 structures that were not successfully generated, most were not endogenous human metabolites but instead included prescription drugs, food-derived compounds, bacterial or fungal natural products, or synthetic compounds from the human ‘exposome’ (**Fig. 2i** and **Supplementary Fig. 2g**). Only 9 of the 313 structures that were *bona fide* human metabolites were not successfully generated by DeepMet (**Supplementary Fig. 2h**). Moreover, we again observed that the sampling frequency alone accurately separated these newly-discovered metabolites from other generated molecules (AUC = 0.97; **Fig. 2j**).

Newly-discovered metabolites were markedly enriched within the uppermost extremities of the sampling frequency distribution. The top-10,000 most frequently generated molecules, for instance, contained 105 of the 252 newly-discovered metabolites, an enrichment of ∼1,500-fold over random expectation (p = 4.3 × 10^−5^, *χ*^2^ test; **Fig. 2k** and **Supplementary Table 1**). This enrichment led us to more carefully examine the 10,000 most frequently generated molecules. This subset contained the vast majority of the known metabolites from the training set (1,905 of 2,046), despite these structures having been withheld in cross-validation. Notably, this subset also included 1,888 metabolites annotated as ‘predicted’ or ‘expected’ in either version 4 or 5 of HMDB (**Fig. 2l**). Because the training set comprised only metabolites that had been experimentally detected in human tissues or biofluids, DeepMet had never seen these metabolites during training, but recapitulated their structures in its output nonetheless. Several of the most frequently sampled ‘predicted/expected’ metabolites were in fact well-studied metabolites that had been misannotated in the HMDB (for instance, ribose 1,5-bisphosphate, meta-octopamine, or 1-carboxyethylisoleucine; **Fig. 2m**), underscoring the ability of our language model to fill the gaps in existing metabolic databases.

In addition to the experimentally detected and ‘predicted/expected’ metabolites, 6,301 of the top-10,000 most frequently sampled metabolites were absent from any version of the HMDB (**Fig. 2k**). These structures are those considered by DeepMet to represent the most plausible extensions of the known metabolome. We therefore hypothesized that many of these structures were indeed as-of-yet undiscovered metabolites.

We sought to experimentally confirm the existence of a subset of these predicted metabolites in human tissues. We obtained or synthesized chemical standards for 83 putative metabolites that ranked within the top-10,000 structures. Each of these standards was profiled individually by LC-MS/MS to determine their retention times and reference MS/MS spectra. These spectra were then searched against metabolomics data from 17,272 urine samples that had been collected by one laboratory during its routine clinical casework using identical chromatographic and mass spectrometric methods. A total of 21 metabolites predicted by DeepMet were successfully identified in human urine by the combination of retention time and MS/MS (**Supplementary Table 2**). These novel metabolites were chemically diverse, including modified or nonproteinogenic amino acids (e.g., N-carbamylproline), acyl-amino acids (e.g., N-succinyl-tryptophan), and indole derivatives (e.g., 1H-indole-3-carboxamide) (**Fig. 2n-p** and **Supplementary Fig. 3a-t**). Many of these novel metabolites were detected in hundreds or thousands of samples (for instance, N-lactoyl-glutamine in 6,455 urine samples; **Supplementary Fig. 3u**).

Thus, DeepMet can fill the gaps in our understanding of metabolism by predicting the existence of as-of-yet undiscovered metabolites, and these predictions can direct the experimental discovery of novel metabolites.

### Assigning novel structures to unidentified peaks

These experiments introduce a structure-centric approach to metabolite discovery, whereby hypothetical metabolites are prioritized by a chemical language model for synthesis and targeted discovery. We also envisioned, however, that DeepMet could support more conventional approaches to metabolite discovery, whereby individual metabolites detected within a biological sample are targeted for structure elucidation on the basis of mass spectrometric data^33^.

We began by asking whether DeepMet could prioritize plausible chemical structures for an unidentified metabolite, given a single measurement as input: that metabolite’s exact mass. To test this possibility, we again used a cross-validation approach, simulating metabolite discovery by withholding a subset of known metabolites from the training set. For each of these held-out metabolites, we filtered the structures generated by DeepMet to those matching the exact mass (± 10 ppm) of the target metabolite. We then tabulated the total frequency with which each of these structures was generated by DeepMet (**Fig. 3a**).

**Fig. 3.**
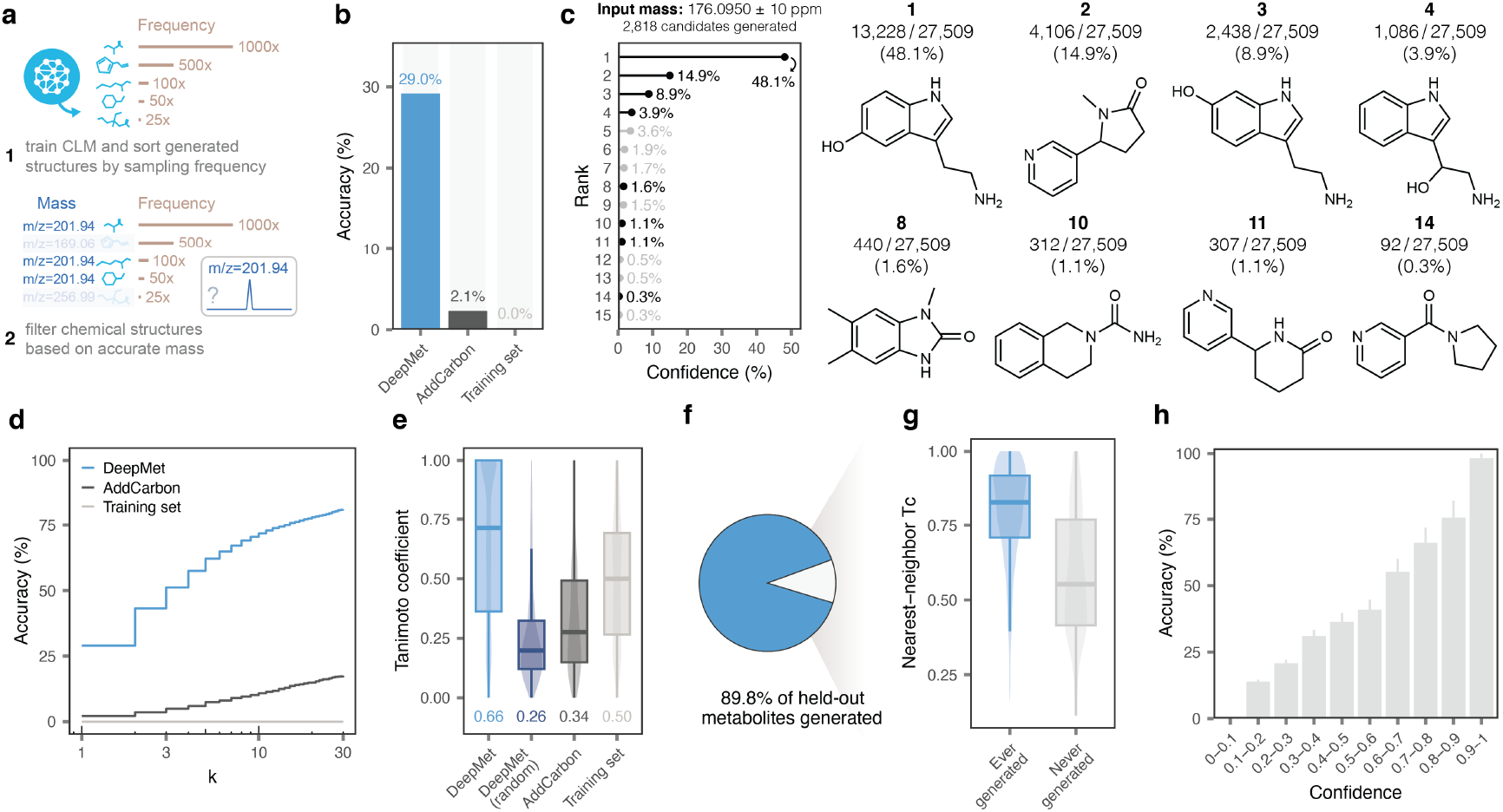
Mass spectrometry-guided prioritization of unknown metabolites. **a**, Schematic overview of the workflow to prioritize chemical structures of unknown metabolites, given an accurate mass measurement as input. **b**, Top-1 accuracy with which the complete chemical structures of held-out metabolites were assigned by DeepMet or two baseline approaches, Add-Carbon or searching within the training set. **c**, Illustrative example demonstrating the use of DeepMet to prioritize candidate structures for an unknown metabolite, based on its accurate mass. A total of 27,509 sampled SMILES strings matched the input mass of 176.0950 *±* 10 ppm, corresponding to 2,818 unique structures. Left, lollipop plot shows the sampling frequencies of the 15 most frequently generated molecules as a proportion of the 27,509 SMILES strings. Right, a subset of the generated molecules is shown, including the four most frequently generated as well as a selection of less frequently generated structures. Structures 1, 2, and 3 are known human metabolites that were not present in the training set. **d**, As in **b**, but showing the top-*k* accuracy curve, for *k* ≤ 30. **e**, Tanimoto coefficient between the structures of held-out metabolites and the top-ranked structures prioritized by DeepMet, random structures generated by DeepMet, or two baseline approaches. **f**, Proportion of held-out metabolites that were ever generated by the language model in cross-validation. **g**, Tc between held-out metabolites and their nearest neighbour in the HMDB, for metabolites that were ever versus never generated in cross-validation. **h**, Proportion of correct structure assignments for held-out metabolites as a function of model confidence.

Across all cross-validation folds, the most frequently generated structure matched that of the held-out metabolite in 29% of cases (**Fig. 3b**). For instance, providing the mass of serotonin (176.0950 Da ± 10 ppm) as input yielded 27,509 SMILES strings, representing 2,818 unique chemical structures; of these, the single most frequently sampled structure was that of serotonin itself (**Fig. 3c**). Because serotonin had been withheld from the training set, this required DeepMet to simultaneously generate the chemical structure of an unseen metabolite, and to prioritize this structure from among thousands of chemically valid candidates.

In cases where the top-ranked structure was not that of the held-out metabolite, the correct structure was often found within a relatively short list of candidates. For example, in the case of serotonin, the second- and third-ranked structures were also those of withheld human metabolites, cotinine and 6-hydroxytryptamine (**Fig. 3c**). Overall, we found that DeepMet ranked the correct structure within its top-3 predictions in 51% of cases and in the top-10 in 72% of cases (**Fig. 3d** and **Supplementary Fig. 4a**). Moreover, when the top-ranked structure was incorrect, it nevertheless typically demonstrated a high degree of chemical similarity to the true metabolite, as quantified either by the Tanimoto coefficient or by continuous molecular embeddings^34^ (**Fig. 3e** and **Supplementary Fig. 4b-g**). Quantitatively similar results were obtained when applying DeepMet to simulated mass-to-charge ratios for each held-out metabolite and allowing matches to multiple positively or negatively charged adducts (e.g., protonated, sodiated, or potassiated adducts in the positive mode; **Supplementary Fig. 4h-j**). Only 10% of held-out metabolites were never reproduced by the language model, and these metabolites tended to demonstrate a low degree of structural similarity to any other metabolite in the training set (**Fig. 3f-g**).

To contextualize the performance of our language model, we compared DeepMet to the AddCarbon baseline proposed by Renz *et al*.^35^. Although this simple approach has frequently outperformed more sophisticated generative models^35^, we nonetheless found that DeepMet markedly outperformed AddCarbon on all metrics (**Fig. 3b,d-e** and **Supplementary Fig. 4c-e**). Structures prioritized by DeepMet also demonstrated a higher degree of structural similarity to the held-out metabolites than isobaric known metabolites, reflecting the model’s ability to generalize beyond the training set into unseen chemical space (**Fig. 3b,d-e** and **Supplementary Fig. 4c-e**).

We also asked whether the confidence scores emitted by DeepMet were well-calibrated, and found that these confidence scores indeed correlated well with the likelihood that any given structure assignment was correct (**Fig. 3h** and **Supplementary Fig. 4k**). This observation highlights a particularly useful property of DeepMet: namely, that its most confident predictions are expected to be the best candidates for experimental follow-up.

We then turned again to the 313 newly-discovered metabolites added in version 5 of the HMDB, and asked whether DeepMet would demonstrate similar performance in this prospective test set. This is a challenging task, as these metabolites are structurally distinct from those in the training set, to the extent that the two sets of metabolites can easily be separated by a random forest classifier in cross-validation (**Supplementary Fig. 4l**). Nonetheless, DeepMet demonstrated comparable performance in this prospective test set, achieving a top-1 accuracy of 22.9%, and a top-10 accuracy of 61.7% (**Supplementary Fig. 4m-p**). For instance, DeepMet correctly prioritized the structures of chemically diverse metabolites such as chenodeoxycholic acid glucuronide, metanephrine 2-sulfate, or adipoylglycine from their exact masses alone (**Supplementary Fig. 4q-s**).

Together, these experiments establish that DeepMet can simultaneously generate and prioritize candidate structures for unidentified metabolites detected by mass spectrometry.

### Integration of DeepMet and MS/MS

Because it is impossible to distinguish between isomeric metabolites with the same formula based on accurate mass information alone, most mass spectrometry-based metabolomics workflows rely on MS/MS for metabolite identification. We therefore next sought to integrate our language model with MS/MS data.

A number of existing computational approaches leverage MS/MS data to search databases of known chemical structures^36^. One such method, CFM-ID^37,38^, learns from a training dataset of experimental MS/MS spectra and their associated structures to predict MS/MS spectra for unseen chemical structures. Applying CFM-ID to a database of known chemical structures produces an *in silico* MS/MS spectral library that can be used to identify compounds by comparing predicted and experimental spectra. We reasoned that this approach could be adapted to search in a database of novel metabolites generated by DeepMet. Moreover, as the sampling frequency in DeepMet effectively prioritized structures based on accurate mass alone, we envisioned that both this sampling frequency and the MS/MS spectral match could be integrated for enhanced accuracy (**Fig. 4a**).

**Fig. 4.**
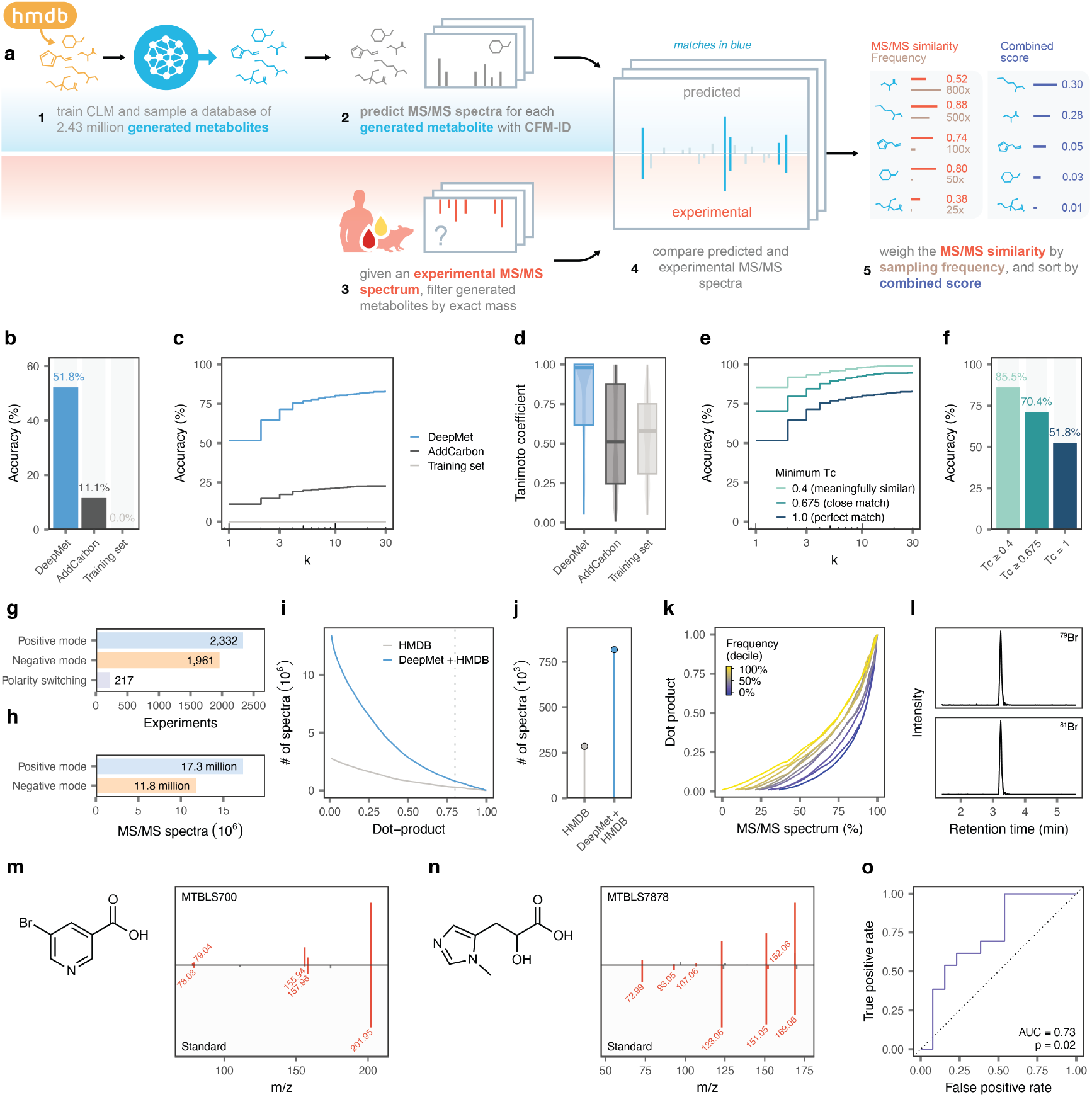
Structure elucidation of unknown metabolites via MS/MS. **a**, Schematic overview of the workflow for automated structure elucidation via MS/MS. **b**, Top-1 accuracy with which the complete chemical structures of held-out metabolites were assigned by the combination of DeepMet with CFM-ID in the Agilent MS/MS library for positive ion mode spectra, as compared to the combination of CFM-ID with two baseline approaches, AddCarbon or searching within the training set. **c**, As in **b**, but showing the top-*k* accuracy curve, for *k* ≤ 30. **d**, Tanimoto coefficient between the structures of held-out metabolites and the top-ranked structures prioritized by the combination of CFM-ID with DeepMet or two baseline approaches. **e**, As in **c**, but also showing the top-*k* accuracy when considering prioritized structures with minimum Tanimoto coefficient of 0.4 or 0.675 as matches. **f**, As in **b**, but also showing the top-1 accuracy when considering prioritized structures with minimum Tanimoto coefficient of 0.4 or 0.675 as matches. **g**, Total number of experiments comprising the human blood metabolome meta-analysis dataset, grouped by polarity. **h**, Total number of MS/MS spectra comprising the human blood metabolome meta-analysis dataset, shown by polarity. **i**, Number of MS/MS spectra in the human blood metabolome dataset linked to a chemical structure when searching against a database of predicted spectra for known human metabolites only (“HMDB alone”), or a combined library containing both known and generated metabolites (“HMDB + Deep-MetDB”). **j**, As in **i**, but for a minimum cosine similarity of 0.8. **k**, Cumulative distribution of cosine similarities between predicted and experimental spectra in the human blood metabolome dataset, for generated metabolites binned into deciles by their sampling frequencies. **l**, Extracted ion chromatograms for an unidentified peak in the human blood metabolome dataset assigned by DeepMet as a bromine-containing metabolite and its ^81^Br isotopologue. **m**, Left, structure of 5-bromonicotinic acid; right, mirror plot showing the similarity between MS/MS spectra from the human blood metabolome dataset (top) versus a synthetic standard (bottom). Fragment ions found in both spectra are shown in red. **n**, Left, structure of N1-methyl imidazeolelactic acid; right, mirror plot showing the similarity between MS/MS spectra from the human blood metabolome dataset (top) versus a synthetic standard (bottom). Fragment ions found in both spectra are shown in red. **o**, ROC curve showing the separation of patients with sepsis from healthy controls by the abundance of N1-methyl imidazeolelactic acid in the MT-BLS7878 dataset.

To test this possibility, we applied CFM-ID to predict MS/MS spectra for 2.4 million novel metabolites generated by DeepMet (the subset of anticipated structures generated at least five times, which included virtually all of the held-out metabolites; **Supplementary Fig. 5a-c**). We again simulated unknown metabolite structure prediction through cross-validation, and found that the incorporation of MS/MS data further improved accuracy. The combination of DeepMet and CFM-ID correctly assigned the exact chemical structures for 52% and 49% of held-out metabolites in the positive and negative ion modes, respectively (**Fig. 4b** and **Supplementary Fig. 5d**). The addition of MS/MS information also robustly increased the number of cases in which the correct structure was ranked among the top-3 or top-10 candidates; the chemical similarity between the predicted and true metabolite structures; and the proportion of spectra for which a close or meaningfully similar match^39^ was retrieved (**Fig. 4c-e** and **Supplementary Fig. 5e-n**). We observed similar performance in a second dataset of MS/MS spectra, or when using alternative machine-learning methods for MS/MS prediction (**Supplementary Fig. 5i-n**).

Over the last decade, thousands of untargeted metabolomics experiments in human tissues and biofluids have been deposited to public repositories. We reasoned that the combination of DeepMet with MS/MS database search could provide a mechanism to systematically annotate the unknown metabolites within these datasets at the repository scale, and thereby begin to uncover the enormous metabolic diversity that has been captured, but not identified, within existing metabolomics data^40^.

To explore this possibility, we first assembled a repository-scale resource of human metabolomics data, focusing on peripheral blood. We identified a total of 4,510 metabolomic analyses of human blood, from which 29.1 million MS/MS spectra were extracted (**Fig. 4g-h** and **Supplementary Fig. 5o-r**).

We first tested the hypothesis that adding structures generated by DeepMet to an *in silico* MS/MS spectral library would increase the number of MS/MS spectra that could be putatively annotated with a chemical structure, as compared to searching only for known metabolites. To this end, we searched the human blood metabolome data against a library comprising predicted MS/MS spectra for all structures in HMDB, or a combined library including predicted MS/MS spectra for both HMDB and DeepMet structures. Searching against the combined library markedly increased the number of MS/MS spectra that could be assigned a putative (MSI level 3) annotation^41^ at any threshold. For instance, the inclusion of DeepMet structures approximately tripled the number of experimental spectra that could be matched to a predicted MS/MS spectrum with a cosine similarity of 0.8 or greater (**Fig. 4i-j**)^42^. We also tested the hypothesis that structures generated more frequently by DeepMet would be disproportionately likely to be detected in published metabolomic datasets. Consistent with this expectation, we observed a strong relationship between the sampling frequency of a putative metabolite and its propensity to match an experimentally collected MS/MS spectrum (**Fig. 4k**).

Together, these results position DeepMet as a powerful hypothesis-generating resource for metabolite discovery in repository-scale metabolomics data. To demonstrate that these hypotheses can enable retrospective metabolite discovery, we sought to experimentally corroborate a subset of novel metabolite annotations. We initially focused on an unidentified metabolite that DeepMet had annotated as a brominated derivative of nicotinic acid, and which demonstrated an isotopic pattern consistent with the presence of bromine (**Fig. 4l**)^43^. We confirmed the structure assigned by DeepMet by comparison to a synthetic standard (MSI level 2 annotation; **Fig. 4m**). Similarly, we confirmed the annotation of an unidentified peak as an N-methylated derivative of imidazolelactic acid in a metabolomic dataset from patients with sepsis (**Fig. 4n**)^44^. Notably, the abundance of this metabolite significantly separated patients with sepsis from healthy controls (**Fig. 4o** and **Supplementary Fig. 5s**), underscoring the potential to discover new biomarkers by re-interrogating published datasets with DeepMet.

### Structure elucidation from multidimensional metabolomics data

Because there are inherent limitations to the confidence with which metabolites can be identified through re-examination of published datasets without access to the original samples, we sought to apply DeepMet to a newly-collected metabolomic dataset that would allow for unambiguous metabolite identification by comparison to chemical standards profiled on an identical analytical setup (**Fig. 5a**). We opted to perform these studies in mice, owing to the ease of accessing many different tissues for metabolomic profiling, the possibility of experimental manipulation, and the assumption that most endogenous metabolic pathways would be conserved from mouse to human.

**Fig. 5.**
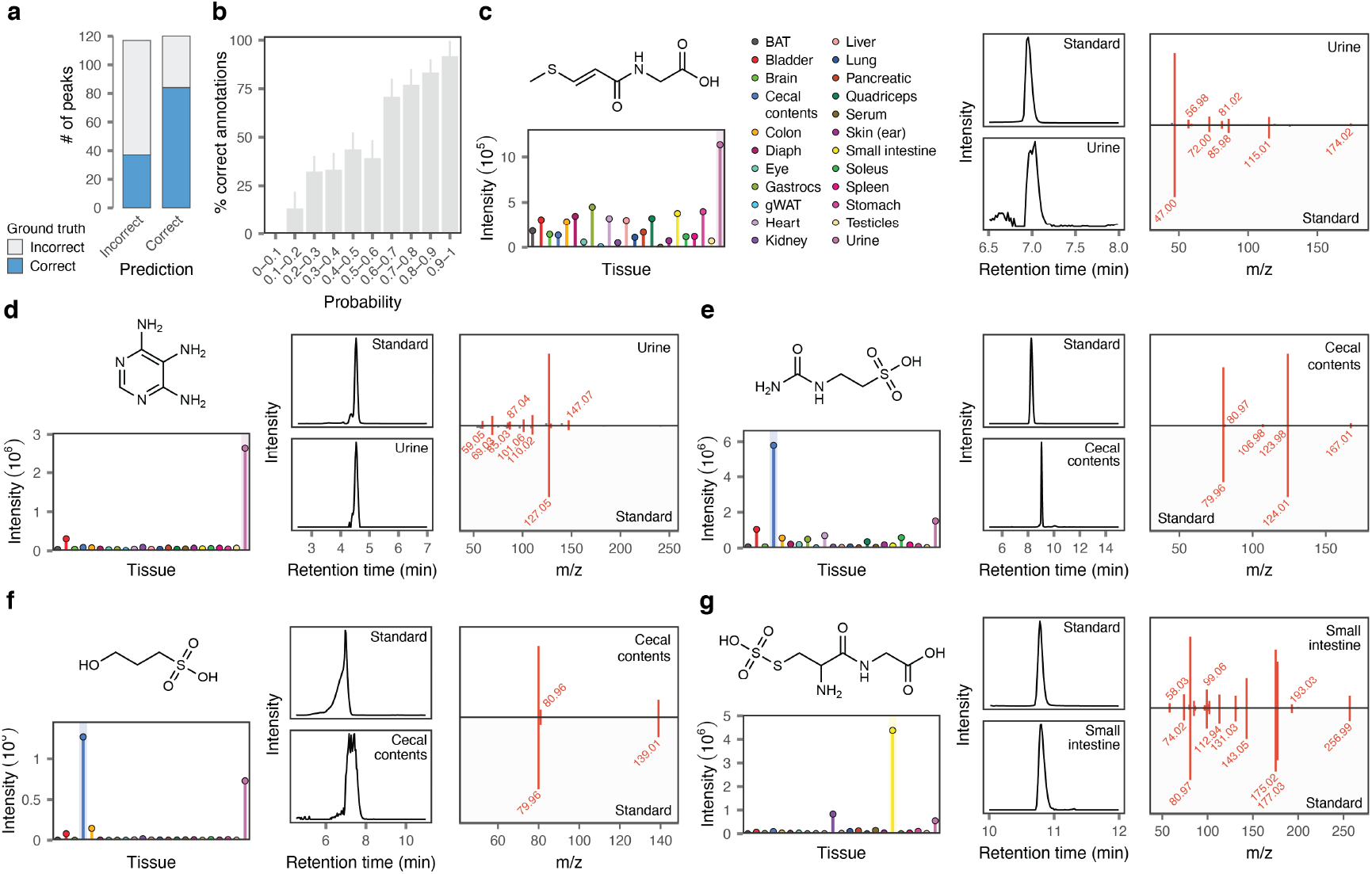
Metabolite discovery in mouse tissues. **a**, Proportion of correct structure assignments for held-out metabolites by a meta-learning model, shown separately for annotations predicted to be correct versus incorrect. **b**, Proportion of correct structure assignments for held-out metabolites as a function of meta-learning model confidence. **c**, Left, MS1 intensity of 3-(methylthio)acryloyl-glycine across 23 mouse tissues. Middle, extracted ion chromatograms (EICs) for the 3-(methylthio)acryloyl-glycine synthetic standard, top, and in mouse urine, bottom. Right, mirror plot showing the similarity between MS/MS spectra from 3-(methylthio)acryloyl-glycine synthetic standard versus the experimental spectrum from mouse urine. **d**, As in **c**, but for 4,5,6-triaminopyrimidine. **e**, As in **c**, but for N-carbamyl-taurine. **f**, As in **c**, but for 3-hydroxypropane-1-sulfonic acid. **g**, As in **c**, but for S-sulfocysteinylglycine.

We profiled the metabolomes of 23 mouse tissues and biofluids by LC-MS/MS. After an initial round of filtering to discard isotopic peaks, adducts, in-source fragments, and artefacts^45^, a total of 4,814 peaks were detected that represented presumptive metabolites (**Supplementary Fig. 6a-b**). Of these, just 250 (5.2%) could be identified by comparison to an in-house library of metabolite standards, whereas the remaining 94.8% remained unidentified (**Supplementary Fig. 6c**).

We first leveraged these identifications to benchmark the accuracy of metabolite discovery with DeepMet within murine tissues, again simulating structure elucidation of unknown metabolites through cross-validation. The combination of DeepMet and CFM-ID correctly identified 49% of the known metabolites withheld from the training set.

To further corroborate the performance of DeepMet, we studied a subset of peaks that were annotated as known metabolites by DeepMet, but for which the corresponding standards were absent from our library. Because both DeepMet and CFM-ID were trained in cross-validation, none of these known metabolites were present in the training sets of either model. We obtained standards for 97 of these metabolites, and experimentally validated 58 of these annotations (60%; **Supplementary Table 3** and **Supplementary Fig. 6**).

Metabolomic profiling collects additional sources of information that are typically not recorded in MS/MS spectral libraries, including retention times and isotopic patterns at the MS1 level. We hypothesized that this additional data could further increase the accuracy of metabolite discovery. To this end, we reasoned that we could train a second machine-learning model to predict whether each structure assigned by DeepMet represented a correct or incorrect annotation. We implemented this meta-learning approach by training a random forest classifier in cross-validation to identify correct annotations by integrating multiple sources of information, including the confidence emitted by the chemical language model, the similarity between predicted and experimental MS/MS spectra, the isotope pattern match at the MS1 level, and the discrepancy between predicted and experimentally measured chromatographic retention times. This meta-learning approach further increased the accuracy of metabolite annotation to 70% (**Fig. 5a**). Moreover, annotations that were assigned a higher probability by the meta-learning model were commensurately more likely to be correct, suggesting a mechanism to move beyond the ordinal levels that are currently used to communicate confidence in metabolite annotation^41,46^ and quantitatively prioritize high-confidence identifications (**Fig. 5b** and **Supplementary Fig. 7a**).

### Metabolite discovery in mouse tissues

We then deployed DeepMet to assign chemical structures to all 4,654 unidentified peaks in the mouse tissue dataset. To validate a subset of the proposed structures, we purchased or synthesized reference standards and profiled these under identical LC-MS/MS conditions. These experiments confirmed the structures of 16 novel mammalian metabolites (**Supplementary Table 2**).

Novel metabolites discovered with DeepMet were structurally diverse. For instance, we identified a series of N-acyl amides, such as 3-(methylthio)acryloyl-glycine, N-isobutyryl histamine, N-isobutyryl methionine, or (2-(4-hydroxyphenyl)acetyl)-aspartic acid (**Fig. 5c** and **Supplementary Fig. 7b-d**). Other novel metabolites were nucleotide or nucleoside derivatives, such as methylthioinosine, 2’-O-methyl-5-methyluridine, or a triaminopyrimidine that resembled formamidopyrimidines produced by oxidative DNA damage^47^ (**Fig. 5d** and **Supplementary Fig. 7e-f**). A fourth series included sulfonate-containing metabolites such as N-carbamyl-taurine, 3-hydroxypropane-1-sulfonic acid, and homotaurine (**Fig. 5e-f, Supplementary Fig. 7g** and **Supplementary Fig. 8a**). Novel metabolites additionally encompassed carbohydrate derivatives (2-sulfoglycerate, (2-aminoethyl)phosphate-1-hexopyranose, 6-O-sulfo-hexopyranose, and glycerylphosphorylethanol), nonproteinogenic dipeptides (S-sulfocysteinylglycine and N-acetyl-phenylalanylleucine), and dicarboxylic acids (7-hydroxyazelaic acid; **Fig. 5g** and **Supplementary Fig. 7h-m**). Novel metabolites were significantly more tissue-specific than known metabolites (p = 1.1 ∼ 10^−5^, t-test; **Supplementary Fig. 7n**), an observation that may explain why the former had eluded discovery until now.

DeepMet also identified a series of putatively novel metabolites that were revealed on closer review of the literature to be known metabolites absent from HMDB (and, in several cases, even PubChem), including S-sulfohomocysteine^48^, thioprolylglycine^49^, 3-sulfopropanoic acid^50^, glutathione-cysteinylglycine mixed disulfide^51^, 3-carboxymethyl-thiolactic acid^52^, N-acetyl-S-carboxymethyl-cysteine^53^, and diacetylputrescine, which was discovered during the preparation of this manuscript^54^ (**Supplementary Fig. 7o-u**). That DeepMet recapitulated the existence of metabolites not captured in existing maps of the metabolome underscores its ability to fill the gaps within these maps, and raises the possibility that DeepMet could facilitate AI-guided curation efforts to more comprehensively catalogue the known metabolome.

A subset of the chemical structures assigned to specific peaks by DeepMet were found to be mismatches on the basis of MS/MS or retention time data acquired from mouse tissues versus synthetic standards. We hypothesized that some of these new structures might, in fact, represent *bona fide* metabolites, just not the specific peaks observed in the mouse tissues. Indeed, four of these metabolites matched peaks in human urine based on MS/MS and LC retention time, leading to the discovery of four additional human metabolites (**Supplementary Fig. 9a-d** and **Supplementary Table 2**).

Motivated by this observation, we searched all of the reference MS/MS spectra acquired in this study against all human metabolomics data—a total of 35,530 datasets, collectively comprising 367.5 million MS/MS spectra—deposited to the MetaboLights and Metabolomics Workbench repositories. This search tentatively identified three additional metabolites, including a glutamyl conjugate of the nucleoside acadesine that was identified in 643 samples, and brought the total number of metabolites discovered in these studies to 47 (**Supplementary Fig. 9e-j**).

### Origins of the novel metabolites

We finally sought to further characterize a subset of the novel metabolites. To identify metabolites that originate from the diet, we collected metabolomics data from the cecal contents in mice fed standard chow, which is rich in dietary metabolites, or a diet comprising purified macromolecules with few metabolites. To identify metabolites that are produced by the microbiota, we collected metabolomics data from the feces of mice treated with broad-spectrum antibiotics and untreated controls. Finally, to establish the biosynthetic origins of novel metabolites, we infused mice with ^13^C-labeled precursors, including glucose, methionine, cysteine, and serine.

These experiments situated several of the novel metabolites at the nexus of diet, the microbiome, and host metabolism (**Fig. 6** and **Supplementary Fig. 8b**). For instance, 3-methylthioacrylic acid is known as a metabolite of methionine in soil-dwelling Streptomyces bacteria^55^. In mice, 3-(methylthio)acryloyl-glycine demonstrated a significant decrease after antibiotic treatment and incorporated ^13^C-methionine, suggesting that gut microbes may encode parallel biosynthetic pathways to those in soil bacteria. N-carbamyl-taurine likewise demonstrated reduced abundance in mice treated with antibiotics, but also decreased in mice fed a purified diet, suggesting contributions from both the diet and the microbiome. This metabolite also incorporated a single carbon from ^13^C-glucose, suggesting the carbamyl group itself is derived from glucose metabolism, likely via glucose oxidation to carbon dioxide and subsequent incorporation of bicarbonate into the carbamyl group. In contrast, 4,5,6-triaminopyrimidine was abundant in standard chow but virtually absent from mice fed a purified diet, did not respond to perturbation of the microbiome, and did not incorporate any isotopically labelled precursors, indicating that this is an exclusively diet-derived metabolite. Finally, S-sulfocysteinylglycine incorporated ^13^C-cysteine and ^13^C-serine, and did not respond to perturbations of the diet or microbiota, allowing us to assign this as an endogenous metabolite.

**Fig. 6.**
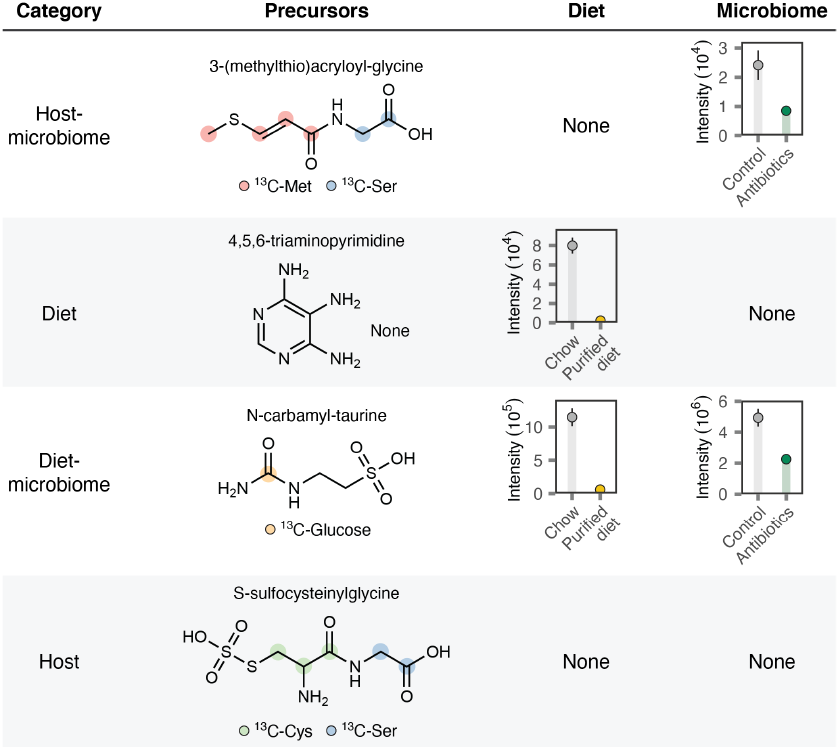
Origins of selected novel metabolites. Far left, inferred origins of each metabolite. Middle left, structures of novel metabolites and positional labelling from the indicated infused ^13^C-labelled precursors. Positional labelling is manually inferred from the structure and experimental isotope labeling patterns in **Supplementary Fig. 8b**. Middle right, metabolite intensities in cecal contents from mice fed chow versus purified (casein) diets. Far right, metabolite intensities in the feces of mice treated with broad-spectrum antibiotics versus untreated controls. None, no significant difference (p < 0.05 and two-fold change).

Collectively, these data experimentally validate our framework for the AI-guided discovery of unknown metabolites from mass spectrometry-based metabolomics data by showcasing the application of DeepMet to reveal dozens of novel mammalian metabolites.

## Discussion

Despite dramatic advances in analytical technologies, large parts of the metabolome remain unexplored. Mass spectrometry-based metabolomics routinely detects thousands of unidentified small molecule-associated peaks in human tissues or biofluids, but unknown metabolites are rarely characterized because of the effort required for structure elucidation of even a single novel metabolite. Here, we address this gap through the development of DeepMet, a language model trained on the chemical structures that populate the known metabolome. We demonstrate that DeepMet has learned the biosynthetic logic embedded within the structures of known metabolites, and can leverage this understanding to anticipate the existence of as-of-yet undiscovered metabolites. In turn, we show that DeepMet’s understanding of metabolism opens up new approaches to metabolite discovery. We deploy DeepMet to discover dozens of novel human and mouse metabolites that span diverse chemotypes.

DeepMet introduces several paradigms that have little conceptual precedent in the study of metabolism. First, we demonstrate the possibility of anticipating the chemical structures of metabolites that likely exist, but which have not yet been discovered. In turn, we show that this approach is able to systematically fill gaps in existing maps of the metabolome. DeepMet recapitulates the existence of well-studied metabolites that were absent from or misannotated within the HMDB due to curation errors, and of newly-discovered metabolites that were added to the HMDB in a later release^32^. We provide a resource of 34.4 million generated structures, and their corresponding sampling frequencies, to facilitate the discovery of metabolites prioritized as those most likely to account for the chemical “dark matter” of the metabolome.

Second, we show that DeepMet enables new paradigms for the interpretation of scalar mass-to-charge ratios. By leveraging DeepMet to generate a large database of metabolite-like chemical structures, and then filtering this database on the basis of a mass-to-charge ratio, we prioritize the structures most likely to account for this mass spectrometric signal. These prioritizations are well-calibrated and remarkably accurate, even in the absence of any other analytical data. This functionality may prove particularly powerful as a hypothesis-generating tool for metabolomic techniques in which MS1 spectra are the primary basis for metabolite annotation: for instance, imaging mass spectrometry^56^ or intraoperative techniques such as rapid evaporative ionization mass spectrometry^57^. More broadly, this approach allows scalar m/z values to be conceptualized as rich distributions over plausible biogenic structures, including those absent from any existing database of chemical structures. As such, this paradigm significantly expands the conceptual repertoire of tools for mass spectrometric data interpretation.

Third, we demonstrate the possibility of integrating language model-guided prioritization of unknown metabolites with existing approaches that compare MS/MS spectra against databases of chemical structures. DeepMet is agnostic to the specific approach by which chemical structures are matched to MS/MS spectra^38,58,59^. We leverage an *in silico* MS/MS library for 2.4 million metabolites anticipated by DeepMet to annotate 29.1 million experimental MS/MS spectra compiled from 4,510 published metabolomic analyses of human blood, and find that DeepMet substantially increases the number of MS/MS spectra that can be linked to a chemical structure at any confidence level. With vast quantities of published metabolomics data now deposited in public repositories, this provides an attractive approach to identify metabolic “dark matter” that went unidentified at the time of the original data acquisition. DeepMet can also enable metabolite discovery in newly-collected datasets, as we demonstrate by discovering and characterizing a series of chemically diverse metabolites in mouse tissues. That DeepMet can catalyze the unbiased discovery of metabolites spanning diverse chemotypes differentiates it from approaches such as reverse metabolomics, in which expert human biochemists elaborate derivatives of a given scaffold via combinatorial synthesis to enable valuable but narrow exploration of metabolite-like chemical space (**Supplementary Fig. 10**)^9^.

Fourth, we introduce a meta-learning approach that integrates the outputs of multiple independent machine-learning models to distinguish correct from incorrect metabolite annotations. This approach provides a principled framework to integrate chemical language models with sources of information (e.g., predicted MS/MS spectra, retention times, and theoretical isotopic distributions) that are currently treated in isolation or combined in a heuristic, often *ad hoc* manner. This meta-learning approach not only increases the overall accuracy of metabolite annotation, but also produces well-calibrated probabilities, raising the possibility of moving beyond the ordinal confidence levels that are currently used to communicate confidence in metabolite annotations^41,46^ and ultimately towards false discovery rate-controlled annotation of unknown metabolites.

DeepMet is not without limitations. Our language model was trained and evaluated on the structures of human metabolites, and evolutionarily distant applications (for instance, to plant or bacterial metabolism) will likely require bespoke models. Learning from the structures of known metabolites allows DeepMet to anticipate unexpected connections between known biosynthetic pathways, but implies inherent limitations for its ability to anticipate metabolites that originate from divergent, as-of-yet undiscovered biosynthetic routes. The finding that some chemical structures assigned by DeepMet were in fact previously undiscovered human metabolites that had been incorrectly matched to MS/MS spectra points to opportunities to more tightly integrate DeepMet with analytical data; despite extensive effort, predicting MS/MS spectra or retention times from chemical structures remain open problems.

## Methods

### Training dataset

A training dataset of known human metabolites was obtained from the Human Metabolome Database (HMDB), the largest and most comprehensive database of human metabolism^24^. Chemical structures were downloaded from the HMDB website in XML format (version 4.0; file ‘hmdb_metabolites.xml’). The HMDB assigns each metabolite to one of four classes: quantified, detected, expected, or predicted. Of the 114,222 metabolites recorded in this XML file, the vast majority fell into the ‘expected’ or ‘predicted’ classes (*n* = 95,202 and 9,929, respectively), indicating that they had not actually been experimentally detected in human tissues or biofluids. These classes instead include structures identified in cell or tissue cultures or in other species, structures predicted based on rule-based enzymatic derivatizations of known human metabolites, and structures predicted based on combinatorial enumeration (for instance, of lipid head groups and acyl/alkyl chains). To avoid conflating predictions made by our chemical language model with an orthogonal set of predictions based on chemical reaction rules or combinatorial enumeration, we trained our language model exclusively on metabolites annotated as ‘detected’ or ‘quantified.’ Moreover, we found that among the 8,970 detected or quantified metabolites, the vast majority (6,791) of these were lipids. Because comprehensive profiling of lipids generally relies on a distinct set of analytical approaches as compared to efforts to comprehensively profile small (polar) metabolites, and because the preponderance of lipids led language models trained on this dataset to almost exclusively generate structurally simple molecules with long acyl chains, we excluded lipids from the training set. This was achieved by removing structures assigned to the ClassyFire superclass “Lipids and lipid-like molecules,” using annotations included in the metabolite XML file^60^. These filters yielded a training set of 2,046 small molecule metabolites that had been experimentally detected in human tissues or biofluids.

The SMILES strings for these 2,046 metabolites were parsed using the RDKit, and stereochemistry was removed. Salts and solvents were removed by splitting molecules into fragments and retaining only the heaviest fragment containing at least three heavy atoms, using code adapted from the Mol2vec package^61^. Charged molecules were neutralized using code adapted from the RDKit Cookbook, after which duplicate SMILES (for instance, stereoisomers or alternatively charged forms of the same molecule) were discarded. Molecules with atoms other than Br, C, Cl, F, H, I, N, O, P or S were removed, and molecules were converted to their canonical SMILES representations. The resulting canonical SMILES were then tokenized by splitting the SMILES string into its constituent characters, except for atomic symbols composed of two characters (Br, Cl) and environments within square brackets, (such as [nH]), and any SMILES containing tokens found in 10 or fewer structures was removed, on the basis that a language model was unlikely to learn how to use these tokens correctly from such a small number of training examples. Metabolites were subsequently split into ten cross-validation folds, and data augmentation was then performed on each fold by enumeration of 30 non-canonical SMILES for each canonical SMILES string in the training fold^62^. This approach takes advantage of the fact that a single chemical structure can be represented by multiple different SMILES strings, and was used here on the basis of previous studies showing that this data augmentation procedure led to more robust chemical language models, particularly when training these models on small datasets^63,64^.

To prospectively evaluate DeepMet predictions, we obtained the structures of a further 313 experimentally detected metabolites that were added to version 5.0 of the HMDB^32^. These metabolites were extracted by applying the same filters as described above to the metabolite XML file from version 5.0, and then removing structures also found in the version 4.0. We additionally removed several thousand exogenous and largely synthetic compounds that had been identified through a text mining approach^65^.

### Language model architecture and training

Our approach to generating novel, metabolite-like chemical structures was based on the use of a language model to generate textual representations of molecules in the SMILES format^25^. As in previous work^17,18,66,67^, we trained a recurrent neural network (RNN) on the SMILES strings of the molecules in our training set. SMILES were tokenized as described above, such that the vocabulary consisted of all unique tokens detected in the training data, as well as start-of-string and end-of-string characters that were prepended and appended to each SMILES string, respectively. The language model was then trained in an autoregressive manner to predict the next token in the sequence of tokens for any given SMILES, beginning with the start-of-string token. Language models based on the long short-term memory (LSTM) architecture were selected on the basis of their excellent performance in previous studies, whereby these were found to outperform both alternative models based on RNNs (e.g., gated recurrent units) as well as more complex models based on the transformer architecture^64,68,69^. LSTMs were implemented in PyTorch, adapting code from the REINVENT package^70^. The architecture consisted of a three-layer LSTM with a hidden layer of 1,024 dimensions, an embedding layer of 128 dimensions, and a linear decoder layer. Models were trained to minimize the cross-entropy loss of next-token prediction using the Adam optimizer with default parameters, a batch size of 64, and a learning rate of 0.001. Ten percent of the molecules in the training set were reserved as a validation set and used for early stopping with a patience of 50,000 minibatches.

To further address the data-limited context of the human metabolome, we employed a transfer learning strategy that we reasoned would first allow our model to learn the syntax of the SMILES representation and subsequently adapt this understanding to the generation of new metabolite-like chemical structures. This was achieved by first pre-training the LSTM until convergence on drug-like small molecules from the ChEMBL database, using the same early stopping criteria as above^71^. ChEMBL (version 28) was obtained from ftp.ebi.ac.uk/pub/databases/chembl/ChEMBLdb/latest/chembl_28_chemreps.txt.gz and processed as described above, except with a single round of non-canonical SMILES enumeration rather than 30. After pre-training using the same stopping criterion as described above, the model was fine-tuned on the HMDB training set, without freezing any layers.

### Metabolite-likeness of generated molecules

We carried out a series of analyses to first establish that the language model had indeed learned to generate novel, metabolite-like structures. To this end, we first trained a chemical language model as described above on a single cross-validation split of the HMDB, then sampled 500,000 SMILES strings from the trained model, and removed those corresponding to known metabolites from the training set. To visualize the areas of chemical space occupied by generated molecules and known metabolites, we implemented an approach based on nonlinear dimensionality reduction. Briefly, we computed a continuous, 512-dimensional representation of each molecule using the Continuous and Data-Driven Descriptors (CDDD) package^34^ (available from http://github.com/jrwnter/cddd). These continuous, 512-dimensional descriptors are derived from a machine translation task in which recurrent neural networks are used to translate between enumerated and canonical SMILES in a sequence-to-sequence modelling framework, a task that forces the latent space to encode the information required to reconstruct the complete chemical structure of the input molecule. We then sampled CDDD descriptors for an equal number of known metabolites and generated molecules, then embedded the CDDD descriptors for both sets of molecules into two dimensions with UMAP^26^, using the implementation provided in the R package ‘uwot’ with the ‘n_neighbors’ parameter set to 5.

To more quantitatively evaluate the chemical similarity of generated molecules to known metabolites, we used a supervised machine learning approach to test whether the two sets of molecules could be distinguished from one another on the basis of their structures. This was achieved by again sampling an equal number of known metabolites and generated molecules, computing extended-connectivity fingerprints with a diameter of 3 and a length of 1,024 bits, and then splitting the resulting fingerprints into training and test sets in an 80/20 ratio. A random forest classifier was then trained in cross-validation to distinguish between known metabolites and generated molecules, using the implementation in scikit-learn. The performance of the classifier was measured using the area under the receiver operating characteristic curve (AUROC), and statistical significance was assessed using the Wilcoxon rank-sum test^72^.

Third, we evaluated whether the molecules generated by the language model overlapped with an orthogonal set of enzymatic biotransformations of known metabolites that had been predicted in a rule-based manner by BioTransformer^27^. BioTransformer comprises a knowledgebase of enzymatic reaction rules that are used to predict generic biotransformation products of endogenous metabolites or xenobiotics based on phase I and II metabolism, promiscuous enzymatic metabolism, and gut microbial metabolism, as well as a machine-learning framework to specifically predict human CYP450-calyzed phase I metabolism of xenobiotics^73^. BioTransformer was applied recursively to the training set in order to generate biotransformation products after one to four steps of enzymatic reactions, and the total fraction of these predictions that were recapitulated by DeepMet was quantified. For this analysis, we used the sample of 1 billion SMILES from all cross-validation folds described below, rather than 500,000 from a single cross-validation split.

Fourth, we computed the Tanimoto coefficient (Tc) between each generated molecule and its nearest neighbor in the training set, again using 1,024-bit extended connectivity fingerprints of radius 3 to calculate the Tc. As a negative control, for each generated molecule, we drew at random a molecule with the same molecular formula from PubChem. The nearest-neighbor Tc was then computed for molecules sampled from PubChem in order to provide a baseline against which the enrichment for metabolite-like chemical structures within the language model output could be compared.

### Sampling frequencies of generated molecules

We initially sought to leverage the trained language model for metabolite discovery by identifying the generated molecules that it viewed as the most plausible extensions of the training set, in analogy to the use of protein language models to forecast the emergence of new protein sequences. Because the use of non-canonical SMILES enumeration implied that multiple SMILES strings could be generated for any given novel structure, and because we found that different SMILES representations of the same novel metabolite were often sampled with very different losses, we drew a very large sample of SMILES strings from the trained model and tabulated the frequency with which each novel chemical structure appeared in this output. This was achieved by drawing samples of 100 million SMILES strings from language models trained on each of the ten cross-validation folds, for a total of 1 billion SMILES. The sampled molecules were then parsed with the RDKit, invalid outputs were discarded, and the frequency with which each canonical SMILES appeared in the model output was tabulated. Sampling frequencies were then averaged across all 10 cross-validation folds, removing molecules in the training set from the language model output for each fold before calculating the average such that all of the analyses described below excluded molecules reproduced from the training set.

To evaluate the relationships between the sampling frequency and metabolite-likeness, generated molecules were then divided into six bins on the basis of sampling frequency, and a series of metrics were calculated that quantified the similarity between the generated molecules in this bin and the molecules in the training set. First, we calculated the nearest-neighbor Tc between each generated molecule and the training set, as described above, and tested for a significant trend with increasing sampling frequency using the Jonckheere-Terpstra test. Second, we again quantified the overlap between generated molecules and rule-based enzymatic transformations predicted by BioTransformer within each sampling frequency bin. Third, we measured the chemical similarity between the generated molecules and the training set as quantified by the Fréchet ChemNet distance^31^. This metric is calculated from the hidden representations of molecules learned by a neural network trained to predict biological activities in thousands of biological assays recorded in ChEMBL, ZINC, and PubChem, and therefore captures both structural properties as well as inferred biological activity; it was previously found to be among the most reliable metrics for evaluating generative models of small molecules^64^ and is included in multiple benchmark suites^74,75^. Fourth, we determined the Murcko scaffolds of generated molecules and the training set^30^, and then calculated the Jensen-Shannon distance between the scaffold distributions of the training set and generated molecules in each frequency bins^64^.

### Anticipation of undiscovered metabolites

To test whether the frequency with which novel molecules were generated could be used to prioritize as-of-yet undiscovered metabolites, we again leveraged the cross-validation set up described above, whereby 10% of the training set was withheld at a time to simulate the appearance of unknown metabolites and frequencies were averaged over cross-validation folds. We then quantified the extent to which sampling frequency alone would separate the held-out metabolites from the background of all generated molecules, using ROC curve analysis and excluding metabolites from the training set. The same analysis was repeated in a prospective setting for the metabolites newly added in version 5.0 of the HMDB, excluding all version 4.0 metabolites. In addition to ROC analysis, we calculated the fold enrichment of newly-discovered metabolites within the top-10 to 100,000 most frequently sampled molecules over random expectation, and evaluated statistical significance using a *χ*^2^ test.

### Structure-centric discovery of predicted metabolites

To experimentally confirm the existence of as-of-yet undiscovered human metabolites prioritized by DeepMet, we leveraged a resource of large-scale human metabolomics data collected by the Provincial Toxicology Centre at the British Columbia Centre for Disease Control as part of its routine clinical operations. Clinical urine samples were subjected to full-scan mass spectrometry as part of routine urine drug screening. Samples were received in sterile urine containers or vacutainers, with no preservatives. Samples were assigned anonymized identifiers for all analyses described here. The study was approved by the UBC Clinical Research Ethics Board (#H22-02722).

Urine samples were analyzed by liquid chromatography-high-resolution mass spectrometry. Samples were hydrolysed using IMCSzyme genetically modified *β*-glucuronidase at 60°C for one hour and filtered using a Biotage Isolute PPT+ protein precipitation plate. After cooling, acetonitrile was added to wash the filter. The acetonitrile was evaporated and the extract reconstituted using methanol:type I water (1:1, v:v). 1 *μ*L was injected on a Thermo Scientific Vanquish LC coupled to a Q Exactive Hybrid Quadrupole Orbitrap mass spectrometer (Waltham, MA, USA). Chromatographic separation was achieved using a Thermo Scientific Accucore Phenyl-Hexyl Column (2.1 × 100 mm, 2.6Å) using a gradient elution. Mobile phase A was 2 mM ammonium formate with 0.1% formic acid in Type I water. Mobile phase B was 2 mM ammonium formate with 0.1% formic acid in a 1:1 (v:v) mixture of acetonitrile and methanol. The flow rate was 0.5 mL/min. The total run time was 12.5 minutes. The column and the autosampler temperatures were set at 40°C and 10°C, respectively. Full scan with targeted data-dependent MS^2^ (full MS/dd-MS^2^) was performed in the positive electrospray ionization mode with an inclusion list containing over 200 drugs. The top eight most intense precursors were selected for fragmentation, unless one or more masses from the inclusion list was detected, in which case those masses were prioritized for fragmentation. The sheath gas flow-rate and the auxiliary gas flow-rate were set at 60 and 20 au, respectively. The spray voltage was set at 3000 V. The capillary and the auxiliary gas heater temperatures were set at 380°C and 375°C, respectively. The S-lens RF was set to 60 V.

To prioritize generated metabolites for discovery, we cross-referenced the 6,301 novel structures within the top-10,000 most frequently sampled molecules with catalogues of commercially available compounds. A total of 107 standards were acquired from Mcule, and standards for two additional predicted metabolites that were not available from commercial suppliers were selected for custom synthesis on the basis of manual review (N-lactoyl-glutamine and N-lactoyl-serine, as described below; “Chemical synthesis”).

The compounds were diluted with methanol to a final stock concentration of 1 mg/mL. These stock solutions were further diluted to concentrations of 100 ng/mL or 1 *μ*g/mL with methanol:water 1:1, v:v. Each of the standards was then analyzed individually using the same chromatographic and mass spectrometric methods that were used to profile clinical samples. The resulting data files were then manually inspected to determine retention times and extract reference MS/MS spectra; 26 standards did not afford high-quality MS/MS spectra (at least two fragments with intensities greater than 1% of the base peak) and were discarded at this stage. The resulting library of 81 reference spectra and their retention times was then queried against the mass spectrometric data from all 17,272 clinical urine samples. Initial identification of the predicted metabolite standards was performed on the the basis of a dot-product of 0.75 or greater between reference and experimental MS/MS spectra and a retention time difference of less than 15 seconds, which was followed by manual inspection to corroborate these matches.

### Prioritization of unknown metabolite structures from accurate masses

The finding that the sampling frequency of any given novel, generated structure was correlated with its metabolite-likeness led us to further hypothesize that we could leverage the sampling frequency to suggest chemical structures for unannotated signals in an untargeted metabolomics experiment. Specifically, we posited that given some experimental measurement as input, such as an accurate mass, we could filter the language model output to a subset of molecules matching this measurement, and rank this subset of generated molecules in descending order by sampling frequency in order to produce a ranked list of candidates. We tested this possibility in cross-validation by filtering the language model output with the exact mass of each held-out metabolite, allowing for a mass error of up to 10 ppm, and ranking the resulting structures by frequency.

To evaluate the accuracy of this approach, we computed the fraction of held-out HMDB version 4.0 metabolites for which the correct structure was found within the top-*k* candidates, systematically varying the value of *k* between 1 and 30. In addition, we calculated the Tc between the topranked candidate and the held-out molecule; whereas extended connectivity fingerprints were used for all other analyses in the paper, here RDKit fingerprints were used because these had previously been calibrated based on a user study of expert chemists to define quantitative thresholds that approximated these chemists’ subjective judgements of ‘meaningful similarity’ or a ‘close match’ between true and predicted structures^39^. The use of chemical similarity thresholds allowed us to also identify cases in which the language model nominated a structure closely related to the ground truth (for instance, where the correct scaffold of the held-out metabolite was predicted, but a single functional group was misplaced). As a secondary measure of chemical similarity, we computed the Euclidean distance between CDDD descriptors^34^. We additionally hypothesized that held-out metabolites that the model failed to ever generate would tend to occupy more distinct regions of chemical space with few similar structures in the training set; we tested this hypothesis by calculating the nearest-neighbor Tc between each metabolite in version 4.0 of the HMDB and the remainder of the training set. Each of the above analyses was then repeated for the newly-discovered metabolites added to version 5.0 of HMDB.

Because, to our knowledge, no existing approaches are capable of prioritizing structures of unknown human metabolites from accurate mass measurements alone, we sought to place the performance of DeepMet in context by comparing our model to simple baseline approaches. First, to assess the model’s ability to generalize beyond the chemical space of the training set, we searched by accurate mass in the training set itself, with the recognition that this would yield a top-*k* accuracy of 0% by definition, but with the goal of comparing the Tanimoto coefficients between the true molecule and structures prioritized by the language model to plausible matches from the training set. A substantial fraction of held-out metabolites had no molecules with matching masses in the training set; these were omitted from the evaluation. Second, we applied the AddCarbon approach that has been advocated as a simple and universal baseline for more complex generative models^35^. This model inserts a carbon atom (‘C’) at random positions within the SMILES representation of a molecule from the training set. If the insertion of the carbon atom produces a valid SMILES string and the corresponding molecule is not itself in the training set, then the modified SMILES string is retained. Surprisingly, this trivial baseline was found to outperform numerous more complex approaches to molecule generation on the distribution learning tasks proposed in one widely used benchmark suite^74^. We adapted the Python source code available from https://github.com/ml-jku/mgenerators-failure-modes to exhaustively enumerate all possible ‘AddCarbon’ derivatives of the training set metabolites. Invalid SMILES were removed, the remaining SMILES were converted to their canonical forms, and derivatives that were also found in the training set were removed. For both baselines, the same 10 ppm error window was used as for the language model, and when more than one candidate structure matched, the candidates were ordered at random.

We additionally tested whether this prioritization was robust to the presence of multiple positively or negatively charged adducts. This was achieved by computing the protonated or deprotonated mass of the held-out metabolite in the positive or negative modes, respectively, and then searching in the language model output as described above but here considering three adduct types in each mode, including [M+H]^+^, [M+NH_4_]^+^, and [M+Na]^+^ in the positive mode, and [M–H]^−^, [M+Cl]^−^, and [M+FA–H]^−^ in the negative mode.

We further assessed the calibration of confidence scores emitted by DeepMet for any given accurate mass input. Confidence scores were computed by dividing the sampling frequency of the top-ranked molecule from the sum of sampling frequencies of all generated molecules matching the query mass. These scores were then divided into ten bins of equal widths, and the proportion of correct matches within each bin was determined.

### Structure elucidation of unknown metabolites via MS/MS

We next sought to integrate DeepMet with tandem mass spectrometry (MS/MS) data as a means to experimentally differentiate between isobaric metabolites, which cannot be distinguished by accurate mass measurements alone. Our efforts to this end began by applying CFM-ID^38,76^ to create an *in silico* MS/MS library for metabolites generated by DeepMet. CFM-ID is a machine-learning method that is trained on a reference library of MS/MS spectra for known small molecules, and learns to predict MS/MS spectra for unseen chemical structures on the basis of the information within this dataset. During the training phase, for each input molecule, CFM-ID first employs a combinatorial bond cleavage approach to enumerate all theoretically possible fragments. The output of this procedure is a molecular fragmentation graph, in which each node represents a theoretically possible fragment from the parent molecule with one bond cleavage, and each edge (also known as transition) between nodes encodes the chance that one fragment directly produces another fragment through a fragmentation event. The probability of each transition is estimated by parameters that CFM-ID learns from its training data set of known molecules and their associated MS/MS spectra. These parameters are learned by minimizing a negative log-likelihood loss within a training dataset of known molecule-MS/MS spectrum pairs using expectation maximization (EM). Finally, CFM-ID uses the fragmentation graph and associated transition probability estimates for each molecular fragment to reconstruct the corresponding MS/MS spectrum for the input molecule. CFM-ID predicts MS/MS spectra at three different collision energies (at 10 eV, 20 eV, 40 eV) and in both positive and negative ionization modes, functionality which differentiates it from the majority of alternative machine-learning methods for MS/MS spectrum prediction from chemical structures.

To balance performance with the computational requirements necessary to predict MS/MS spectra for tens of millions of generated structures, we limited these predictions to a subset of 2.4 million molecules that were generated at least five times by DeepMet. This threshold was selected by removing molecules sampled less than 2, 3, 4, 5, or 10 times and repeating the analyses of unknown metabolite prioritization based on exact mass information described above, which indicated that the ability of DeepMet to prioritize unknown metabolites from exact masses was minimally affected by removing molecules sampled less than five times.

To evaluate the performance of the combination of DeepMet and CFMID in the context of metabolite discovery, we again implemented a cross-validation strategy whereby the structures and MS/MS spectra of known metabolites were withheld from both of these models to simulate the emergence of a novel metabolite. CFM-ID was trained in ten-fold cross-validation on ESI-QTOF MS/MS spectra from the Agilent MassHunter METLIN Metabolite reference spectral library. These models were used to predict MS/MS spectra for metabolites in the held-out test set for each cross-validation fold. An eleventh model was trained on the entire Agilent MS/MS library and used to predict MS/MS spectra for metabolites without reference spectra in the training set. Spectra predicted at multiple collision energies were merged to produce a single predicted MS/MS spectrum per generated structure. For each reference MS/MS spectrum in the test set, candidates were retrieved from the database of generated metabolites produced by applying DeepMet in cross-validation to the HMDB, and a final score was assigned to each candidate structure by multiplying the confidence assigned by the language model on the basis of the precursor m/z by the dot-product between the predicted and experimental MS/MS spectra. Performance was then evaluated using the same metrics as described above, i.e., the top-*k* accuracy for values of *k* between 1 and 30; the Tc between the top-ranked candidate and the held-out molecule, using RDKit fingerprints; and the top-*k* accuracy when considering predicted metabolites with a Tc ≥ 0.675 (“close match”) or ≥ 0.40 (“meaningfully similar”) as matches^39^. In addition, we applied CFM-ID to structures proposed by the same two baseline methods as above, AddCarbon and searching within the training set, and compared the performance of these approaches to the combination of CFM-ID with DeepMet.

To demonstrate the robustness of our approach to metabolite identification by combining DeepMet with MS/MS prediction, we carried out several additional experiments. First, we re-trained CFM-ID in cross-validation on a second library of MS/MS reference spectra, obtained from the HMDB, and again evaluated performance as described above. Second, we benchmarked alternative models for MS/MS prediction from chemical structures, including FraGNNet^59^ and NEIMS^58^, here employing five-fold rather than ten-fold cross-validation. Each model employs a different approach to MS/MS prediction. CFM-ID models fragmentation as a Markov decision process, and is trained to predict the probability of each fragmentation (i.e., bond cleavage) event. FraGNNet applies a graph neural network to a combinatorial fragmentation graph in order to model mass spectra as distributions over molecule fragments. NEIMS performs MS/MS prediction via vector regression, taking molecular fingerprints as input and passing these through a multilayer perceptron (MLP) to predict binned MS/MS spectra. NEIMS was modified to predict high-resolution MS/MS spectra at a resolution of 0.01 m/z, rather than 1 m/z as originally described by the authors, a modification which required replacing the fully-connected MLP output layer with a low-rank layer to fit the high-resolution model into memory. Both FraGNNet and NEIMS were additionally modified to condition MS/MS prediction on adduct type and collision energy as input, in order to match their output MS/MS spectra with those predicted by CFM-ID. Notably, each of the three models has limitations that precluded the prediction of MS/MS spectra for some generated metabolites. CFM-ID and FraGNNet cannot predict MS/MS spectra for structures with a formal charge. FraGNNet additionally cannot predict MS/MS spectra for structures with more than 60 heavy atoms. NEIMS cannot predict MS/MS spectra for structures with a precursor m/z greater than 1,500 Da. An empty spectrum was assigned to structures that violated these constraints, which comprised 5%, 8%, and 1% of the generated metabolites for CFM-ID, FraGNNet, and NEIMS, respectively.

### Meta-analysis of the human blood metabolome

To showcase the potential for DeepMet to enable metabolite discovery in published metabolomics data at the repository scale, we carried out a meta-analysis of the human blood metabolome. Data was obtained from the MetaboLights database^77^, as this resource requires extensive metadata annotation for each deposited sample, including the species and tissue of origin. An XML record of all studies deposited to MetaboLights was obtained (file ‘eb-eye_metabolights_studies.xml’) and filtered to only mass spectrometry-based metabolomics studies that included at least one sample from human serum, plasma, or whole blood. Complete data depositions for this subset of studies were then downloaded from MetaboLights. The assay-level metadata (‘a_*’ files) were parsed to obtain a complete list of all mass spectrometric runs for all of the human blood metabolome studies and to exclude GC-MS, imaging MS, and targeted MS experiments, inspecting the relevant MTBLS pages and the corresponding publications as necessary to ensure that no LC-MS metabolomics studies were inadvertently removed and manually correct any filenames that were discordant between the assay-level metadata and the deposited raw files. Compressed archives (.tar, .gz, .zip) were decompressed, and vendor-specific file formats (.d, .raw, .wiff) were converted to mzML format using the msconvert utility bundled with ProteoWizard^78^. MS/MS spectra from each run were then extracted and written to MGF files after ensuring the following quality control (QC) criteria were met: at least 50 unique precursor m/z values; at least 100 non-empty MS/MS spectra; both precursor m/z and fragment m/z recorded to at least four decimal places. LC-MS/MS files that did not meet one or more of these filters were manually inspected to ascertain why they did not pass these QC criteria. A handful of duplicate files, representing cases where the same mass spectrometry run was uploaded as part of more than one accession, were identified by their checksums and removed. Finally, sample-level metadata files (‘s_*’), study protocols, and/or the original publications were manually reviewed for each of the files that passed all of the above steps in order to confirm that the run in question was indeed a human blood sample. In total, these steps afforded a resource comprising 29.1 million MS/MS spectra from 4,510 mass spectrometry runs.

All 29.1 million MS/MS spectra were then searched against the resources of spectra predicted by CFM-ID for both known metabolites and molecules generated by DeepMet, retaining only the predicted spectrum with the highest cosine similarity across collision energies for each known or generated structure. We first calculated the total number of human blood MS/MS spectra with at least one match to a predicted MS/MS spectrum above any given cosine similarity threshold between 0 and 1, when considering known metabolites alone or when combining known metabolites with molecules generated by DeepMet. Separately, we asked whether metabolites generated more frequently demonstrated a greater propensity to match to experimentally collected MS/MS spectra. To address this question, we iterated over each experimentally collected MS/MS spectrum and annotated this spectrum on the basis of the linear combination of cosine similarity and DeepMet sampling frequency, as described above, excluding spectra without at least one match to a predicted spectrum with a dot-product greater than zero. We then binned these annotated metabolites by their sampling frequencies into deciles, and computed the proportion of annotations within each decile where the predicted and experimental MS/MS spectra matched with a cosine similarity score exceeding a given threshold between 0 and 1.

We last sought to experimentally corroborate a subset of the annotations made by the combination of DeepMet and CFM-ID by acquiring MS/MS spectra from synthetic standards. We initially focused on a peak detected in MTBLS700^44^ (sample ns94, RT 3.25 min, precursor m/z 201.9492 in negative mode) that was annotated as a brominated derivative of nicotinic acid, for which MS1 data supported the presence of bromine. Synthetic 5-bromonicotinic acid was obtained from Sigma (cat. no. 228435) and a reference MS/MS spectrum was acquired using a Maxis II Q-Tof MS system from Bruker, coupled to an Ultimate 3000 LC system from Thermo Scientific. The LC system was equipped with a Zorbax Eclipse XDB-C18 Solvent Saver Plus column (80 Å, 4.6 × 150 mm, 5 *μ*m) (Agilent, Santa Clara, CA). The standard was dissolved in a 1:1 mixture of MS-grade methanol and water to a final concentration of 100 *μ*M, 5 *μ*L of which was injected into the LC-MS/MS system described above. Chromatographic separation was performed using MS-grade water with 0.1% formic acid as mobile phase A (MPA) and MS-grade methanol with 0.1% formic acid as mobile phase B (MPB). The gradient was programmed as follows with a constant flow rate of 0.5 mL/min: minute 0, 100% MPA; minute 13, 25% MPA; minute 15, 25% MPA; minute 15.2, 100% MPA; minute 18, 100% MPA. To visualize the match between the experimental spectrum from MTBLS700 and the synthetic standard, the former was preprocessed to remove MS2 fragments that were uncorrelated with the MS1 precursor, as described in more detail below, with a minimum Pearson correlation coefficient of 0.95. We also sought to corroborate an annotation of a peak assigned as in MTBLS7878^44^ (sample neg_C13, RT 1.76 min, precursor m/z 169.0596 in negative mode). Synthetic N-methyl imidazolelactic acid was obtained from Enamine (cat. no. Z8914008850) and a reference MS/MS spectrum was acquired as described below (“Metabolite standards”). To evaluate the relationship between the quantitative abundance of this metabolite and disease status in this study, the dataset was re-processed with xcms^79^ and the intensity of this peak was compared between patients with sepsis and healthy controls using ROC curve analysis as implemented in the R package AUC.

### Mouse sample collection

Animal studies adhered to protocols approved by the Princeton University Institutional Animal Care and Use Committee (IACUC). Male C57BL/6 mice (Charles River), aged 10-12 weeks, were maintained on standard mouse chow. On the sample collection day, urine was collected after a 6 h fast. Blood samples were taken via tail snip, kept on ice for up to 60 minutes, and then centrifuged at 10,000 rcf for 10 minutes at 4°C. The plasma was transferred to another tube and stored at –80°C. The mice were then euthanized by cervical dislocation, and tissues were dissected, wrapped in foil, clamped with a Wollenberger clamp precooled in liquid nitrogen, and subsequently immersed in liquid nitrogen. All samples were stored in a –80°C freezer. A total of 23 tissue and fluid samples were collected, including the brain, liver, kidney, spleen, pancreas, gWAT, stomach, small intestine, lung, heart, quadriceps, BAT, soleus, diaphragm, gastrocnemius, colon, skin (ear), testicles, bladder, cecal content, eye, serum, and urine.

### Metabolite extraction

Frozen solid tissue samples were first weighed to aliquot approximately 40 mg of each tissue and then transferred to 2.0 mL Eppendorf tubes on dry ice. Samples were then ground into powder with a cryomill machine (Retsch, Newtown, PA) maintained at cold temperature using liquid nitrogen. Thereafter, for every 30 mg tissue, 1 mL 40:40:20 acetonitrile:methanol:water with 0.5% formic acid was added to the tube, vortexed, and allowed to sit on ice for 10 minutes and 85 *μ*L 15% NH4HCO3 (w:v) was added and vortexed to neutralize the samples^80^. The samples were incubated on ice for another 10 min and then centrifuged at 14000 rpm for 25 min at 4°C. The supernatants were transferred to another Eppendorf tube and centrifuged at 14000 rpm again for 25 min at 4°C with supernatant collected for analysis. For metabolite extraction from serum and urine, frozen samples were allowed to thaw on ice. For 10 *μ*L urine or serum, 200 *μ*L methanol was added and vortexed for 10 seconds, and centrifuged for 25 min. The supernatant was collected, then dried down under N2 stream, and re-dissolved into 200 *μ*L 40:40:20 acetonitrile:methanol:water for analysis.

### Liquid chromatography-mass spectrometry (LC-MS)

LC-MS analysis was performed on a Vanquish UHPLC system coupled with an Orbitrap Exploris 480 mass spectrometer. LC separation was achieved using a Waters XBridge BEH Amide column (2.1 × 150 mm, 2.5 *μ*m particle size, cat. no. 186006724), with column oven temperature at 25°C and injection volume of 5 *μ*L. The method has a running time of 25 minutes at a flow rate of 150 *μ*L/min. Solvent A is 95:5 water:acetonitrile with 20 mM ammonium hydroxide and 20 mM ammonium acetate, pH 9.4. Solvent B is acetonitrile. The gradient is, 0 min, 90% B; 2 min, 90% B; 3 min, 75%; 7 min, 75% B; 8 min, 70%, 9 min, 70% B; 10 min, 50% B; 12 min, 50% B; 13 min, 25% B; 14 min, 25% B; 16 min, 0% B, 20.5 min, 0% B; 21 min, 90% B; 25 min, 90% B^81^. The Exploris 480 mass spectrometer was operated in full scan mode at MS1 level for the 23 metabolite extracts. This allows the relative quantitation of the individual metabolite across all tissues and fluids by ion counts. In addition, the 23 extracts were mixed to generate a “mixture” sample and analyzed using the same LC-MS method. Peak picking was performed from the mixture sample for further analysis. Full scan parameters are: resolution, 120,000; scan range, m/z 70-1000 (negative mode); AGC target, 1e7; IT^max^, 200 ms. Other instrument parameters are: spray voltage 3000 V, sheath gas 35 (Arb), aux gas 10 (Arb), sweep gas 0.5 (Arb), ion transfer tube temperature 300°C, vaporizer temperature 35°C, internal mass calibration on, RF lens 60.

### Peak picking

Thermo Raw data files were converted to mzXML format using the msconvert utility bundled with ProteoWizard^78^. Peak picking for the mixture sample was then performed using El-MAVEN version 12.0^82^ with the following parameters, mass domain resolution 10 ppm, time domain resolution 20 scans, minimum peak intensity 10,000, minimum quality 0.5, minimum signal/blank ratio 3.0, minimum signal/baseline ratio 2.0, minimum peak width 10 scans. The analysis resulted in 17,386 peaks (features) in negative mode defined by their m/z and RT. EIC curves for each peak were then retrieved, plotted and saved as grey-scale images of the same format (.png, 700 by 525 pixels). A computer vision algorithm implemented in Matlab was then used to classify these peaks as reliable versus spurious. The classifier comprises a convolutional neural network (CNN), with similar architecture as previously described^83^, and was trained on the dataset provided by the authors of EVA. The CNN flagged 960 poor-quality peaks, which were removed from the dataset, along with 3,012 duplicate peaks, yielding a table of 14,374 peaks for further annotation.

This peak table, along with the intensities of each peak in each mixture sample, was provided to NetID for metabolite annotation^45^, along with a database of 114,014 known metabolites from HMDB; a table containing the m/z and retention times of 500 metabolite standards; and a transformation rule table describing the formula and mass difference of 84 transformations, as previously described^45^. The penalty for 1 ppm m/z difference between annotated formula and measured m/z was set at –0.5. The propagation and recording thresholds were set at 10 and 5 ppm, respectively. All other parameters were set at their default values. NetID annotated 7,015 peaks as artefactual (including isotopes, adducts, fragments, and ringing artifacts), 2,369 peaks as known metabolites, 2,305 peaks as putative derivatives of known metabolites, and 2,685 peaks as unknowns.

Annotated peaks were then further filtered on the basis of their intensities. The intensities of each peak across all tissues were retrieved, and the most abundant tissue along with its intensity (I_max_) was recorded for each peak. Among known metabolites and their putative derivatives, 2,285 and 1,972 peaks with I_max_ > 10^5^ were selected, respectively. Among unknown peaks, 557 peaks with log_10_(I_max_) > 6.5. These filters yielded a total of 4,814 peaks for targeted MS2 analysis.

### Targeted MS/MS analysis

For each of the 4,814 peaks of interest, signal intensity was retrieved from the full-scan data of the 23 tissue extracts to identify the tissue with the highest signal intensity, and MS/MS was performed for the corresponding tissue extract. Samples were analyzed with a full scan, followed by targeted MS2 scans using an inclusion list in the same LC-MS run. Full scan parameters were: resolution 60,000, range m/z 70-1000, AGC target 1e7, IT_max_ 200 ms. MS/MS parameters were: isolation window 1.7 m/z, collision energies 15, 30, 50 eV, resolution 15000, AGC target 1e6, IT_max_ 300 ms, RT window 3 min.

In complex biological samples, the presence of chimeric MS/MS spectra containing fragments from multiple precursor ions within the isolation window can hinder metabolite identification84,^85^. To deconvolve fragment ions from co-isolated precursors, we implemented a procedure based on the Pearson correlation between MS1 and MS2 ions, with the assumption that only fragment ions whose intensities are correlated with that of the precursor ion originated from this precursor. Each precursor ion yielded multiple MS2 scans spanning a RT window of up to 3 min. The scan associated with the highest MS1 intensity was used to obtain the MS2 spectrum for that precursor ion, from which the m/z and intensity for individual fragment ions were retrieved. For each fragment peak, its EIC curve at the MS2 level was constructed and correlated with the EIC of the precursor ion in MS1, after alignment of the scan times by interpolation and filtering to a RT window of 0.3 min around the scan with the highest MS1 intensity. Fragment ions with a Pearson correlation coefficient less than 0.8 were discarded.

### Meta-learning

The availability of additional sources of information in the mouse tissue dataset to support metabolite annotation, including retention times and isotopic patterns at the MS1 level, suggested an avenue to improve the accuracy of metabolite annotation relative to that which had been achieved using MS/MS alone. To explore this possibility, we devised a metalearning framework to combine DeepMet confidence scores and predicted MS/MS spectra from CFM-ID with retention time and isotope patterns. We first used the combination of DeepMet and CFM-ID to annotate the MS/MS spectra for all 250 known metabolites in the mouse tissue dataset, using the weighted combination of normalized sampling frequency and cosine similarities between predicted and experimental MS/MS spectra as described above. A 5 ppm window of error was used to identify candidates for each precursor m/z, searching for only [M-H]-adducts; as above, any DeepMet or CFM-ID predictions were made by models trained in cross-validation to avoid data leakage between training and test sets. Of these annotations, 48.9% were correct. We then calculated a series of features for each annotation that were provided as input to a random forest classifier that was trained to perform a binary classification task, distinguishing correct from incorrect annotations. The features were as follows: (1) the absolute sampling frequency of the generated molecule; (2) the normalized sampling frequency (i.e., to the sum of sampling frequencies of all candidate generated molecules matching the precursor m/z); (3) the rank of the generated structure by sampling frequency among all candidates generated by DeepMet; (4) the cosine similarity between the experimental MS/MS spectrum and that predicted by CFM-ID for each candidate; (5) the number of peaks matching between the predicted and experimental spectra; (6) the mass error between the experimental and theoretical m/z, in parts per million; (7) the cosine similarity between the theoretical and observed isotope patterns at the MS1 level; and (8) the difference between experimental and predicted retention times. The classifier was trained in ten-fold cross-validation on the set of 250 known metabolites. Calibration was assessed as described above by binning annotations by the probability assigned by the meta-learning model into deciles, and calculating the proportions of correct annotations within each bin.

Prediction of unmeasured retention times for structures generated by DeepMet was achieved using a structure-based retention time prediction model based on a graph neural network (GNN). The GNN model was implemented in PyTorch and the Deep Graph Library (DGL) and comprised four GraphSAGE layers^86^ with a LSTM feature aggregator and 4 dense layers, each with a hidden dimension of 256. The RT prediction model was trained in ten-fold cross-validation on an in-house library of metabolite standards and their retention times to minimize a mean absolute error loss for 1,000 epochs using the Adam optimizer, a batch size of 512, and a learning rate of 0.001.

### Metabolite standards

Synthetic standards for putative chemical structures assigned by DeepMet were obtained from commercial suppliers (Mcule, Enamine) or via chemical synthesis. Catalogue numbers for commercially available compounds are provided in **Supplementary Table 2**. Protocols for chemical synthesis are described below (“Chemical synthesis”). Standards were dissolved into 50:50 methanol:H_2_O at 1 mg/mL. The stock solution was further diluted into 40:40:20 acetonitrile:methanol:H_2_O at 2 *μ*g/mL and analyzed by full-scan LC-MS to determine the retention time on the 25 min HILIC method. The 23 tissue extracts were then re-analyzed side-by-side with the synthesized compounds using full-scan followed by targeted MS2 scans using an inclusion list. Full scan parameters were: resolution 60000, range m/z 70-1000, AGC target 1e7, IT_max_ 200 ms. MS2 parameters are, isolation window 1.7 m/z, collision energies 15, 30, 50 eV or 15, 20, 30 eV, resolution 15000, AGC target 1e6, IT_max_ 300 ms.

Where appropriate, two orthogonal approaches were used to further confirm matches between synthetic standards and metabolites identified in tissue extracts. The first such approach involved differentiating chromatographic drift from slight differences in retention time by spiking the synthetic standard into the corresponding tissue extract to establish whether the two features (retention time, MS1 and MS2) merged into a single peak, as expected. These spike-in experiments were performed for N-carbamyltaurine and glycerylphosphorylethanol. The second such approach involved re-acquiring data from both the synthetic standard and from the tissue extract in positive mode. Data acquired in positive mode is shown in the manuscript for the following metabolites: diacetylputrescine, 2’-O-methyl-5-methyluridine, 4,5,6-triaminopyrimidine, and isobutyryl-histamine.

Thermo raw files were then analyzed by the Xcalibur QualBrowser to determine the retention time for each synthetic standard and visualize the MS2 spectra for the standards and corresponding metabolite peaks in the tissue samples. Raw files were then converted to mzML files using ProteoWizard, and contaminating fragment ions from co-isolated precursors were identified as those that showed low correlation with the precursor m/z and removed, as described above (“Targeted MS/MS analysis”) but with the following modifications: (1) internal mass calibration ions were removed; (2) fragment ions with an absolute difference of less than 1 m/z to the precursor ion were removed as these could not be explained by a neutral loss; (3) the minimum correlation between MS1 and MS2 ions was adjusted in a data-adaptive manner as a function of retention time and MS1 signal intensity; and (4) fragment ions with an absolute intensity greater than 120% of the precursor ion in MS1 were removed.

### Metabolomics of antibiotics-treated mice

Wild type C57BL/6NCrl mice (strain #:027) were obtained at 8 weeks of age from Charles River Laboratories and used for experiments at age 8-15 weeks. Antibiotics were provided in the drinking water for 14 days as a mixture containing ampicillin (1 g/L), neomycin (1 g/L), metronidazole (1 g/L), and vancomycin (0.5 g/L) (all from Sigma-Aldrich). To improve the taste of the drinking water, 0.5% aspartame (from Bulk Supplements) was added to the antibiotic solution, and drinking water with 0.5% aspartame was used as control. To collect fecal samples, mice were restrained and gently massaged on the belly to induce defecation, and the fecal pellets were immediately frozen on dry ice. All samples were stored at –80°C until further analysis.

### Metabolomics of dietary perturbations

To assess the difference between chow and purified diets, 8-week-old male C57BL/6NCrl mice were fed either PicoLab Rodent Diet 20 (5053, *n* = 4) or a standard casein protein based purified diet (Research diets Inc, D11112201i, *n* = 4). After 10 days on the respective diets, feces were collected at 7 AM. The fecal samples were collected fresh and immediately flash frozen. For extraction, feces were ground at liquid nitrogen temperature with a cryomill (Restch, Newtown, PA). The resulting powder was extracted with 40:40:20 methanol: acetonitrile: water (40 ul extraction solvent per 1 mg tissue) for 10 min on ice and centrifuged at 15,000 x g for 10 min.

### Isotope tracing

Infusions were performed in conscious, free-moving mice which had been catheterized in the jugular vein at least 5 days prior. Specifically, male C57BL/6 mice, housed in a normal light cycle and aged 12-16 weeks, were brought to a procedure room at 9 AM. Mice (n = 2 per tracer) were placed in a new cage and the infusion line was connected. For the U13C-glucose infusion, animals were provided food in their new cage, but other animals were not provided food and remained fasting to the end of the infusion. Immediately after connecting the infusion line, it was primed with 14 microliters of infusate to replace the dead volume of saline. At 11 AM, a sample of blood was collected by tail snip, and then the infusion started. Infusion rates and concentrations were designed to target 50% enrichment based on previously published measurements of *F*_circ_ (circulatory flux, also known as rate of appearance; ^13^C_3_-serine, 141.667 mM, 3 μL/min, 2 h; ^13^C_5_-methionine, 33.333 mM, 3 μL/min, 2 h; ^13^C_6_-glucose, 1875 mM, 4 μL/min, 6 h; ^13^C_3_-cysteine, 30 mM, 2.5 μL/min, 6 h)^87,88^. Urine was collected from the animal if it urinated when scruffed at the end of the infusion. The mice were euthanized by cervical dislocation, and tissues were quickly dissected and snap frozen in liquid nitrogen using a Wollenberger clamp. If urine had not already been collected, then urine was withdrawn from the bladder using an insulin syringe. In addition, a set of mice were not infused with anything but handled in parallel to the infused mice as controls.

### Chemical synthesis

Synthetic methods are described in **Supplementary Note 1**.

### Repository search

The observation that several chemical structures assigned by DeepMet yielded poor matches to the corresponding peaks in mouse tissues, but that these putative metabolites could nonetheless be identified in human biofluids via LC-MS/MS analysis of synthetic standards, motivated us to more comprehensively search the reference MS/MS spectra acquired in this study against published human metabolomics data. To this end, we assembled a compendium of 35,530 untargeted metabolomic datasets from human tissues, comprising 367.5 million MS/MS spectra, from the MetaboLights and Metabolomics Workbench repositories. Human files were identified by a combination of automated metadata search followed by extensive manual review to remove non-human samples and blanks. For MetaboLights, study metadata was downloaded in XML format and filtered as described above (“Meta-analysis of the human blood metabolome”) to include only liquid chromatography-mass spectrometry datasets, but here including all tissues rather than limiting this analysis to blood. For Metabolomics Workbench, the REST API was queried first to retrieve all LC-MS studies (endpoint ‘/rest/study/study_id/ST/summary’) and then manually curated in order to subset these to studies of human tissues. These files were then downloaded, decompressed, and converted from vendor-specific formats (e.g., .raw, .d, .wiff) to mzML, and MS/MS spectra from each run were then extracted and written to MGF files after quality control, again as described above. The reference MS/MS spectra acquired from chemical standards were then used to search the entire compendium of published human metabolomics data, using the implementation of the normalized dot-product in the ‘Spectra’ R package^89^, with a precursor m/z tolerance of 20 ppm, a fragment m/z tolerance of 50 ppm, and considering only peaks in the reference spectrum that were within the scan limit of the experimental spectrum. Matches with a cosine similarity greater than 0.8 and at least three matching peaks greater than 1% relative intensity in both spectra were retained and manually inspected to discard unreliable matches.

### Reverse metabolomics

Structures of metabolites discovered via reverse metabolomics were extracted from the mass spectral data and supplementary materials accompanying the paper^9^. For N-acyl amides, detected metabolites were identified by matching compound names from the Figure 2 source data with mass spectra from the GNPS library ECG-ACYLAMIDES-C4-C24-LIBRARY. For acyl esters, detected metabolites were identified by matching compound names from the Extended Data Figure 3 source data with mass spectra from the GNPS library ECG-ACYL-ESTERS-C4-C24-LIBRARY. For bile acid conjugates, compound names provided in Supplementary Tables 1, 2, and 3 were used to identify detected metabolites, and their structures were retrieved from the GNPS mass spectral library BILELIB19. Metabolites discovered by DeepMet or reverse metabolomics were then visualized within the same two-dimensional UMAP representation of metabolite-like chemical space as described above, using the uwot function ‘umap_transform’ to assign coordinates for newly-discovered metabolites in the UMAP shown in **Fig. 1**. ClassyFire was additionally used to classify each metabolite discovered by DeepMet or reverse metabolomics according to the ChemOnt taxonomy, and the total number of classes represented by at least one metabolite discovered by either approach was calculated at each level of the ChemOnt hierarchy.

## Data and code availability

We make all of the data and code generated in this study available as a resource to accelerate the mapping of the human metabolome. An interactive web application, available at http://deepmet.org, allows users to explore the structures of generated molecules and prioritize unknown metabolites based on an accurate mass, inferred molecular formula, or MS/MS spectrum. The raw data, including all of the generated structures and predicted MS/MS spectra, can also be accessed directly via Zenodo (doi: 10.5281/zenodo.12999324). Last, we release a Snakemake pipeline that allows investigators to rapidly train, sample from, and evaluate chemical language models, allowing for the paradigm described here to be extended to other classes of metabolites (available from GitHub at https://github.com/skinniderlab/clm).

## Acknowledgements

This work was supported by funding from Ludwig Cancer Research; Genome Canada, Genome British Columbia, and Genome Alberta (project 284MBO); the Canada Foundation for Innovation (CFI MSIF 42495); the Princeton Language Institute; Princeton Precision Health; the National Institutes of Health (R50CA211437 to W.L.); NSERC; AMII; and CIFAR.

---

**Supplementary Fig. 1.**
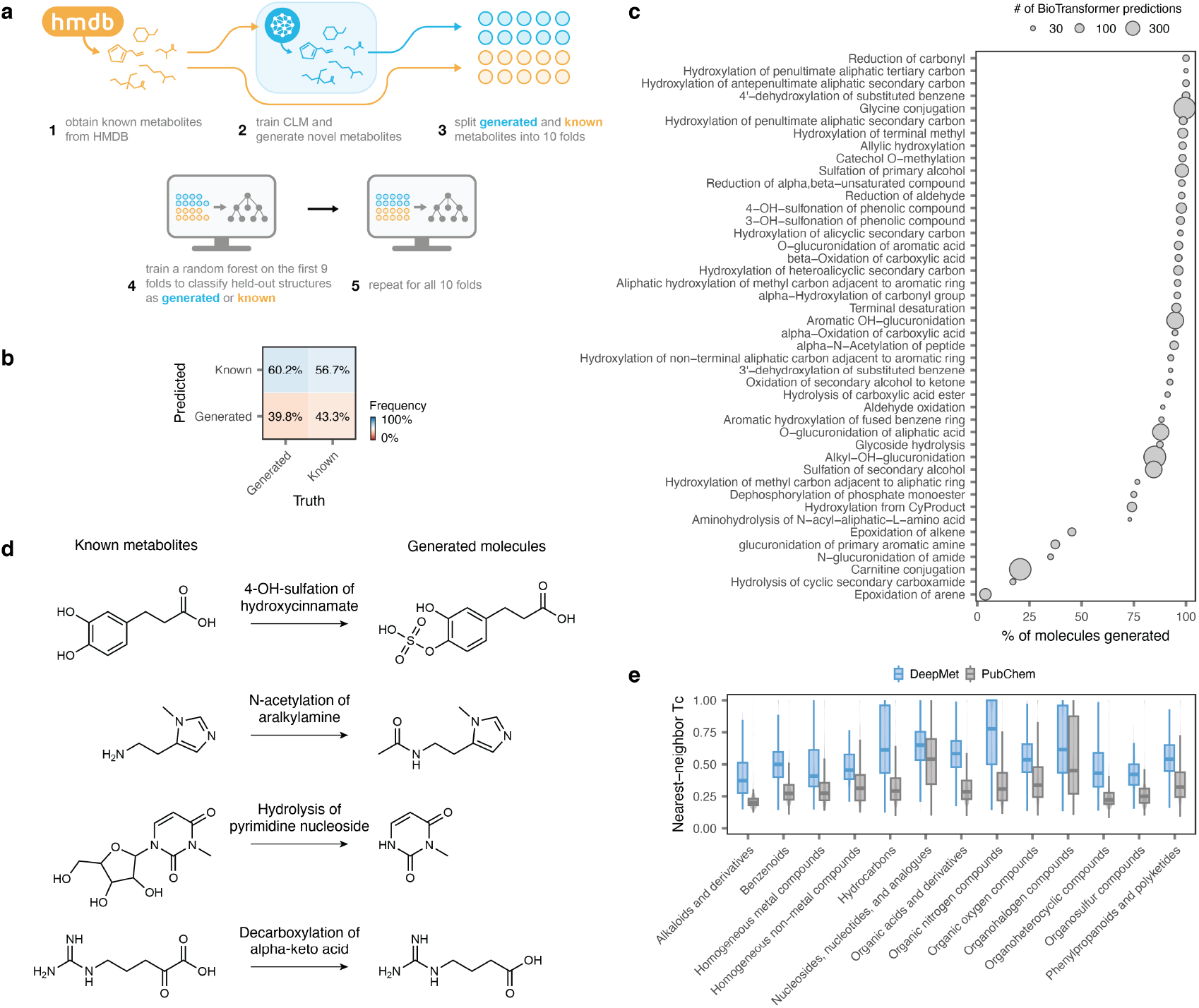
A language model of the human metabolome. **a**, Schematic overview of the random forest classifier trained in cross-validation to distinguish generated molecules from a held-out set of human metabolites, withheld from the language model during training. **b**, Confusion matrix showing predictions by the random forest classifier versus true classes. **c**, Proportion of one-step biotransformations of known human metabolites recapitulated by DeepMet, shown across individual enzymatic reactions. **d**, Examples of enzymatic biotransformations recapitulated by DeepMet in a rule-free manner. **e**, Nearest-neighbor Tc to a known human metabolite for generated molecules versus molecules with identical molecular formulas sampled randomly from PubChem, shown separately for each superclass in the ClassyFire chemical ontology.

**Supplementary Fig. 2.**
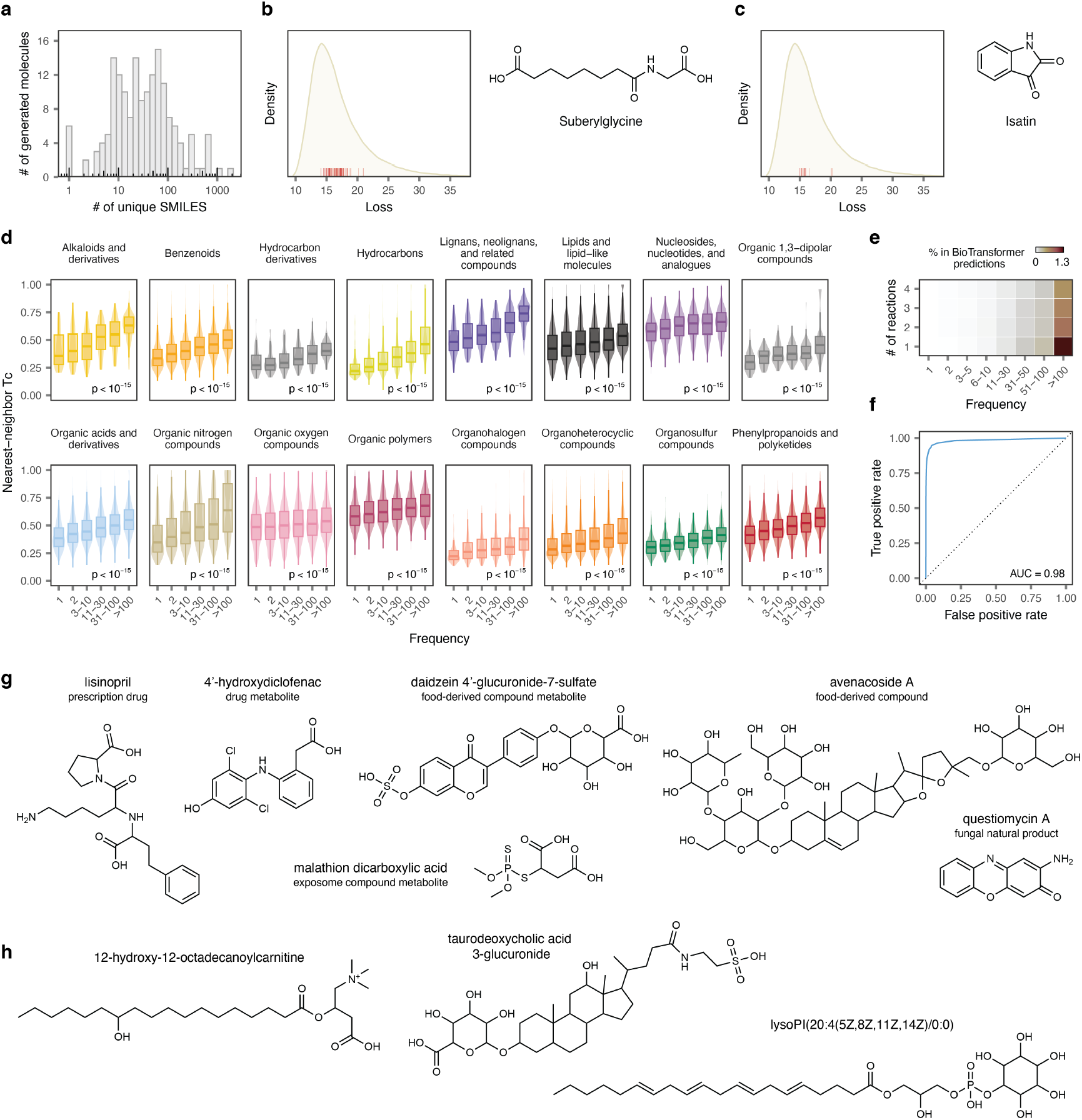
DeepMet anticipates as-of-yet undiscovered metabolites. **a**, Number of unique SMILES generated for each held-out metabolite in the test set, within a sample of 10 million SMILES. **b**, Density plot showing the distribution of losses for all sampled SMILES, with red ticks showing losses for all sampled SMILES corresponding to the structure of suberylglycine, a held-out metabolite from the test set. Right, structure of suberylglycine. **c**, As in **b**, but showing another held-out metabolite, isatin. **d**, Tc between generated molecules and their nearest neighbour in the training set of known metabolites for molecules generated with progressively increasing frequencies, shown separately for each superclass in the ClassyFire chemical ontology. **e**, Heatmap showing the proportion of generated metabolites recapitulating one-to four-step enzymatic transformations of human metabolites predicted by BioTransformer, for molecules generated with progressively increasing frequencies. **f**, ROC curve showing the prioritization of held-out human metabolites from HMDB4 on the basis of sampling frequency, as compared to the background distribution of all generated molecules. **g**, Examples of structures added to version 5 of the HMDB that were not generated by DeepMet, with synthetic or biosynthetic origins outside of endogenous human metabolism. **h**, Examples of *bona fide* human metabolites added to version 5 of the HMDB that were not generated by DeepMet.

**Supplementary Fig. 3.**
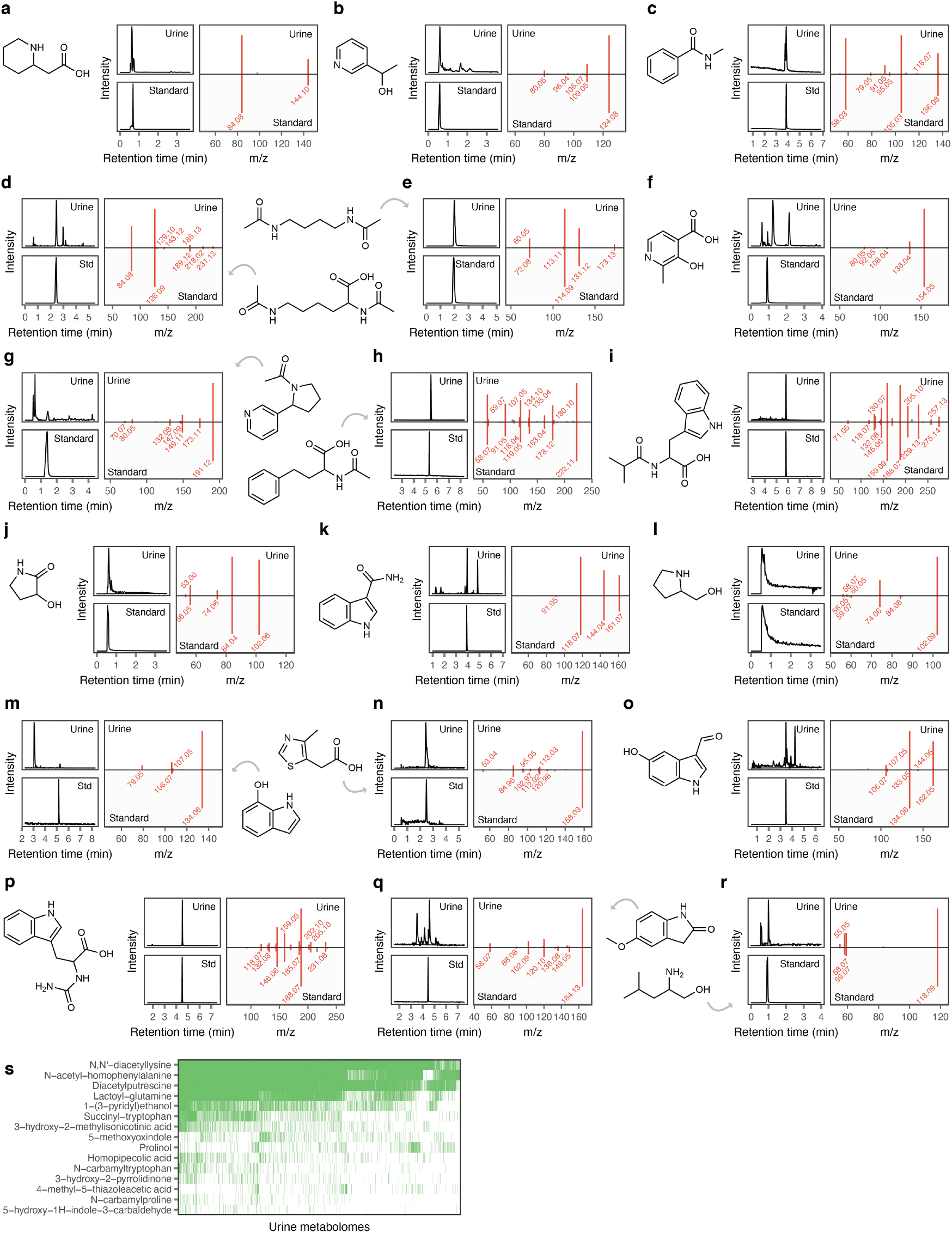
Language-model guided discoveries of additional human metabolites. **a-r**, Additional novel human metabolites predicted to exist by DeepMet and experimentally discovered in human urine, as in **Fig. 2n-p. a**, Homopipecolic acid; **b**, 1-(3-pyridyl)ethanol; **c**, N-methylbenzamide; **d**, N,N’-diacetyllysine; **e**, diacetylputrescine; **f**, 3-hydroxy-2-methyl-4-pyridinecarboxylic acid; **g**, N-acetylnornicotine; **h**, N-acetylhomophenylalanine; **i**, isobutyryl-tryptophan; **j**, 3-hydroxy-2-pyrrolidinone; **k**, indole-3-carboxamide; **l**, prolinol; **m**, 7-hydroxyindole; **n**, 4-methyl-5-thiazoleacetic acid; **o**, 5-hydroxy-1H-indole-3-carbaldehyde; **p**, N-carbamyltryptophan; **q**, 5-methoxyoxindole; **r**, leucinol. **s**, Prevalence of novel metabolites detected in more than 50 urine samples, across all 17,272 urine samples analyzed.

**Supplementary Fig. 4.**
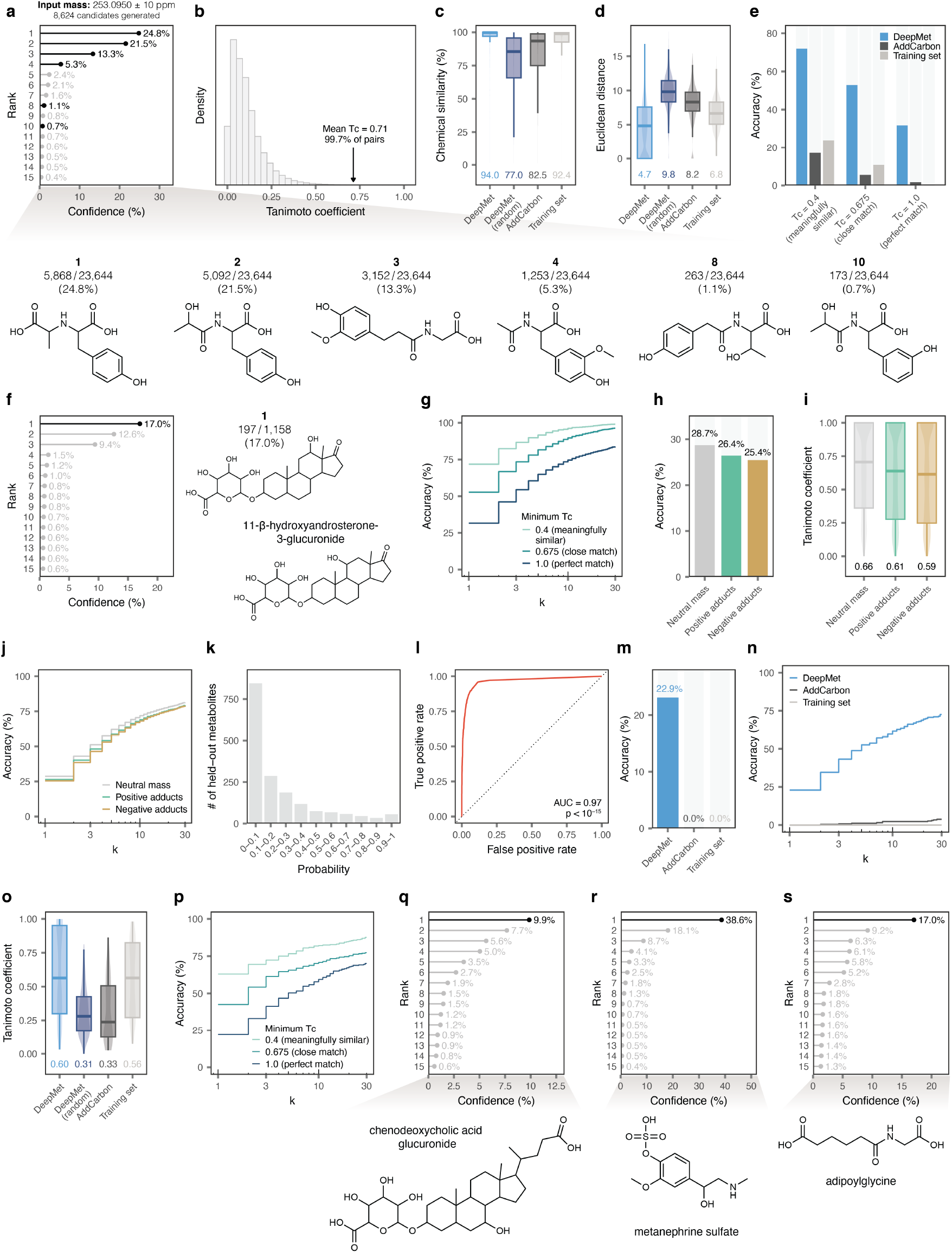
Prioritization of unknown metabolite structures from accurate mass measurements. **a**, Illustrative example demonstrating the use of DeepMet to prioritize candidate structures for an unknown metabolite, in which the correct structure is ranked second. A total of 23,664 sampled SMILES strings matched the input mass of 253.0950 *±* 10 ppm, corresponding to 8,624 unique structures. Top, lollipop plot shows the sampling frequencies of the 15 most frequently generated molecules as a proportion of the 23,664 SMILES strings. Bottom, a subset of the generated molecules is shown, including the four most frequently generated as well as two less frequently generated structures. Here, the held-out metabolite was N-lactoyl-tyrosine (structure 2); however, structure 1, 1-carboxyethyltyrosine, is also a known metabolite. **b**, Histogram showing the distribution of Tanimoto coefficients between random pairs of training set metabolites from version 4 of the HMDB. Arrow, mean Tc between held-out metabolites and structures prioritized by DeepMet. **c**, Tc between the structures of held-out metabolites and the top-ranked structures prioritized by DeepMet, random structures generated by DeepMet, or two baseline approaches, as shown in **Fig. 3e**, but here with the Tc represented as a quantile of the empirical distribution of Tanimoto coefficients between random pairs of training set metabolites. **d**, Euclidean distance between CDDD embeddings for held-out metabolites and the top-ranked structures prioritized by DeepMet, random structures generated by DeepMet, or two baseline approaches. **e**, Top-1 accuracy with which the complete chemical structures of held-out metabolites were assigned by DeepMet or two baseline approaches when considering prioritized structures with minimum Tc of 0.4 or 0.675 as matches. **f**, As in **a**, but showing an example where the top-ranked structure is incorrect but demonstrates a high degree of chemical similarity to the true held-out metabolite. Left, lollipop plot showing sampling frequencies; middle, top-ranked structure; right, true structure of 11-*β*-hydroxyandrosterone-3-glucuronide. **g**, Top-*k* accuracy with which the complete chemical structures of held-out metabolites were assigned by DeepMet when considering prioritized structures with minimum Tc of 0.4 or 0.675 as matches. **h**,Top-1 accuracy with which the complete chemical structures of held-out metabolites were assigned by DeepMet when considering multiple positively or negatively charged adducts. **i**, Tc between the structures of held-out metabolites and the top-ranked structures prioritized by DeepMet when considering multiple positively or negatively charged adducts. **j**, As in **h**, but showing the top-*k* accuracy. **k**, Distribution of model confidence scores assigned to top-ranked structures for each held-out metabolite, given the exact mass of that metabolite as input. **l**, ROC curve showing the performance of a random forest classifier trained in cross-validation to separate metabolites from versions 4 and 5 of HMDB. **m**, Top-1 accuracy with which the complete chemical structures of newly-discovered metabolites added in version 5 of the HMDB were assigned by DeepMet or two baseline approaches. **n**, As in **l**, but showing the top-*k* accuracy curve. **o**, Tc between the structures of newly-discovered metabolites added in version 5 of the HMDB and the top-ranked structures prioritized by DeepMet, random structures generated by DeepMet, or two baseline approaches. **p**, As in **n**, but showing the top-*k* accuracy curves when considering prioritized structures with minimum Tc of 0.4 or 0.675 as matches. **q-s**, Illustrative examples of newly-discovered metabolites added in version 5 of the HMDB that were correctly prioritized by DeepMet, given an accurate mass as input. Top, lollipop plots showing sampling frequencies; bottom, structures of the metabolites.

**Supplementary Fig. 5.**
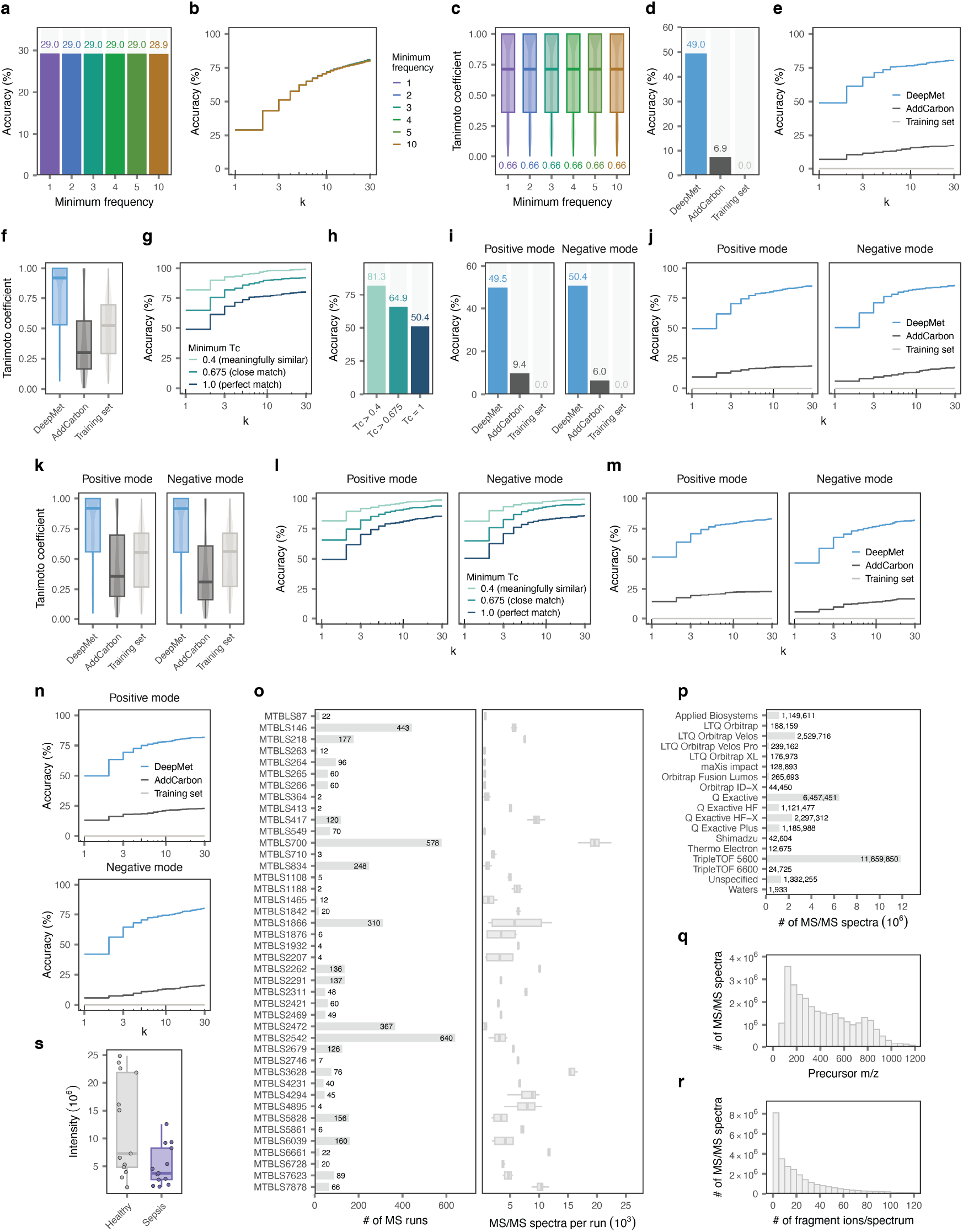
*De novo* structure elucidation of novel metabolites via MS/MS. **a**, Top-1 accuracy with which the complete chemical structures of held-out metabolites were assigned by DeepMet when considering only structures generated at least 1x, 2x, 3x, 4x, 5x, or 10x. **b**, As in **a**, but showing the top-*k* accuracy curve, for *k ≤* 30. **c**, Tc between the structures of held-out metabolites and the top-ranked structures prioritized by DeepMet when considering only structures generated at least 1x, 2x, 3x, 4x, 5x, or 10x. **d**, Top-1 accuracy with which the complete chemical structures of held-out metabolites were assigned by the combination of DeepMet with CFM-ID in the Agilent MS/MS library for negative ion mode spectra, as compared to the combination of CFM-ID with two baseline approaches, AddCarbon or searching within the training set. **e**, As in **d**, but showing the top-*k* accuracy curve, for *k ≤* 30. **f**, Tc between the structures of held-out metabolites and the top-ranked structures prioritized by the combination of CFM-ID with DeepMet or two baseline approaches. **g**, As in **e**, but also showing the top-*k* accuracy when considering prioritized structures with minimum Tc of 0.4 or 0.675 as matches. **h**, As in **d**, but also showing the top-1 accuracy when considering prioritized structures with minimum Tc of 0.4 or 0.675 as matches. **i**, As in **d**, but in the HMDB MS/MS library for positive and negative ion mode spectra. **j**, As in **e**, but in the HMDB MS/MS library for positive and negative ion mode spectra. **k**, As in **f**, but in the HMDB MS/MS library for positive and negative ion mode spectra. **l**, As in **g**, but in the HMDB MS/MS library for positive and negative ion mode spectra. **m**, As in **e**, but when using FraGNNet to predict MS/MS spectra in the positive ion mode. **n**, As in **e**, but when using NEIMS to predict MS/MS spectra in the positive ion mode. **o**, Number of mass spectrometry runs, left, and number of MS/MS spectra per run, right, in the human blood metabolome meta-analysis dataset. **p**, Instruments used to acquire MS/MS spectra in the human blood metabolome meta-analysis dataset. **q**, Distribution of precursor m/z’s in the human blood metabolome meta-analysis dataset. **r**, Number of fragment ions per MS/MS spectrum in the human blood metabolome meta-analysis dataset. **s**, Abundance of N1-methyl imidazolelactic acid in patients with sepsis versus healthy controls in the MTBLS7878 dataset.

**Supplementary Fig. 6.**
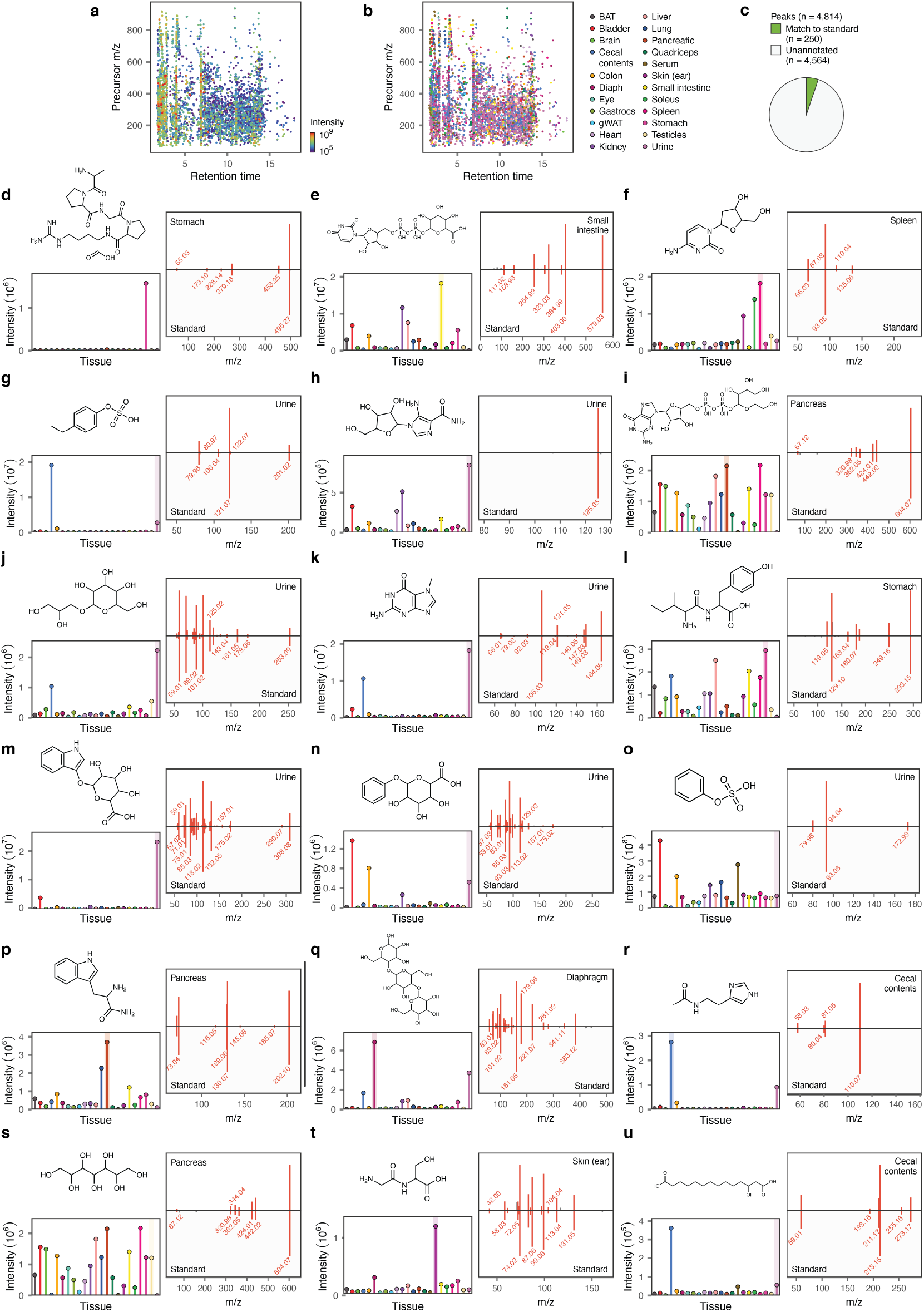
Overview of the mouse tissue dataset and DeepMet-guided identification of simulated unknown metabolites. **a-b**, Visualization of the 4,814 peaks in the mouse tissue metabolomics data, colored by maximum precursor intensity across 23 tissues, left, or the tissue with the highest intensity, right. **c**, Proportion of the 4,814 peaks unambiguously identified by a match to a chemical standard. **d-u**, Representative examples of known metabolites absent from our in-house library of chemical standards but which were correctly identified by the combination of DeepMet and CFM-ID despite being withheld from the training sets of both models. Left, chemical structure of the metabolite (top) and MS1 intensity across 23 mouse tissues (bottom), colored as in b. Right, mirror plot showing the similarity between MS/MS spectra from the synthetic standard versus the experimental spectrum in mouse tissues. **d**, APGPR enterostatin; **e**, UDP glucuronic acid; **f**, deoxycytidine; **g**, 4-ethylphenylsulfate; **h**, acadesine; **i**, GDP glucose; **j**, galactosylglycerol; **k**, 7-methylguanine; **l**, isoleucyltyrosine; **m**, indoxyl glucuronide; **n**, phenylglucuronide; **o**, phenol sulfate; **p**, tryptophanamide; **q**, amylose; **r**, N-acetylhistamine; **s**, peracitol; **t**, glycylserine; **u**, 3-hydroxytetradecanedioic acid.

**Supplementary Fig. 7.**
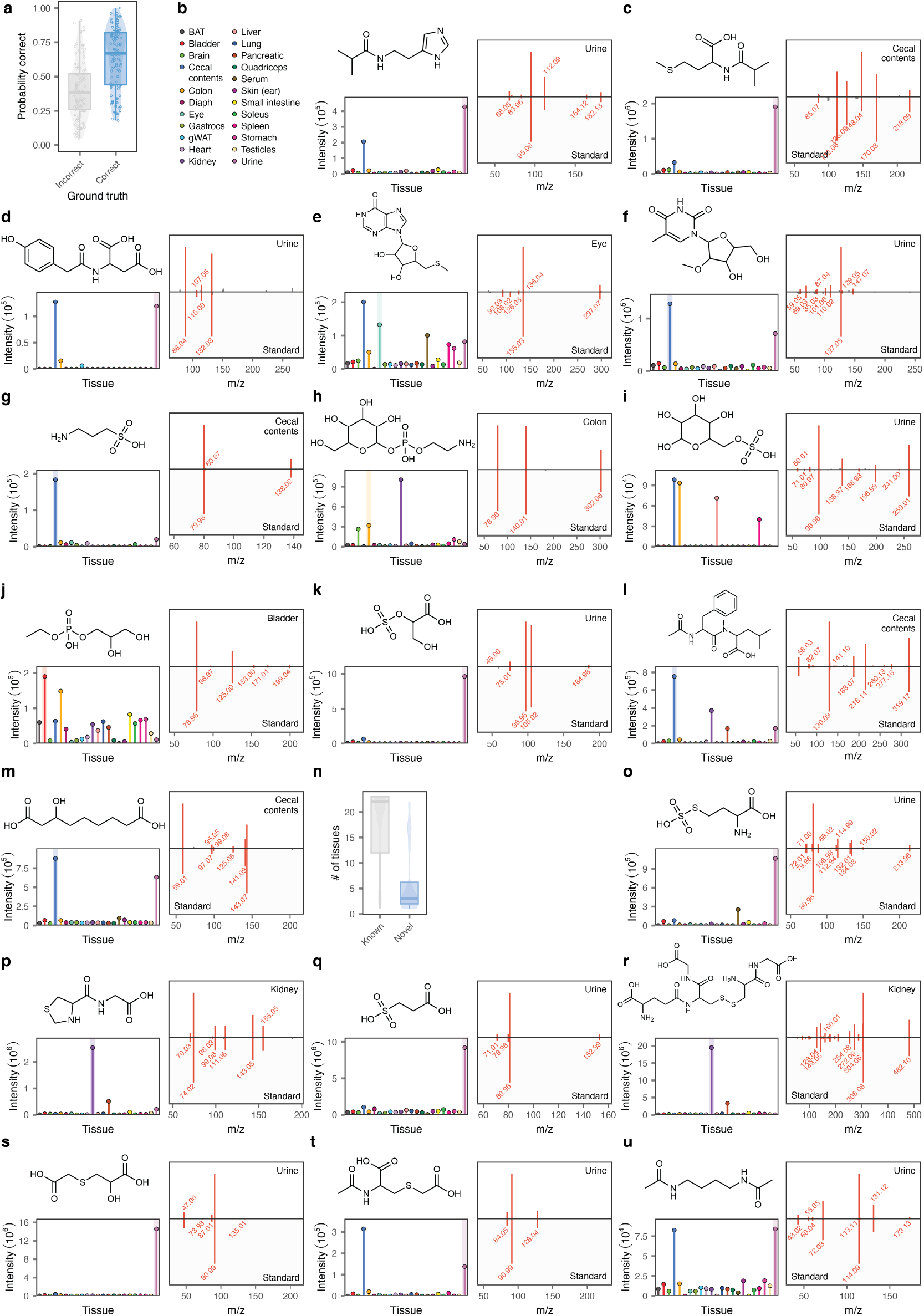
Discovery and characterization of additional mouse and human metabolites. **a**, Probability assigned to each *de novo* structure annotation for peaks representing held-out metabolites by the meta-learning model, shown separately for correct versus incorrect annotations. **b**, Left, structure of N-isobutyryl histamine (top) and MS1 intensity across 23 mouse tissues (bottom). Right, mirror plot showing the similarity between MS/MS spectra from the synthetic standard versus the experimental spectrum from mouse urine. **c**, As in **b**, but for N-isobutyryl methionine. **d**, As in **b**, but for (2-(4-hydroxyphenyl)acetyl)-aspartic acid. **e**, As in **b**, but for methylthioinosine. **f**, As in **b**, but for 2’-O-methyl-5-methyluridine. **g**, As in **b**, but for homotaurine. **h**, As in **b**, but for (2-aminoethyl)phosphate-1-hexopyranose. **i**, As in **b**, but for 6-O-sulfo-hexopyranose. **j**, As in **b**, but for glycerylphosphorylethanol. **k**, As in **b**, but for 2-sulfoglycerate. **l**, As in **b**, but for N-acetyl-phenylalanylleucine. **m**, As in **b**, but for 7-hydroxyazelaic acid. **n**, Tissue specificity of newly-discovered metabolites versus known metabolites identified by comparison to an in-house library of synthetic standards, as quantified by the number of tissues in which the metabolite in question was identified with a MS1 intensity of 10^5^ or greater. **o**, As in **b**, but for S-sulfohomocysteine. **p**, As in **b**, but for thioprolylglycine. **q**, As in **b**, but for 3-sulfopropanoic acid. **r**, As in **b**, but for glutathione-cysteinylglycine mixed disulfide. **s**, As in **b**, but for 3-carboxymethyl-thiolactic acid. **t**, As in **b**, but for N-acetyl-S-carboxymethyl-cysteine. **u**, As in **b**, but for diacetylputrescine.

**Supplementary Fig. 8.**
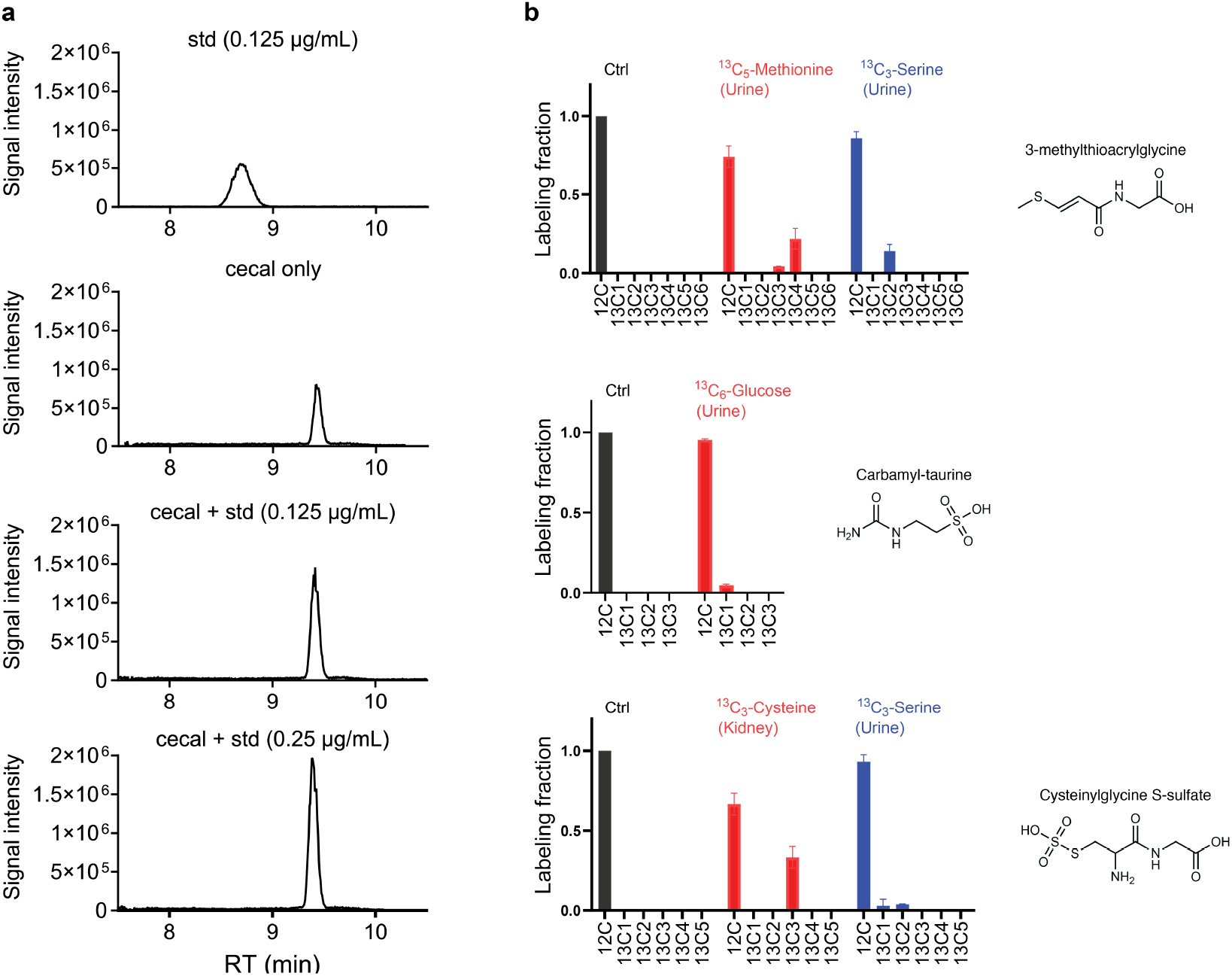
Supporting data for metabolite identification and characterization. **a**, Spiking N-carbamyl-taurine standard into cecal contents extract shows that the two features merge as one peak with the peak intensity increasing with the amount spiked in, confirming the feature detected in the cecal contents to be N-carbamyl-taurine. **b**, Isotope labeling patterns of metabolites shown in **Fig. 6**, after removing contributions from natural isotope abundance. The tracers and the tissues used are indicated.

**Supplementary Fig. 9.**
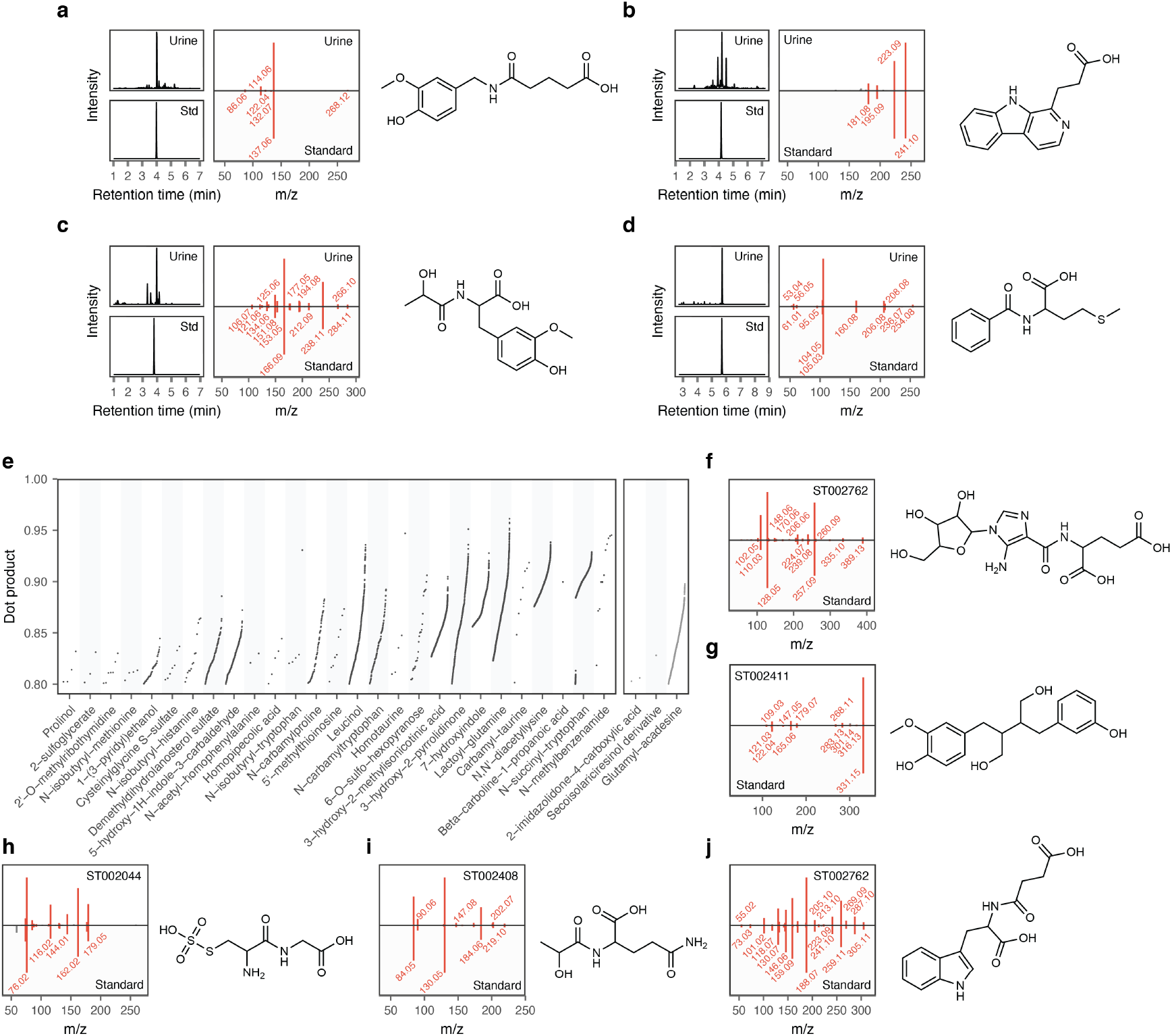
Additional metabolite discoveries. **a-d**, Additional novel human metabolites predicted to exist by DeepMet, and assigned to peaks in the mouse tissue dataset, that were subsequently experimentally identified in human urine. Left, extracted ion chromatograms from the chemical standard and a representative urine metabolome; middle, mirror plots showing the similarity between MS/MS spectra from the standard versus the experimental spectrum from human urine; right, chemical structures of the predicted metabolites. **a**, N-glutarylvanillylamine; **b**, *β*-carboline-1-propionic acid; **c**, N-lactoyl-3-methoxytyrosine; **d**, N-benzoylmethionine. **e**, Overview of novel metabolites tentatively identified in public repositories by comparisons to reference spectra acquired in this study. Points show matches between reference spectra and experimental MS/MS spectra from human metabolomics experiments in MetaboLights or Metabolomics Workbench, as quantified by the dot-product. Left, metabolites also identified through comparisons to chemical standards under identical LC-MS/MS conditions in human or mouse; right, metabolites tentatively identified only by MS/MS search in public repositories. **f-j**, Mirror plots showing similarities between MS/MS spectra from chemical standards and experimental spectra from public repositories. **f**, Glutamylacadesine; **g**, desmethoxy-secoisolariciresinol; **h**, S-sulfocysteinylglycine; **i**, N-lactoyl-glutamine; **j**, N-succinyl-tryptophan.

**Supplementary Fig. 10.**
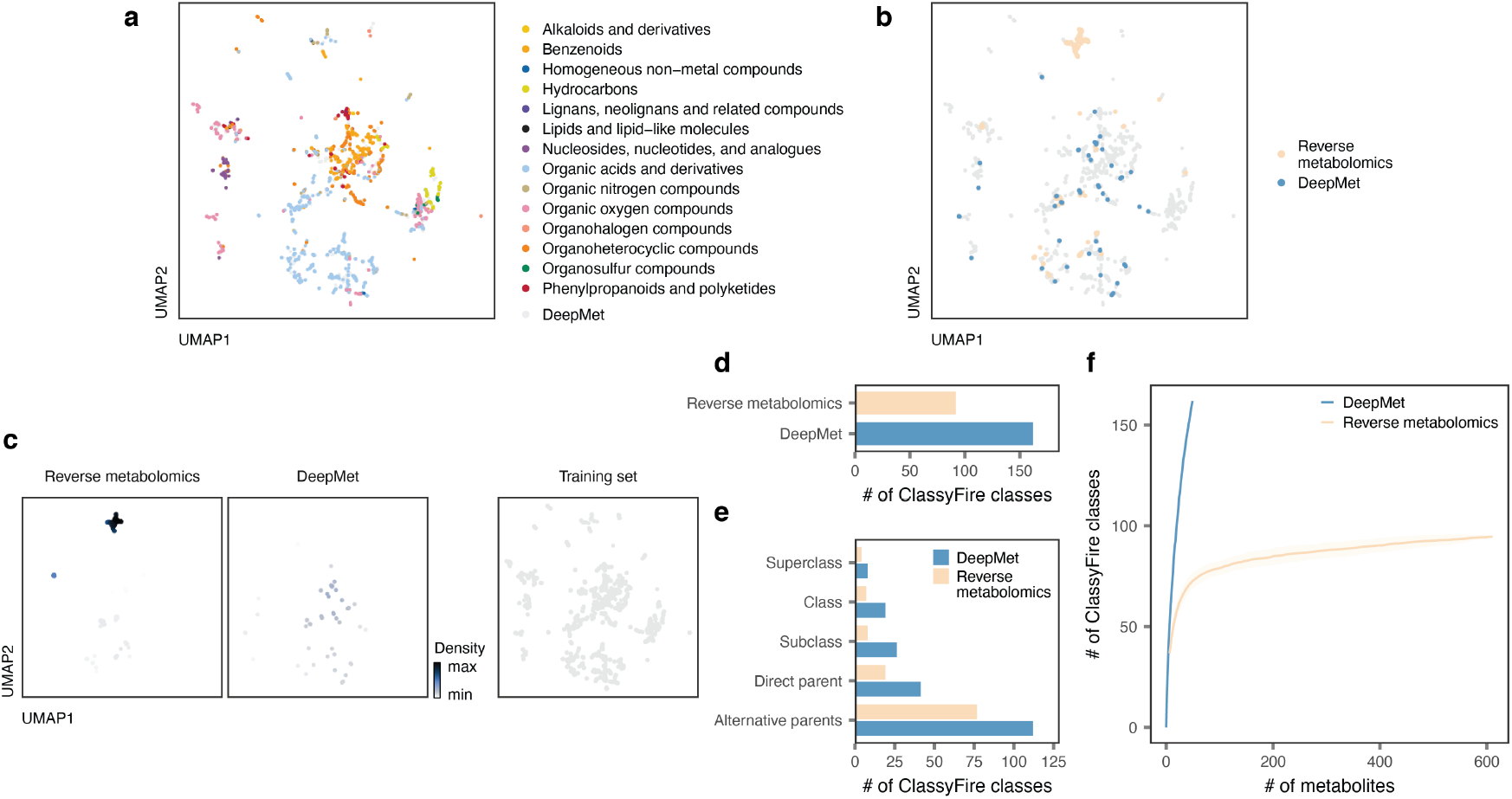
Structural diversity of metabolites discovered by DeepMet versus reverse metabolomics. **a**, UMAP visualization of the chemical space occupied by known metabolites and generated molecules, reproduced from **Fig. 1**. Known metabolites are superimposed over generated molecules and colored by their assigned superclasses in the ClassyFire chemical ontology. **b**, Positions of metabolites discovered by DeepMet versus reverse metabolomics within the same two-dimensional representation of chemical space shown in **a**, with known metabolites in grey. **c**, As in **b**, but showing the density of metabolites discovered by DeepMet versus reverse metabolomics within the same two-dimensional representation of chemical space shown in **a**, as compared to the training set (all in grey). **d**, Number of classes in the ChemOnt ontology assigned by ClassyFire to metabolites discovered by DeepMet versus reverse metabolomics. **e**, As in **d**, but showing each level of the ChemOnt hierarchy separately. **f**, Rarefaction curve showing the number of classes in the ChemOnt ontology assigned by ClassyFire to metabolites discovered by DeepMet versus reverse metabolomics, as a function of total number of metabolites sampled. Error bars show the standard deviation over 100 samples.

