## Supplementary Note 1 for "Language model-guided anticipation and discovery of unknown metabolites"

### Chemical synthesis

*N-lactoyl-glutamine and N-lactoyl-serine.*

The synthesis of these lactoyl-amino acids was performed using a solid phase method. Customized glassware, long needles, and magnets, along with a 250 mL round bottom flask, were heated at 130°C for 24 hours, followed by a 2-hour desiccation. Solvents were dried using 4 Å molecular sieves, with 10 g of sieves added to a flask containing 100 mL DMF and left at room temperature for 12 hours.

The synthesis was initiated by adding 2 grams of Fmoc-Amino Acid-Wang Resin to the prepared glassware. For capping of the resin, 40 mL of acetic anhydride, 6 mL of Hunig's Base, and 2 bits of 4-Dimethylaminopyridine (DMAP) were added to block the unreacted sites. To prevent contamination and the reaction with atmospheric oxygen, the mixture was stirred under a nitrogen atmosphere.

Following the capping, 20% piperidine solution in dry N,N-dimethylformamide (DMF) was used to remove the Fmoc protecting groups. To ensure that the resin was thoroughly cleaned, and all unreacted materials were removed, multiple washing steps were performed with dichloromethane (DCM) and DMF.

The core amide coupling reaction was facilitated by adding 1.5 equivalents of HATU, 3.6 equivalents of Hunig's Base, and 1.2 equivalents of THP-protected lactic acid, dissolved in 100 mL of DCM, to the resin. This step was also conducted under a nitrogen atmosphere to maintain an inert environment.

After the coupling reaction, the product was deprotected and cleaved from the resin using a cleavage cocktail consisting of 9:1:1 trifluoroacetic acid (TFA), water, and triisopropylsilane. This mixture was stirred, allowing the cleavage of the product from the resin. The cleaved product was then extracted using a series of washes with diethyl ether and water, isolating the aqueous phase containing the target compound.

In a fume hood, the lactoyl amino acid-containing aqueous extract was dried overnight under airflow. Afterward, the dried product was characterized by nuclear magnetic resonance (NMR) and mass spectrometry to confirm its purity and structure. A sample dissolved in $D_2O$ was used to obtain the NMR spectra.

*N-carbamyl-taurine (2-ureidoethanesulfonic acid).*

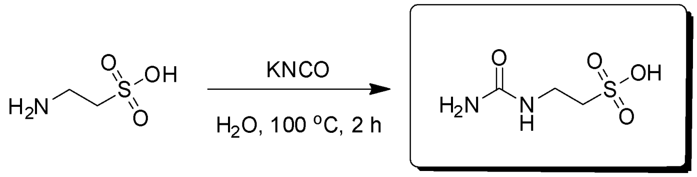


**Step 1** **HKSP-0001**

To a solution of 2-aminoethanesulfonic acid (500 mg, 4.00 mmol, 498.01 µL, 1 eq) in $H_2O$ (1 mL) was added potassium cyanate (324.08 mg, 4.00 mmol, 157.63 µL, 1 eq). The mixture was stirred at 100°C for 2 h. The reaction mixture was concentrated under reduced pressure. The crude was recrystallized by dissolving it in 1 mL of water and adding 5 mL of absolute ethanol slowly with stirring. After 1 h, the resulting white crystals were collected by filtration, washed on the filter with absolute ethanol, and dried in a vacuum desiccator. The free acid was obtained by dissolving this quantity of potassium salt in water (1.5 mL) and acidifying with concentrated hydrochloric acid (0.25 mL), then inducing crystallization by slow addition of absolute ethanol (4 mL) with stirring. The white crystals were filtered from the mixture and washed on the filter with absolute ethanol. Compound 2-ureidoethanesulfonic acid (105.8 mg, 629.12 µmol, 15.75% yield, 100% purity) was obtained as a white solid.

**LCMS:** HKSP-0001 (M+1): 169.0@ 0.236min (0-30% ACN in $H_2O$, 6 min)

**$^1$H NMR:** HKSP-0001 (400 MHz, DMSO-d6)

δ 3.46 (t, *J* = 6.6 Hz, 2H), 3.03 (t, *J* = 6.6 Hz, 2H).

*S-sulfohomocysteine.*

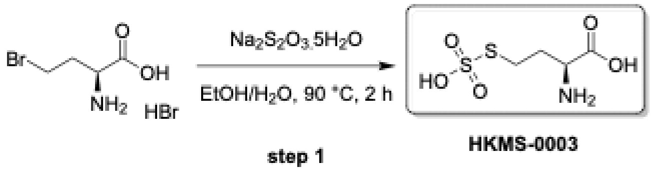


To a solution of (2S)-2-amino-4-bromo-butanoic acid (0.2 g, 760.67 μmol, 1 eq, HBr) in EtOH (0.5 mL) was added $Na_2S_2O_3 \cdot 5H_2O$ (377.56 mg, 1.52 mmol, 373.83 μL, 2 eq) in $H_2O$ (0.5 mL). The mixture was stirred at 90°C for 4 h. It was filtered and concentrated under reduced pressure to give a residue. The residue was purified by prep-HPLC (column: HUAPU 1010-X Amide 100 × 30 mm ×10 μm; mobile phase: [$H_2O$(0.1%TFA)-ACN]; gradient: 90%-70% B over 15.0 min) to afford the compound (7.9 mg, 36.70 μmol, 4.82% yield, 100% purity) as a white solid.
**LCMS:** HKMS-0003 (M-$H^-$):213.9 @ 3.811 min (90-60 % ACN in $H_2O$, 15 min)
**$^1$H NMR:** HKMS-0003 (400 MHz, D2O)
δ = 4.10 (t, *J* = 6.5 Hz, 1H), 3.18 (t, *J* = 7.4 Hz, 2H), 2.50 - 2.21 (m, 2H)

*Glutathione-cysteinylglycine mixed disulfide.*

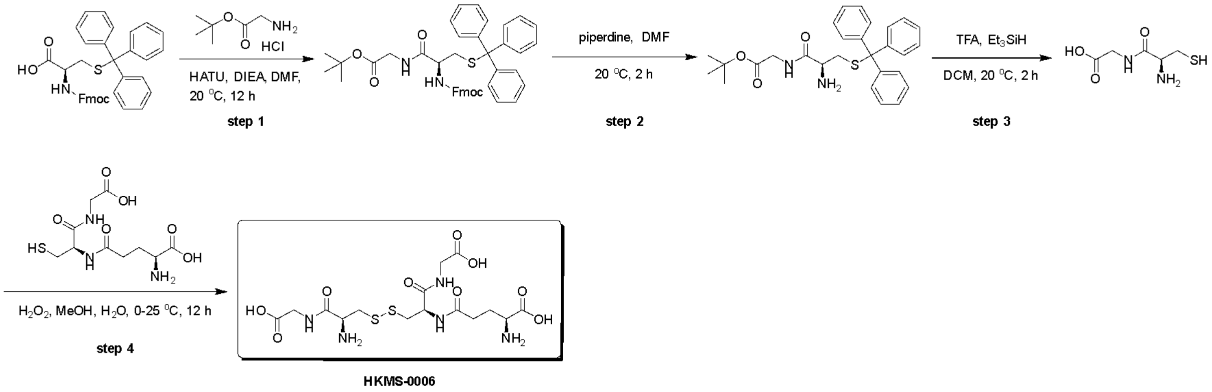


Step 1: Synthesis of *tert*-butyl N-(((9H-fluoren-9-yl)methoxy)carbonyl)-S-trityl-D-cysteinylglycinate

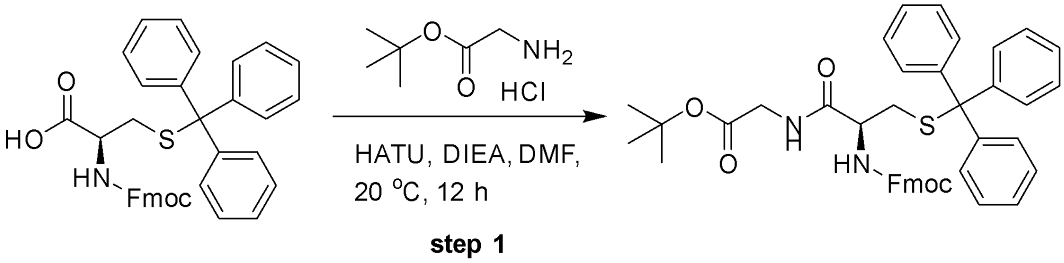


To a solution of (2R)-2-(9H-fluoren-9-ylmethoxycarbonylamino)-3-tritylsulfanyl-propanoic acid (5.61 g, 9.57 mmol, 1.07 eq) in DMF (25 mL) was added tert-butyl 2-aminoacetate (1.5 g, 8.95 mmol, 1.0 eq, HCl) and HATU (3.64 g, 9.57 mmol, 1.07 eq) at 20°C. The mixture was stirred at 20°C for 5 min. DIEA (2.31 g, 17.90 mmol, 3.12 mL, 2.0 eq) was added into the reaction at 20°C. The mixture was stirred at 20°C for 11.9 hr. The reaction was quenched with $H_2O$ (80 mL) and extracted with EtOAc (80 mL ×3). The combined organic layers were washed with brine (80 mL ×3), dried over $Na_2SO_4$, filtered and concentrated under reduced pressure to give a residue. The residue was purified by flash silica gel chromatography (ISCO®; 80 g SepaFlash® Silica Flash Column, eluent of 15~20% ethylacetate/petroleum ether gradient @ 120 mL/min) to give the compound (6.2 g) as colorless oil.

Step 2: Synthesis of *tert*-butyl S-trityl-D-cysteinylglycinate

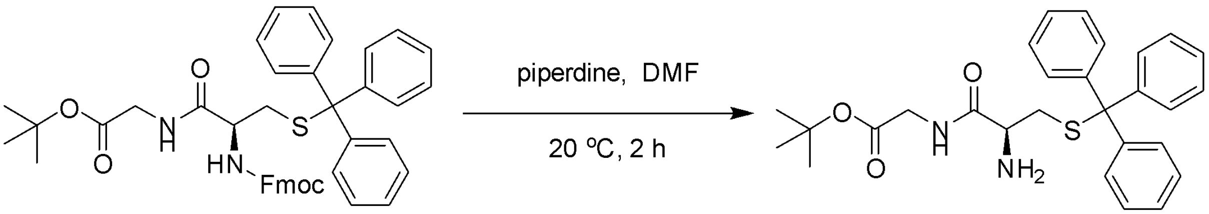


**step 2**

To a solution of *tert*-butyl 2-[[(2R)-2-(9H-fluoren-9-ylmethoxycarbonylamino)-3-tritylsulfanyl-propanoyl]amino]acetate (3.1 g, 4.44 mmol, 1 eq) in DMF (12.5 mL) was added piperidine (2.64 g, 31.05 mmol, 3.07 mL, 7 eq). The mixture was stirred at 20°C for 2 h. Two parallel reactions were worked up together. The reaction mixture was concentrated under reduced pressure to remove DMF. The residue was diluted with $H_2O$ (50 mL) and extracted with EtOAc (50 mL ×3). The combined organic layers were washed with brine (50 mL ×3), dried over $Na_2SO_4$, filtered and concentrated under reduced pressure to give a residue. The residue was purified by flash silica gel chromatography (ISCO®; 80 g SepaFlash® Silica Flash Column, eluent of 75-85% ethyl acetate/petroleum ether @ 100 mL/min) to give the compound (4.0 g, 8.27 mmol, 93.25% yield, 98.58% purity) as a colorless oil.

Step 3: Synthesis of D-cysteinylglycine

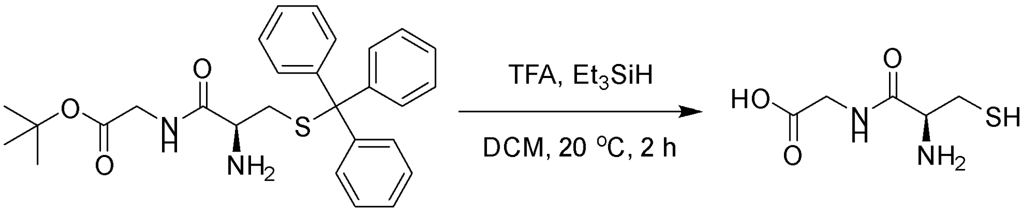


**step 3**

To a solution of tert-butyl 2-[[(2S)-2-amino-3-tritylsulfanyl-propanoyl]amino]acetate (0.2 g, 419.61 μmol, 1 eq) in DCM (12 mL) was added TFA (47.85 mg, 419.61 μmol, 31.17 μL, 1 eq) and $Et_3SiH$ (107.34 mg, 923.15 μmol, 147.45 μL, 2.2 eq). The mixture was stirred at 20°C for 2 h. The reaction mixture was poured into $H_2O$ (10 mL), and extracted with EtOAc (10 mL ×3). The water phase was lyophilized to give the compound (0.07 g, 392.80 μmol, 93.61% yield, 97% purity) as a white solid.

Step 4: Synthesis of N5-((R)-3-(((S)-2-amino-3-((carboxymethyl)amino)-3-oxopropyl)disulfaneyl)-1-((carboxymethyl)amino)-1-oxopropan-2-yl)-L-glutamine

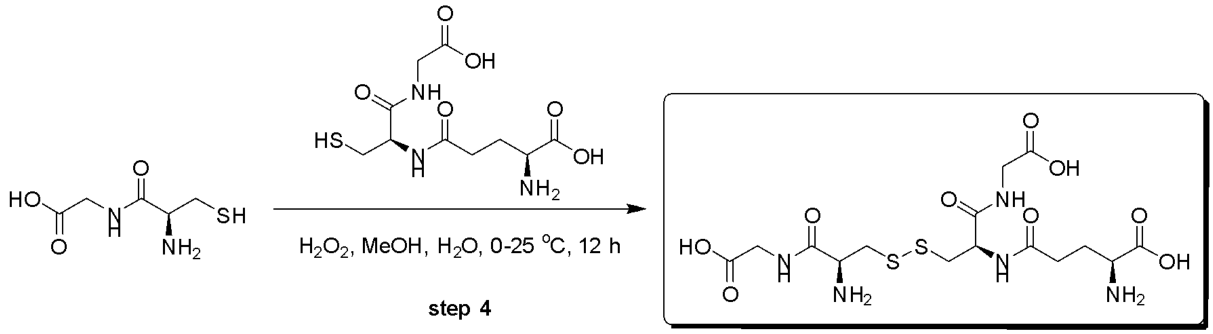

**HKMS-0006**

To a solution of 2-[[(2S)-2-amino-3-sulfanyl-propanoyl]amino]acetic acid (0.05 g, 280.57 µmol, 1 eq) in MeOH (1.5 mL) and to a solution of (2S)-2-amino-5-[[(1R)-2-(carboxymethylamino)-2-oxo-1-(sulfanylmethyl)ethyl]amino]-5-oxo-pentanoic acid (68.98 mg, 224.46 µmol, 0.8 eq) in $H_2O$ (1.5 mL) was added $H_2O_2$ (47.72 mg, 420.85 µmol, 40.44 µL, 30% purity, 1.5 eq) at 0°C. The mixture was stirred at 20°C for 12 h after which the mixture was filtered and concentrated under a stream of $N_2$ where MeOH was evaporated. The concentrated residue was purified by prep-HPLC (column: Ultimate Polar RP C18 100 × 30 mm × 10 µm; mobile phase: [$H_2O$(0.1%TFA)-ACN];B%:1%, isocratic elution mode) to give the compound (22.2 mg, 42.75 µmol, 15.24% yield, 93.1% purity) as a white solid.
**LCMS:** HKMS-0006 ($M+H^+$):484.0 @ 4.199 min (0-30 % ACN in $H_2O$, 10 min)
**$^1$H NMR:** HKMS-0006 (400 MHz, D2O)
δ = 4.73 - 4.72 (m, 1H), 4.30 (t, *J* = 6.4 Hz, 1H), 4.01 - 3.89 (m, 4H), 3.78 (t, *J* = 6.4 Hz, 1H), 3.23 (s, 3H), 3.01 (br d, *J* = 9.1 Hz, 1H), 2.56 - 2.40 (m, 2H), 2.17 - 2.03 (m, 2H)

*[(2R,3S,4S,5R,6R)-3,4,5,6-tetrahydroxytetrahydropyran-2-yl]methyl hydrogen sulfate.*

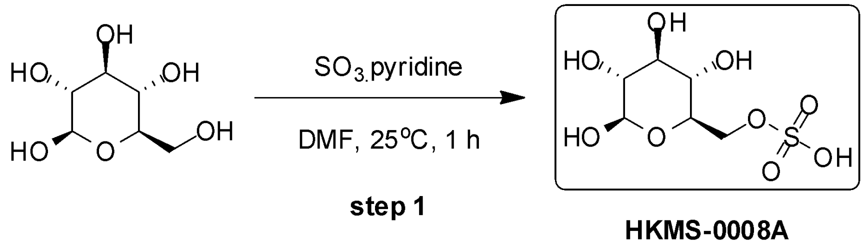

**HKMS-0008A**

To a solution of pyridine-sulfur trioxide (441.74 mg, 2.78 mmol, 1.0 eq) in DMF (2.5 mL) was added the solution of (2R,3R,4S,5S,6R)-6-(hydroxymethyl)tetrahydropyran-2,3,4,5-tetrol (500 mg, 2.78 mmol, 1.0 eq) in DMF (2.5 mL). The mixture was stirred at 25°C for 1 h. The suspension was purified by prep-HPLC ($NH_4HCO_3$ condition, column: Kromasil C18(W) 150 × 30 × 10; mobile phase: [$H_2O$ (10mM $NH_4HCO_3$)-ACN]; B%:1%, isocratic elution mode) to give [(2R,3S,4S,5R,6R)-3,4,5,6-tetrahydroxytetrahydropyran-2-yl]methyl hydrogen sulfate (133 mg, 478.24 µmol, 17.23% yield, 93.57% purity) as a white solid.
**LCMS:** HKMS-0008A (M-H):259.0@ 4.783min (90-50CD_10_HILIC_Amide, 10 min)
**$^1$H NMR:** HKMS-0008A (400 MHz, DMSO-d6)
δ 5.13-4.74 (m, 1H), 4.70-4.21 (m, 1H), 4.13-3.55 (m, 3H), 3.54-3.39 (m, 4H), 3.25-2.76 (m, 2H).

*Synthesis of 2,3-dihydroxypropyl ethyl hydrogen phosphate*

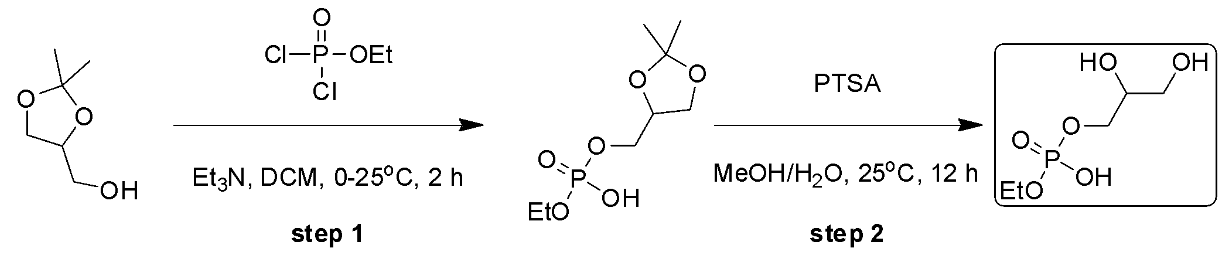


**HKMS-0016**

Step 1: Synthesis of (2,2-dimethyl-1,3-dioxolan-4-yl)methyl ethyl hydrogen phosphate

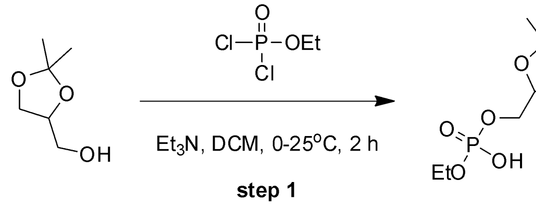


To a solution of (2,2-dimethyl-1,3-dioxolan-4-yl)methanol (300 mg, 2.27 mmol, 281.43 µL, 1.0 eq) in DCM (9 mL) was added $Et_3N$ (574.25 mg, 5.68 mmol, 789.89 µL, 2.5 eq) and 1-dichlorophosphoryloxyethane (554.81 mg, 3.41 mmol, 404.09 µL, 1.5 eq) at 0°C. The mixture was stirred at 25°C for 2 h. The mixture was adjusted to pH 7-8 with saturated aqueous $NaHCO_3$ (20 mL). The mixture was extracted with $CH_2Cl_2$ (20 mL ×3). The combined organic layers were washed with brine (20 mL), dried over anhydrous $Na_2SO_4$, filtered and concentrated under reduced pressure to give a crude product. The crude product (2,2-dimethyl-1,3-dioxolan-4-yl)methyl ethyl hydrogen phosphate (550 mg, crude) was obtained as yellow oil. The crude product was used for the next step without further purification.

Step 2: 2,3-dihydroxypropyl ethyl hydrogen phosphate

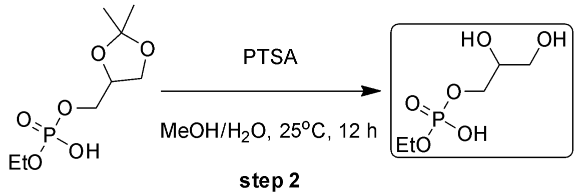


**HKMS-0016**

To a mixture of (2,2-dimethyl-1,3-dioxolan-4-yl)methyl ethyl hydrogen phosphate (150 mg, 624.51 µmol, 1 *eq*) in MeOH (0.5 mL) and $H_2O$ (0.5 mL) was added PTSA (107.54 mg, 624.51 µmol, 1 *eq*) at 25°C. The mixture was stirred at 25°C for 12 h. The reaction mixture was concentrated to remove MeOH. The residue was purified by prep-HPLC ($NH_4HCO_3$ condition; column: Kromasil C18(W) 150 × 30 × 10; mobile phase: [$H_2O$(10mM $NH_4HCO_3$)-ACN]; gradient:1%-20% B over 12.0 min) to give 2,3-dihydroxypropyl ethyl hydrogen phosphate (1.8 mg, 8.99 µmol, 1.44% yield, 99.90% purity) as a white solid.
**LCMS:** HKMS-0016 (M+1):201.0 @ 0.422min (0-30CD-ELSD, 6 min)
**$^1$H NMR:** HKMS-0016 (400 MHz, D2O)
δ 3.94-3.74 (m, 5H), 3.66-3.52 (m, 2H), 1.21 (t, *J* = 7.0 Hz, 3H).

*Synthesis of 3-hydroxynonanedioic acid.*

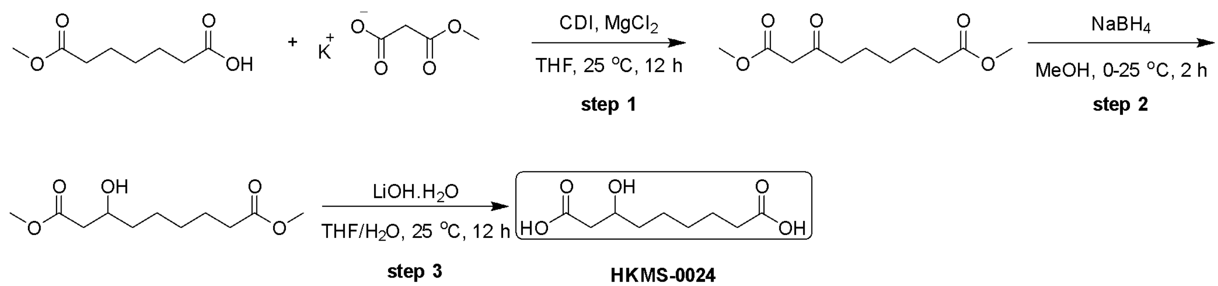


**HKMS-0024**

Step 1: Synthesis of dimethyl 3-oxononanedioate

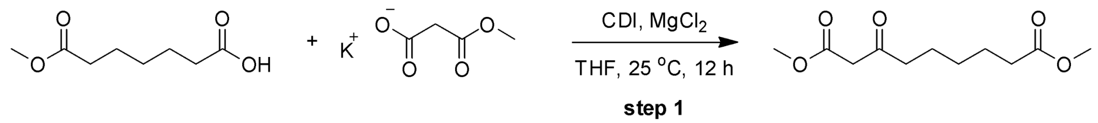


To a solution of 7-methoxy-7-oxo-heptanoic acid (1.0 g, 5.74 mmol, 1.0 eq) in THF (10 mL) was added CDI (1.02 g,6.31 mmol, 1.1 eq) at 25°C. The mixture was stirred at 25°C for 1 h. Then potassium;3-methoxy-3-oxo-propanoate (1.08 g, 6.89 mmol, 1.2 eq) and $MgCl_2$ (819.87 mg, 8.61 mmol, 353.39 µL, 1.5 eq) was added to the mixture. The mixture was stirred at 25°C for 11 h. The reaction mixture was acidized with 1M HCl (5 mL) to pH = 7 at 0ºC. The mixture was poured into $H_2O$ (20 mL) and extracted with EtOAc (10 mL ×3). The combined organic layers were washed with brine (20 mL), dried over anhydrous $Na_2SO_4$, filtered and concentrated under reduced pressure to give dimethyl 3-oxononanedioate (1.4 g, crude) was obtained as yellow oil. The crude product was used for the next step without further purification.

**$^1$H NMR:** (400 MHz, CHLOROFORM-d)

δ 3.73 (s, 3H), 3.66 (s, 3H), 3.44 (s, 2H), 2.54 (t, *J* = 7.3 Hz, 2H), 2.30 (t, *J* = 7.5 Hz, 2H), 1.66-1.58 (m, 4H), 1.37-1.29 (m, 2H)

Step 2: Synthesis of dimethyl 3-hydroxynonanedioate

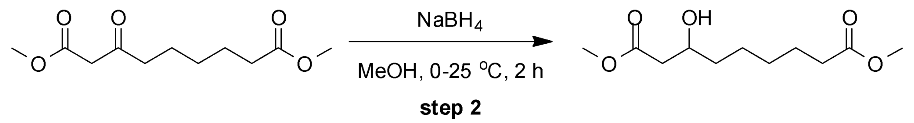


A solution of dimethyl 3-oxononanedioate (600 mg, 2.61 mmol, 1 eq) in MeOH (6 mL) was treated portion-wise with $NaBH_4$ (147.87 mg, 3.91 mmol, 1.5 eq) at 0°C. The mixture was stirred at 25°C for 2 h. The reaction mixture was concentrated to remove MeOH to give the residue. The residue was dissolved into saturated aqueous $NH_4Cl$ solution (10 mL) and then extracted with EtOAc (20 mL ×3). The combined organic layers were concentrated under reduced pressure to give dimethyl 3-hydroxynonanedioate (500 mg, crude) as colorless oil. The crude product was used for the next step without further purification.

Step 3: Synthesis of 3-hydroxynonanedioic acid

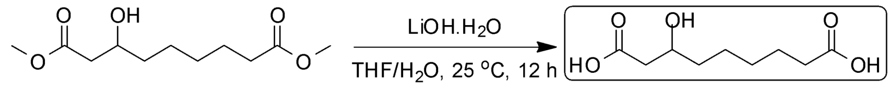


**HKMS-0024**

To a solution of dimethyl 3-hydroxynonanedioate (500 mg, 2.15 mmol, 1 eq) in THF (6 mL) and $H_2O$ (2 mL) was added LiOH.$H_2O$ (271 mg, 6.46 mmol, 3 eq) at 25°C. The mixture was stirred at 25°C for 12 h. The suspension was concentrated under reduced pressure to remove THF. The mixture was purified by

prep-HPLC (HCl condition, column: Phenomenex Luna C18 100 × 30 mm × 3 μm; mobile phase: [$H_2O$(0.04%HCl)-ACN];gradient:1%-40% B over 12.0 min) to give 3-hydroxynonanedioic acid (180 mg, 880.96 μmol, 40.92% yield, 99.95% purity) as a white solid.
**LCMS:** HKMS-0024 (M+1):205.1 @ 2.302min (0-30AB-ELSD, 6 min)
**$^1$H NMR:** HKMS-0024 (400 MHz, DEUTERIUM OXIDE)
δ 4.06-3.95 (m, 1H), 2.62-2.51 (m, 1H), 2.46-2.37 (m, 1H), 2.34 (t, *J* = 7.4 Hz, 2H), 1.60-1.52 (m, 2H), 1.51-1.44 (m, 2H), 1.40-1.24 (m, 4H).

*Synthesis of* (S)-3-hydroxy-2-(sulfooxy)propanoic acid*.*

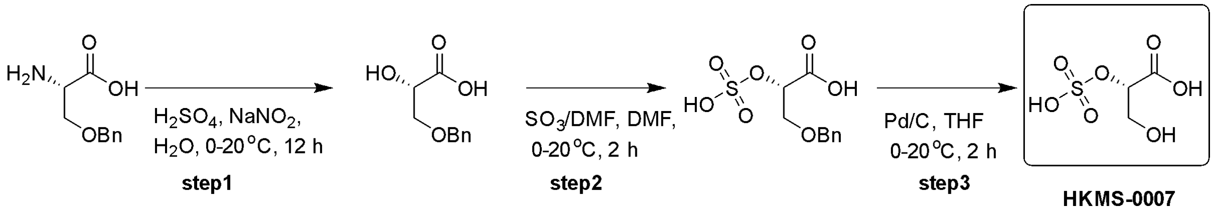


Step 1: synthesis of (R)-3-(benzyloxy)-2-hydroxypropanoic acid
To a mixture of O-benzyl-D-serine (5 g, 25.61 mmol, 1 eq) in $H_2SO_4$ (2 M, 30.74 mL, 2.4 eq) and $H_2O$ (20 mL) at 0°C was added $NaNO_2$ (2.83 g, 40.98 mmol, 1.6 eq). The mixture was stirred for 12 h at 20°C. The reaction mixture was poured into $H_2O$ (50 mL), and extracted with EtOAc (50 mL ×3). The extracted aqueous phase was spin dried to obtain the product. Compound (R)-3-(benzyloxy)-2-hydroxypropanoic acid (10 g, crude) was obtained as a white solid, and used directly in the next step without further purification.
**$^1$H NMR:** (400 MHz, DEUTERIUM OXIDE)
δ 7.49 - 7.31 (m, 5H), 4.60 - 4.53 (m, 2H), 3.91 - 3.72 (m, 2H), 3.71 - 3.62 (m, 1H)

Step 2: synthesis (R)-3-(benzyloxy)-2-(sulfooxy)propanoic acid
To a mixture of (R)-3-(benzyloxy)-2-hydroxypropanoic acid (1.00 g, 5.10 mmol, 1 eq) in DMF (15 mL) at 0°C was added N,N-dimethylformamide;sulfur trioxide (3.12 g, 20.39 mmol, 4 eq), The mixture was stirred for 0.5 h at 20°C. The reaction mixture was poured into $KHCO_3$ (aq 20 mL), and extracted with EtOAc (20 mL ×3). The extracted aqueous phase was spin dried to obtain the product. The residue was purified by prep-HPLC (column: Waters Xbridge BEH C18 250 × 50 mm × 10 μm; mobile phase: [$H_2O$(10mM $NH_4HCO_3$)-ACN]; gradient: 1%-15% B over 10.0 min) to afford (R)-3-(benzyloxy)-2-(sulfooxy)propanoic acid (25 mg, 98% purity) as white solid.

Step 3: synthesis of (S)-3-hydroxy-2-(sulfooxy)propanoic acid
To a solution of (S)-3-(benzyloxy)-2-(sulfooxy)propanoic acid (25 mg, 90.49 μmol, 1 eq) in THF (2 mL) was added Pd/C (25.00 mg, 23.49 μmol, 10% purity, 0.26 eq) under $N_2$ atmosphere. The suspension was degassed and purged 3 times with $H_2$. The mixture was stirred under $H_2$ (15 psi) at 20°C for 2 h. It was filtered and concentrated under reduced pressure to give (S)-3-hydroxy-2-(sulfooxy)propanoic acid (19.6 mg, 86.67 μmol, 95.78% yield, 100% purity) as a white solid.

**LCMS:** (M+H-):185.0 @ 2.005 min (5-95 % ACN in $H_2O$, 6 min)
**1H NMR:** (400 MHz, DEUTERIUM OXIDE)
δ 4.59 (dd, J = 3.0, 5.9 Hz, 1H), 3.92 - 3.86 (m, 1H), 3.84 - 3.76 (m, 1H)

*Synthesis of 2-aminoethyl [(2S, 3R, 4S, 5S, 6R)-3, 4, 5-trihydroxy-6-(hydroxymethyl) tetrahydropyran-2-yl] hydrogen phosphate.*

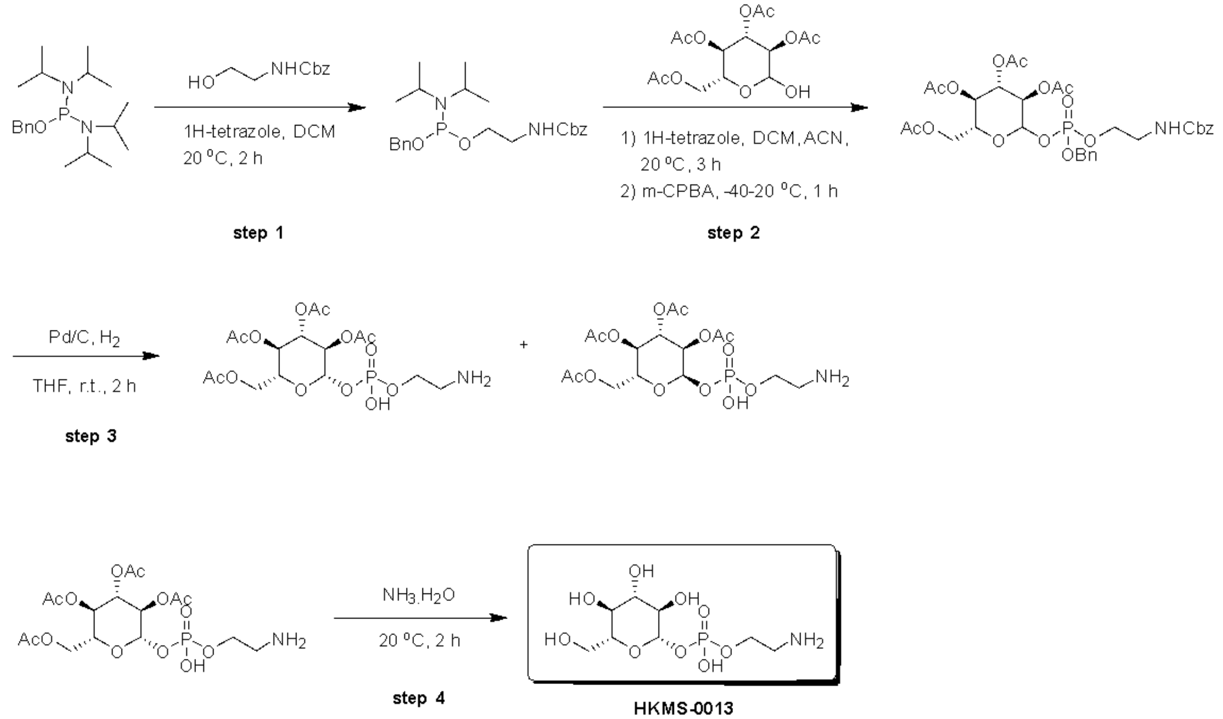


HKMS-0013

Step 1: synthesis of benzyl (2-(((benzyloxy) (diisopropylamino) phosphaneyl) oxy) ethyl) carbamate

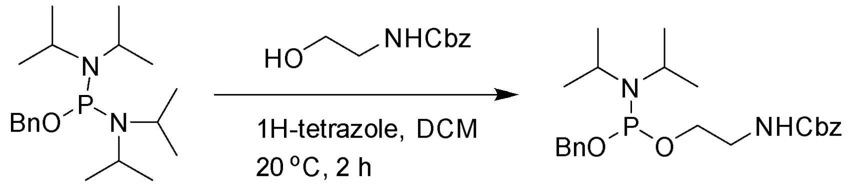


To a solution of N-[benzyloxy-(diisopropylamino)phosphanyl]-N-isopropyl-propan-2-amine (0.25 g, 738.63 μmol, 1 eq) and benzyl N-(2-hydroxyethyl)carbamate (129.77 mg, 664.76 μmol, 0.9 eq) in DCM (2 mL) was added 2H-tetrazole (25.87 mg, 369.31 μmol, 32.75 μL, 0.5 eq). The mixture was stirred at 20°C for 3h. The solution of benzyl N-[2-[benzyloxy-(diisopropylamino) phosphanyl] oxyethyl] carbamate (4 mL) was used in the next step without further purification.

Step 2: synthesis of (2R, 3R, 4S, 5R)-2-(acetoxymethyl)-6-(((benzyloxy) (2-(((benzyloxy) carbonyl) amino)ethoxy)phosphoryl)oxy)tetrahydro-2H-pyran-3,4,5-triyl triacetate

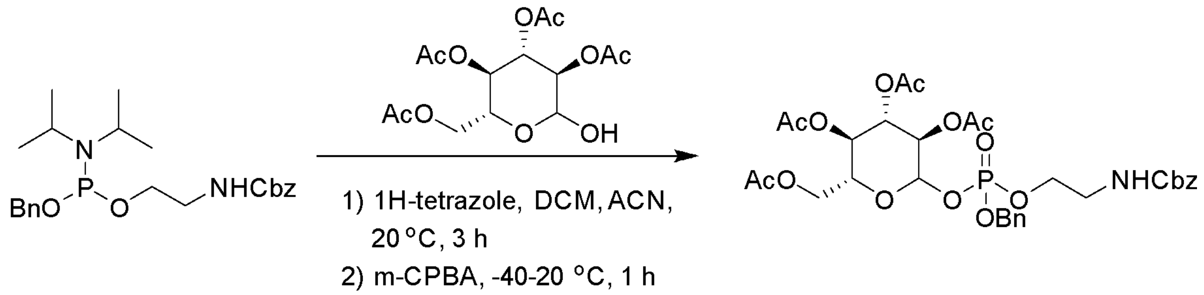

**step 2**

To a solution of [(2R,3R,4S,5R)-3,4,5-triacetoxy-6-hydroxy-tetrahydropyran-2-yl]methyl acetate (0.19 g, 545.50 µmol, 1 eq) in DCM (3 mL) was added 2H-tetrazole (114.64 mg, 1.64 mmol, 145.11 µL, 3 eq) in DCM (1 mL), the reaction mixture was stirred for 5 min, then was added benzyl N-[2-[benzyloxy (diisopropylamino)phosphanyl]oxyethyl]carbamate (306.70 mg, 709.16 µmol, 2 mL, 1.3 eq). The mixture was stirred at 20°C for 2 h. After cooling to –40°C, then was added MCPBA (469.77 mg, 1.63 mmol, 60% purity, 3 eq) in DCM (3 mL), and the mixture was stirred at 20°C for 1 h. The reaction mixture was quenched with $NaHCO_3$ (aq., 3 mL) and water (3 mL). The mixture was extracted with EtOAc (3 mL ×2) and the combined extracts were dried over $Na_2SO_4$, filtered and concentrated under reduced pressure to give a residue. The residue was purified by column chromatography ($SiO_2$, Petroleum ether/Ethyl acetate=1/0 to 1/1) to afford [(2R, 3R, 4S, 5R)-3, 4, 5-triacetoxy-6-[benzyloxy-[2-(benzyloxycarbonylamino) ethoxy] phosphoryl] oxy-tetrahydropyran-2-yl] methyl acetate (0.3 g, 431.28 µmol, 39.61% yield) as a yellow oil.

Step 3: synthesis of (2R, 3R, 4S, 5R, 6S)-2-(acetoxymethyl)-6-(((2-aminoethoxy) (hydroxy) phosphoryl) oxy) tetrahydro-2H-pyran-3, 4, 5-triyl triacetate

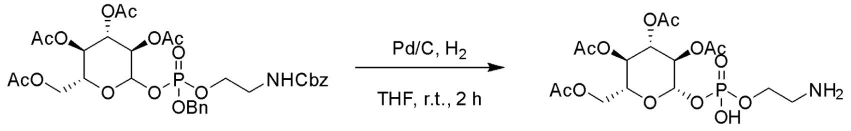

**step 3**

To a solution of [(2R, 3R, 4S, 5R)-3, 4, 5-triacetoxy-6-[benzyloxy-[2-(benzyloxycarbonylamino) ethoxy] phosphoryl] oxy-tetrahydropyran-2-yl] methyl acetate (300.00 mg, 431.28 µmol, 1 eq) in EtOAc (3 mL) was added Pd/C (0.1 g, 93.97 µmol, 10% purity, 2.18e-1 eq) under $N_2$ atmosphere. The suspension was degassed and purged with $H_2$ for 3 times. The mixture was stirred under $H_2$ (15 Psi) at 20°C for 12 h. It was filtered and concentrated under reduced pressure to give a residue. The residue was purified by prep-HPLC (column: Waters Xbridge BEH C18 100 × 30 mm × 10 µm; mobile phase: [$H_2O$ (10mM $NH_4HCO_3$)-ACN]; gradient: 1%-25% B over 12.0 min) to afford [(2R, 3R, 4S, 5R, 6S)-3, 4, 5-triacetoxy-6-[2-aminoethoxy(hydroxy) phosphoryl] oxy-tetrahydropyran-2-yl]methyl acetate (50 mg, 106.08 µmol, 24.60% yield) as a white solid, and [(2R, 3R, 4S, 5R, 6R)-3, 4, 5-triacetoxy-6-[2-aminoethoxy(hydroxy) phosphoryl] oxy-tetrahydropyran-2-yl]methyl acetate (11 mg, 23.34 µmol, 5.41% yield) as a white solid.

[(2R, 3R, 4S, 5R, 6S)-3, 4, 5-triacetoxy-6-[2-aminoethoxy(hydroxy) phosphoryl] oxy-tetrahydropyran-2-yl]methyl acetate:

**$^1$H NMR** (400 MHz, DMSO-$d_6$)

δ = 8.18 (br s, 2H), 5.36 - 5.15 (m, 3H), 4.91 (br t, *J* = 9.8 Hz, 1H), 4.76 (br dd, *J* = 8.0, 9.8 Hz, 1H), 4.19 -

4.14 (m, 2H), 4.05 - 3.98 (m, 3H), 3.81 (br dd, $J$ = 3.7, 8.3 Hz, 2H), 2.98 - 2.92 (m, 2H), 2.08 - 1.90 (m, 12H)

Step 4: synthesis of 2-aminoethyl ((2S, 3R, 4S, 5S, 6R)-3, 4, 5-trihydroxy-6-(hydroxymethyl) tetrahydro-2H-pyran-2-yl) hydrogen phosphate

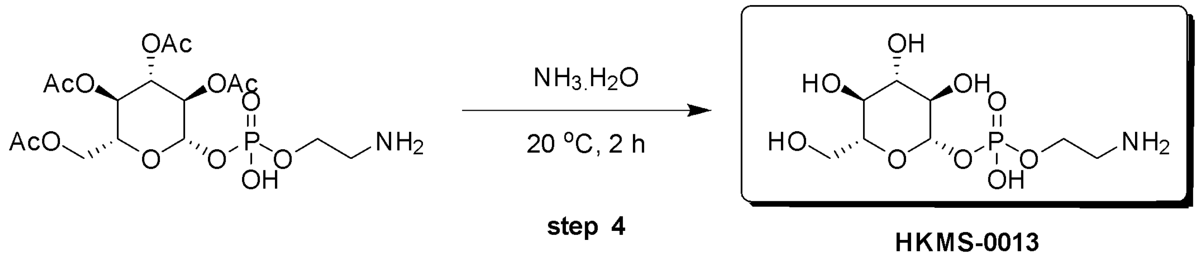

**HKMS-0013**

Solution of [(2R, 3R, 4S, 5R, 6S)-3, 4, 5-triacetoxy-6-[2-aminoethoxy (hydroxy) phosphoryl] oxy-tetrahydropyran-2-yl] methyl acetate (0.04 g, 84.86 µmol, 1 eq) in $NH_3$•$H_2O$ (1 mL) was stirred at 20°C for 2 h. The reaction mixture was concentrated under reduced pressure to give a residue. The residue was purified by prep-HPLC (column: HUAPU 1010-X Amide 100 × 30 mm × 10 µm; mobile phase: [$H_2O$ (10mM $NH_4HCO_3$)-ACN]; gradient: 90%-50% B over 15.0 min) to afford 2-aminoethyl [(2S, 3R, 4S, 5S, 6R)-3, 4, 5-trihydroxy-6-(hydroxymethyl) tetrahydropyran-2-yl] hydrogen phosphate (13.4 mg, 44.04 µmol, 51.89% yield, 99.64% purity) was obtained as a white solid.
**LCMS:** ($M+H^+$):304.2 @ 5.844 min (90_50 % ACN in $H_2O$, 10 min)
**$^1$H NMR:** (400 MHz, METHANOL-$d_4$)
δ = 4.90 (br s, 1H), 4.23 - 4.07 (m, 2H), 3.92 (dd, $J$ = 2.2, 12.0 Hz, 1H), 3.66 (dd, $J$ = 6.6, 11.9 Hz, 1H), 3.44 - 3.36 (m, 2H), 3.29 - 3.11 (m, 4H)

*Synthesis of 2-acetamido-3-(carboxymethylsulfanyl)propanoic acid.*

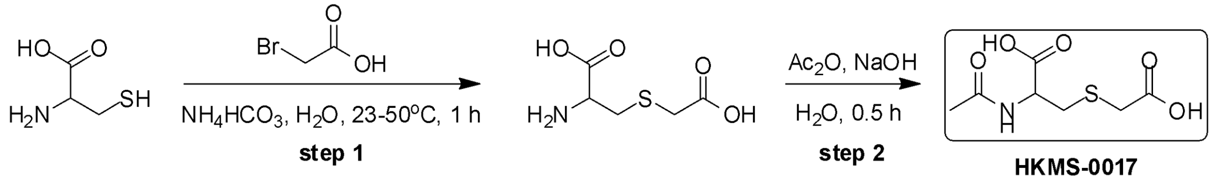

**HKMS-0017**

Step 1: synthesis of 2-amino-3-(carboxymethylsulfanyl)propanoic acid

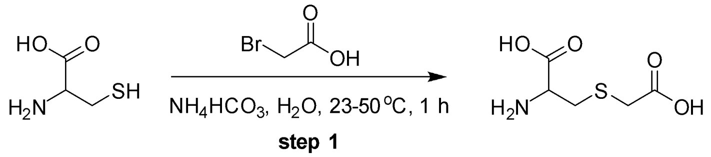


2-amino-3-sulfanyl-propanoic acid (1 g, 8.25 mmol, 1 eq) and 2-bromoacetic acid (688.10 mg, 4.95 mmol, 355.79 µL, 0.6 eq) was dissolved in $H_2O$ (4 mL) at 23°C. To the solution was added $NH_4HCO_3$ (1.30 g, 16.51 mmol, 2 eq). Then the mixture was adjusted to pH = 7.5 using $NH_4OH$, while the reaction mixture temperature was maintained at about 45-50°C. The reaction mixture was stirred at 50°C for 1 hr. After the reaction was completed, pH of the reaction mixture was adjusted to pH = 2.8 with hydrochloric acid. The whole regulation process is controlled at about 1 minute, cooling to give 2-amino-3-(carboxymethylsulfanyl)propanoic acid (440 mg, 2.46 mmol, 29.75% yield) as a white solid.
**$^1$H NMR:** (400 MHz, DMSO-$d_6$)
δ = 3.48 (br dd, $J$ = 4.0, 7.9 Hz, 5H), 3.27 (br d, $J$ = 2.6 Hz, 2H), 3.05 (br dd, $J$ = 4.0, 14.4 Hz, 1H), 2.87 (br dd, $J$ = 7.9, 14.5 Hz, 1H)

Step 2: synthesis of 2-acetamido-3-(carboxymethylsulfanyl)propanoic acid

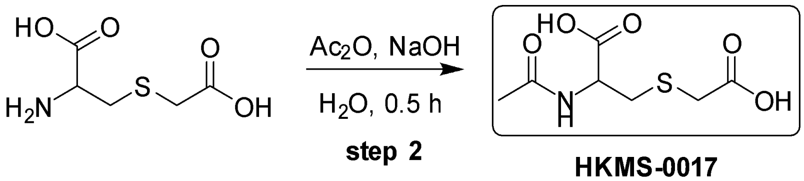


**HKMS-0017**

2-amino-3-(carboxymethylsulfanyl)propanoic acid (0.2 g, 1.12 mmol, 1 eq) was dissolved in $H_2O$ (1.3 mL) at 0°C. The pH was adjusted to 7 with 4 N NaOH at 0°C. $Ac_2O$ (227.88 mg, 2.23 mmol, 209.65 μL, 2 eq) was added slowly dropwise together with 4N NaOH (in order to maintain pH = 7). The mixture was stirred at 25°C at 0.5 h. To the reaction mixture was added Amberlite (R) IRC120H (100 mg) and then the mixture was stirred for 30 min. The suspension was filtered. The filtrate was purified directly. The filtrate was purified by prep-HPLC ($HCOONH_4$ condition, column: HUAPU 1010-X Amide 100 × 30 mm × 10 μm; mobile phase: [H2O(10mM $HCOONH_4$)-ACN]; gradient:90%-60% B over 15.0 min) to give 2-acetamido-3-(carboxymethylsulfanyl)propanoic acid (32.8 mg, 139.54 μmol, 12.50% yield, 94.12% purity) as colorless oil, which was confirmed by LCMS and $^1$H NMR.

**LCMS:** (M+1):222.11@ 4.824 min (90_50CD, 10 min)
**$^1$H NMR:** (400 MHz, DMSO-d6)
δ = 8.43 (s, 1H), 7.89 (br d, *J* = 7.1 Hz, 1H), 4.15 - 4.05 (m, 2H), 3.02 (s, 2H), 2.85 - 2.70 (m, 2H), 1.82 (s, 3H)
**$^{2D}$H NMR:** HSQC, COSY, HMBC conforms to desired structure.

*Synthesis of 3-(carboxymethylsulfanyl)-2-hydroxy-propanoic acid.*

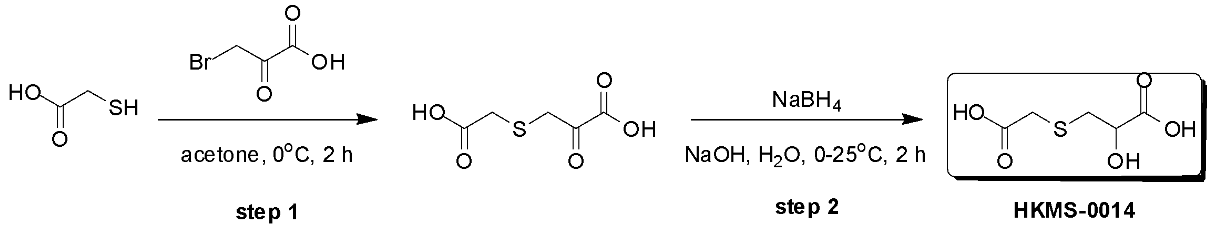


**HKMS-0014**

Step 1: synthesis of 3-(carboxymethylsulfanyl)-2-oxo-propanoic acid

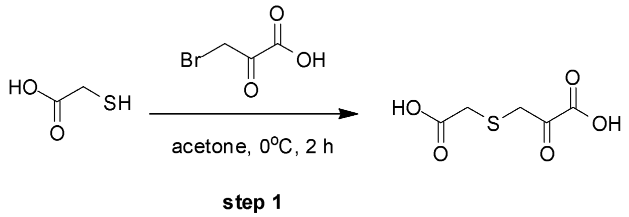


To a solution of 3-bromo-2-oxo-propanoic acid (380.45 mg, 2.28 mmol, 1.3 *eq*) in acetone (5 mL) was added carboxymethylsulfanylsodium (200 mg, 1.75 mmol, 1 *eq*) at 0°C. The mixture was stirred at 0°C for 2 h. The mixture was filtered off and filtrate was concentrated to give 3-(carboxymethylsulfanyl)-2-oxo-propanoic acid (320 mg, crude) as a white solid. The crude product was used for the next step without further purification.

**$^1$H NMR:** (400 MHz, acetonitrile-d3)
δ = 3.78 (s, 2H), 3.26 (s, 2H)

Step 2: synthesis of 3-(carboxymethylsulfanyl)-2-hydroxy-propanoic acid

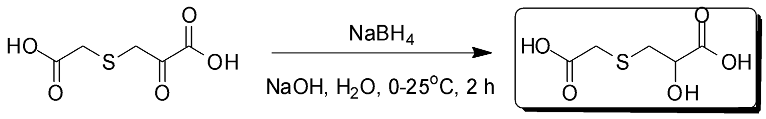


**step 2** **HKMS-0014**

A solution of 3-(carboxymethylsulfanyl)-2-oxo-propanoic acid (220 mg, 1.23 mmol, 1 *eq*) in $H_2O$ (2.5 mL) was adjusted to pH = 10 with NaOH (2 M, 1.23 mL, 2 *eq*), and then treated portion-wise with $NaBH_4$ (46.72 mg, 1.23 mmol, 1 *eq*). The mixture was stirred at 25°C for 2 h. The mixture was poured into 2M HCl (0.5 mL) to pH = 6 at 0°C. The residue was purified by prep-HPLC (TFA condition, column: Kromasil C18(W) 150 × 30 × 10; mobile phase: [$H_2O$(0.1%TFA)-ACN]; gradient:1%-30% B over 12.0 min) and (HCl condition, column: Kromasil C18(W) 150 × 30 × 10; mobile phase: [$H_2O$ (0.04%HCl)-ACN]; gradient:1%-10% B over 10.0 min) to give 3-(carboxymethylsulfanyl)-2-hydroxy-propanoic acid (18 mg, 99.20% purity, 99.10 µmol, 4.56% yield over 2 steps) as colorless oil.

**LCMS:** (M+23):203.0 @ 5.037min (T3-20MIN-220-254-ELSD, 20 min)
**$^1$H NMR:** (400 MHz, Deuterium Oxide)
δ = 4.49 (dd, *J* = 4.0, 6.6 Hz, 1H), 3.51 - 3.38 (m, 2H), 3.18 - 3.07 (m, 1H), 3.03 - 2.88 (m, 1H)
**13C-NMR HSQC HMBC** Conforms to the desired structure.
